# Orchids, hawkmoths and Darwin revisited: How adaptation to pollination by hawk-moths spurred the speciation of Old and New World angraecoids

**DOI:** 10.1101/2024.12.06.627257

**Authors:** João Farminhão, Géromine Collobert, Benoît Perez-Lamarque, Simon Verlynde, Brigitte Ramandimbisoa, Esra Kaymak, Laura Azandi, Vincent Droissart, Murielle Simo-Droissart, Steve Johnson, Florent Martos, Tariq Stévart

## Abstract

Specialised pollination has often been proposed as a major driver of the unrivalled diversification of angiosperms. With its unparalleled variation in flower depth, ranging from spurless to 40 cm spurred flowers pollinated by hawkmoths, angraecoid orchids (Angraecinae) provide unique opportunities to reveal the impact of floral specialisation on diversification rates in the Afrotropics and Neotropics. We compiled floral characters for 327 angraecoid species, and assigned these species to newly formalised floral syndrome categories calibrated against pollination case studies. We then estimated ancestral character states and state-dependent speciation rates using the latest time-calibrated phylogeny of angraecoids. We found that white flowers consistent with micro-sphingophily (2 cm≥ spurs <8.7 cm), macro-sphingophily (8.7 cm≥ spurs <18.6 cm) and mega-sphingophily (spurs ≥18.6 cm) evolved repeatedly in angraecoids, and are ancestral in some non-sphingophilous clades. Reversals to non-sphingophily and pigmented flowers suggest higher evolutionary lability of these traits than what is traditionally thought. Increasing floral specialisation with long spurs does not seem to enhance angraecoid orchids’ speciation rates, since micro-sphingophily appears to be associated with the highest speciation rates. An abundance of short-proboscid hawkmoths in Madagascar, as an ecological opportunity, may have accelerated the speciation of micro-sphingophilous taxa.

## INTRODUCTION

Since Darwin [1], the apparent correlation between species richness and specialised pollination has often led to the assumption that the latter is a major driver of the unrivalled diversification of angiosperms, either by increasing speciation rates and/or decreasing extinction rates [2–4]. However, the global patterns of species richness and diversification rates are seemingly unlinked in plants [5]. Instead, the role of specialised pollination in enhancing speciation rates may depend on the system [6]. Indeed, it has been reported that despite specialised pollination, a significant pollen flow among sympatric species tends to prevent the complete reproductive isolation conducive to speciation [3]. Orchidaceae may be a notable exception in this respect, because of their agglutinated pollen grains named pollinia, which are often precisely placed on the pollinator so that it comes into contact only with the stigma of conspecifics [3]. Accordingly, pollen attachment site shifts are surmised to be important in orchids and other plants exhibiting high floral integration [3,7,8]. In Orchidaceae, both the evolution of pollinia and specialised pollination by lepidopterans are associated with increased speciation and overall diversification rates [9]. Pollination by lepidopterans, and especially hawkmoths (Sphingidae), often involves a wide range of nectary spur lengths that are only accessible to pollinators with sufficiently long mouthparts [4,9,10]. The longer the spur, the lower the number of visitor species [10], a measure of ecological specialisation [4].

Angraecoid orchids (subtribe Angraecinae) — consisting of approximately 800 species distributed in Africa, the Southwestern Indian Ocean islands and the Neotropics — present the largest nectar spurs of all angiosperms [11,12]. Hawkmoth pollination in long-spurred angraecoids was first suggested by Darwin [13] in *Angraecum sesquipedale* Thouars, and later investigated in several Malagasy [14–18] and continental African taxa [19–22], and more recently in Neotropical species [23–25]. Furthermore, angraecoid species vary in flower resupination [26], which translates into pollen transfer from below to the ventral side (i.e., sternotribic pollination) or from above to the dorsal side (i.e., nototribic pollination) of the pollinator [14], potentially leading to pre-zygotic isolation. However, the impact of hawkmoth pollination and shifts in the orientation of pollen attachment sites on the diversification of angraecoids remains to be investigated.

The study of the macroevolutionary impact of specialised hawkmoth pollination is contingent upon the inventory of known pollination systems within a lineage. In light of the incomplete nature of the pollination record, the recognition of floral syndromes — trait combinations required for occupying particular pollinator ecological niches — represents a crucial step in completing this task [27]. However, these syndromes must first be calibrated against actual pollination case studies to be considered reliable [28]. Sphingophily, involving pollination by hawkmoths, is one of the most easily recognisable syndromes, with a set of traits including long nectar tubes and predominantly white nocturnally fragrant flowers [10,29,30]. There are distinct sub-syndromes among hawkmoth-pollinated plants that are characterised by differences in flower depth, and these reflect adaptations to various functional groups (guilds) of hawkmoths that differ in proboscis length. Depending on the biogeographical region, there can be either two [10,31] or three [32] hawkmoth guilds [10,28].

In view of their unparalleled spur length disparity, high floral integration, and high species richness, angraecoid orchids represent an ideal system to examine the impacts of sphingophily on speciation. In this study, we sought to (1) identify sub-syndromes of sphingophily reflecting different levels of ecological specialisation, by calibrating the evolutionary correlation between orchid spur and hawkmoth proboscis lengths to reliably assign angraecoid species to hawkmoth pollination guilds; (2) reconstruct the evolutionary history of flower traits associated with sphingophily, including resupination, in angraecoids; and (3) examine the impact of sphingophily and shifts in the orientation of pollen attachment sites on angraecoid speciation rates. We hypothesise that the longer the spur, the more specialised the pollination and consequently the higher the speciation rates. In addition, since tongue length in sphingids is positively correlated with body size [33], we expect that long-proboscid hawkmoths provide more sites for pollen attachment than short-proboscid sphingids, thus increasing the available pollination niches for angraecoid species and, consequently, the speciation within the same hawkmoth guild.

## MATERIALS AND METHODS

### Survey of floral traits and syndromes in angraecoids

Mean spur length, flower colour, and resupination of 327 angraecoid species were collated from floristic and monographic treatments, species protologues (see full list of references in electronic supplementary material), and complemented with photographs from databases and direct measurements of spur length on herbarium specimens housed at BRLU, K, P and WAG (acronyms following *Index Herbariorum* [34]). Geographical ranges of species were retrieved from the World Checklist of Selected Plant Families (now the Plants of the World Online database) [35] and follow the TDWG geographical scheme [36]. The frequency of mean spur length, discretised into 1 cm intervals, was calculated for all angraecoids, and then separately for species from the Southwest Indian Ocean (SWIO) islands, continental Africa, and the Neotropics (electronic supplementary material, figure S1a). Flower colour was binarily classified as ‘white/partially white’ or ‘fully pigmented’. Flowers were classified as either resupinate, when presenting a lowermost lip, or hyper-resupinate, when presenting an uppermost lip.

To recognise sphingophilous flowers among the angraecoids, we classified species according to their flower colour first, and then to spur length. Species with fully pigmented flowers were scored as non-sphingophilous, regardless of their spur length, as were white/partially white flowers with a spur <2 cm (this includes examples of butterfly, settling moth, orthoptera or bird pollinations) [37–41]. White/partially white flowers with a spur ≥2 cm were scored as sphingophilous. Although a few of these long-spurred white angraecoids are actually autogamous (viz. *Aerangis punctata*, *Angraecum borbonicum*, *A. cornigerum*, *Jumellea divaricata*, *J. exilis*, *J. recurva*, *J. stenophylla*), their impact on the estimation of ancestral states and speciation rates should be limited, since they likely correspond to recent, isolated events of colonisation of oceanic islands where their pollinators were missing [42], whereas creating a special category for them may lead to over-parameterisation of our models. To identify sub-syndromes reflecting pollination of angraecoid species by hawkmoth functional groups, we comprehensively surveyed the case studies reporting visits of angraecoid orchids by hawkmoths (31 observations, 19 species), with and without pollinia removal (20 and 11 observations, respectively) [15–20,22,24,25,31]. We recorded a further 56 observations of hawkmoth pollination in 19 species of non-angraecoid orchids, including 42 observations of pollinia removal by the hawkmoth visitor and 14 visits without pollinia removal [32,43–59]. For each observation, the mean length of the flower spur was plotted against the mean length of the proboscis of the moths observed. We evaluated the correlation between spur length of specimens with confirmed pollinia removal and the corresponding pollinator tongue lengths using a linear model (figure 1a). In addition, we evaluated the relationship between mean spur length and depth to the nectar meniscus for 12 of the orchid species surveyed in the pollination case studies. Based on previous studies, the surveyed hawkmoth visitors were divided into functional groups of short-proboscid (mostly Macroglossinae), long-proboscid (e.g. *Agrius convolvuli*, *Coelonia fulvinotata, Panogena lingens*) and ultra-long proboscid hawk-moths (viz. *Xanthopan morganii*, *X. praedicta* and *Coelonia solani*) [10,31,32]. The intersection between the regression line and the largest mean proboscis length in each hawkmoth guild was established as the spur length threshold delimiting our sub-syndromes (Fig. 1a, dotted lines). Based on our correlation analysis, we identified as (1) micro-sphingophilous the species with a mean spur length ≥2 cm and <8.7 cm, most likely to be pollinated by short-proboscid hawkmoths; (2) macro-sphingophilous the species with a mean spur length ≥8.7 cm and <18.6 cm, most likely to be pollinated by long-proboscid hawkmoths; or (iii) mega-sphingophilous the species with a mean spur length ≥18.6 cm, most likely pollinated by ultra-long proboscid hawkmoths (figure 1b–e). All floral traits and syndromes are summarised in table 1.

**Figure 1.**
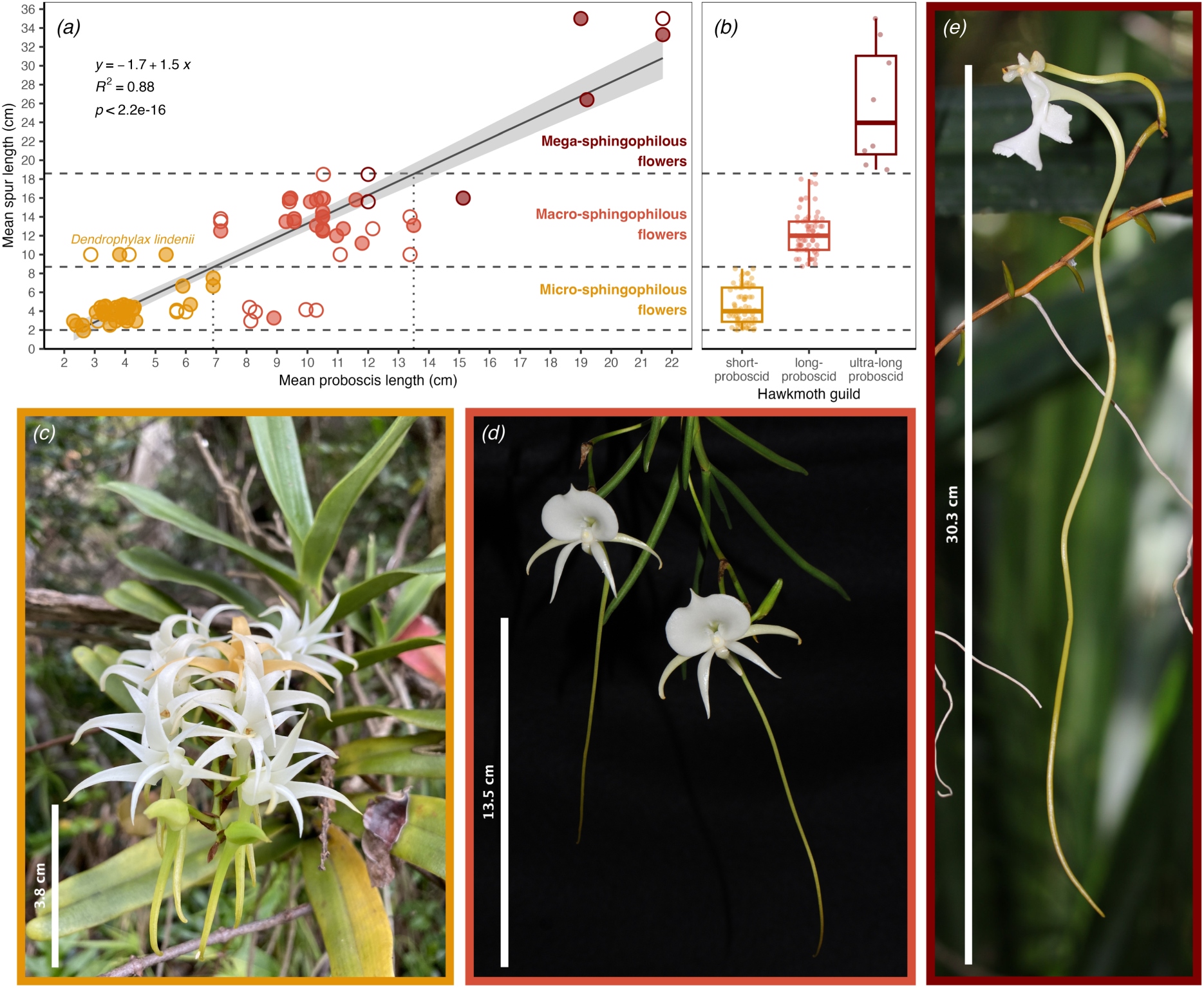
Formalisation of sub-syndromes of sphingophily. *(a)* Linear regression (black line) with confidence interval (in grey) between mean hawkmoth proboscis and orchid spur lengths in pollination case studies; filled circles: observed pollinium removal; empty circles: visit without pollinium removal; colours correspond to hawkmoth guilds (see methods). *(b)* Assignation of thresholds from *(a)* to 327 angraecoid species reflecting pollination by the different hawkmoth guilds. *(c)* The micro-sphingophilous, resupinate flowers of *Cyrtorchis arcuata*, note the orange-coloured senescent flower; photograph by Simon Atwood. *(d)* The macro-sphingophilous, hyper-resupinate flowers of *Angraecum scottianum*; photograph by Leo Klemm. *(e)* The mega-sphingophilous, resupinate flower of *Solenangis impraedicta*; photograph by Marie Savignac.

**Table 1:** Definition of floral syndromes in Angraecinae. White flowers tinged at tips, or flowers for which only the lip is white, are grouped together as partially white flowers. Spur length are mean values of species. Resupinate flowers result from an 180° twist, hyper-resupinate from a 360° twist of the ovary and pedicel.

| Floral syndrome | Spur length | Flower colour | Resupination |
| --- | --- | --- | --- |
| non-sphingophilous | $\geq 0$ cm | fully pigmented | resupinate |
|  |  |  | hyper-resupinate |
| | $< 2$ cm | white/partially white | resupinate |
|  |  |  | hyper-resupinate |
| micro-sphingophilous | $\geq 2$ cm to $< 8.7$ cm | white/partially white | resupinate |
|  |  |  | hyper-resupinate |
| macro-sphingophilous | $\geq 8.7$ cm to $< 18.6$ cm | white/partially white | resupinate |
|  |  |  | hyper-resupinate |
| mega-sphingophilous | $\geq 18.6$ cm | white/partially white | resupinate |
|  |  |  | hyper-resupinate |

### Time-calibrated phylogenetic and ancestral range inferences

We produced a molecular dataset for 327 angraecoid species (42% of species diversity of the group) and five outgroups in Aeridinae, for a total of 332 taxa, 1709 sequences (of which 137 are new compared to the previous phylogenetic analysis of angraecoids [60]) and 7,451 characters (electronic supplementary material, table S1). Six markers, i.e. nuclear ITS-1 (309 sequences), plastid *matK–trnK* (323), *rps16* (291), *trnC–petN* (199), *trnL–trnF* (308) and *ycf1* (279), were aligned with MUSCLE in Geneious 9.1.7 (Biomatters, New Zealand), manually edited, and concatenated. The best-fitting partition scheme was selected using ModelFinder in IQ-TREE 1.6.12 [61]. Time-calibrated tree inference from the molecular dataset was performed using a parallel tempering approach implemented in CoupledMCMC template in BEAST 2.6.3 [62], under a relaxed uncorrelated log-normal molecular clock, and the birth-death process as tree prior. A diffuse gamma distribution (α = 0.001 and β = 1000) was applied to both clock mean and birth rate priors. Other priors were left as default. Site model averaging using bModelTest 1.2.1 was performed, as recommended by Bouckaert and Drummond [63], with mutation rate estimated and fixed mean substitution rate. Due to the few known orchid fossils and none within angraecoids, a secondary calibration point relying on the stem age of Angraecinae estimated by Givnish *et al.* [9] was chosen, allowing estimation of the clock rate. The divergence time between Angraecinae and Aeridinae was given a normal distribution with a mean set to 21.21 My and a standard deviation to 3.14, therefore encompassing the 95% Highest Posterior Density (HPD) interval found by Givnish *et al.* [9]. We performed four independent runs of 50 million steps and 6 chains each, resampling every 5,000 steps. Stationarity and convergence of runs were assessed using Tracer 1.7.1 [64]. After discarding the first 10% as burn-in, trees were combined using LogCombiner 2.6.3. A maximum clade credibility (MCC) tree with median node heights was generated using TreeAnnotator 2.6.3 (electronic supplementary material, figure S2). Its topological consistency was evaluated in comparison with phylograms calculated using maximum likelihood (ML) in RAxML-HPC2 under GTRGAMMA [65], and Bayesian inference (BI) in MrBayes 3.2.7 [66], with model parameters independently estimated for each partition.

Because transitions between pollinator guilds can be correlated with changes in biogeographic range due to the different composition of pollinator communities, we estimated from the time-calibrated MCC tree the ancestral areas of angraecoids (electronic supplementary material, figure S3) using BioGeoBEARS 1.1.3 [67] in R 4.4.0 [68]. The best-fitting model was selected by AICc comparison (electronic supplementary material, table S2).

### Ancestral state estimation and trait-dependent speciation rates

The evolution of discrete traits was jointly modelled with species diversification using state-dependent speciation and extinction (SSE) models implemented in the R package hisse 2.1.11 [69,70]. Because of identifiability concerns about extinction rates and extinction fractions [71–73], we fixed the extinction fraction to be constant across states, and focused only on the state-dependent speciation rates. We ran a constant-rate (CR) birth-death model; an examined trait-dependent model (ETD) corresponding to a BiSSE/MuSSE model; a CID-2, CID-4 or CID-8 models (trait-independent speciation with two, four and eight hidden states, respectively, equalling the parameter number of trait-dependent models); and an examined and concealed state-dependent model (ECTD) with 2 hidden states, corresponding to a HiSSE/MuHiSSE model. Hidden states allow for heterogeneity in transition rates across different parts of a phylogeny, where hidden (unmeasured) factors may have either promoted or constrained diversification, in concert with the measured character [72]. The transition matrix for the four-state floral syndrome (non-/micro-/macro-/mega-sphingophilous) was ranked i.e., only transitions to states directly under or above were allowed. However, direct transitions from any of the three sphingophilous states to non-sphingophily were also allowed, because a homeotic mutation could lead to the complete loss of the spur, as observed in Aeridinae [74]. Confidence intervals of the parameters were estimated by adaptively sampling 1,000 points of the likelihood surface of the Maximum Likelihood Estimates (MLE) using the functions hisse:: *SupportRegionHiSSE* and hisse:: *SupportRegionMuHiSSE*. Marginal reconstruction of states and rates was performed using functions hisse::*MarginReconHiSSE* and hisse::*MarginReconMuHiSSE* for all models with a δAICc <2 relative to the model minim-ising the AICc. Ancestral states and net speciation rates at tips were then model averaged in proportion to AICc weights using hisse::*ConvertManyToMultiState* and hisse::*GetModelAveRates*, respectively.

## RESULTS

### Survey of floral traits and floral syndromes in angraecoids

Surveyed mean spur lengths vary from zero in four species (e.g. *Angraecum corrugatum*) to 35 cm in *A. sesquipedale*. Short spurs (<2 cm) predominate in all regions (55% of all sampled species). The frequency of spur length tends to follow an exponential distribution decreasing towards long spurs, except in the SWIO. This latter region stands out with a pronounced peak of spur length centred around a mode at 11–12 cm accounting for 9.4% of species, and mean values surpassing 26 cm in four species (*Angraecum longicalcar*, *A. sesquipedale*, *A. sororium* and *Solenangis impraedicta*) (electronic supplementary material, figures S1a, S2). White or partially white flowers predominate among sampled angraecoids (71%), with regional variations: 82% in the SWIO, 66% in continental Africa and 47% in America. Accordingly, sphingophilous floral syndromes are represented in 62% of species in the SWIO (45% in all sampled species), against only 36% and 21% in continental Africa and America respectively. Within sphingophilous flowers, micro-sphingophily is the most common syndrome in continental Africa (27% of all floral syndromes, 23% in all sampled species), whereas macro-sphingophily predominate in the SWIO (39%, against 20% in all sampled species). Mega-sphingophily account for 2–3% of species, except in America where this syndrome is absent (electronic supplementary material, figure S1b). Distribution of spur lengths within sub-syndromes of sphingophily are shown with the median, first and third quartiles, and whiskers on figure 1b. Resupinate flowers are more common (84%) than hyper-resupinate flowers (16%), with no major variation across regions and floral syndromes.

### Ancestral state estimates

The MRCA of Angraecinae was confidently inferred to have had resupinate (P = 0.96), entirely pigmented flowers (P = 0.94), indicating that a non-sphingophilous floral syndrome is most likely ancestral, although this state was found to be unlikely (P = 0.2) in the explicit estimation of the ancestral syndromes (figure 2, electronic supplementary material, figures S4–S5). The very high transition rates between macro-sphingophily and mega-sphingophily, and to a lesser extent micro-sphingophily, may indicate that these states are somewhat highly labile on evolutionary timescales. Consequently, close-relatives are as likely to share the same sub-syndrome of sphingophily as to have different ones (electronic supplementary material, figure S6a). Indeed, the evolution of syndromes appears to have been highly dynamic, resulting in high uncertainty at several nodes and multiple reversions to non-sphingophily (figure 2). Nevertheless, mega-sphingophily has evolved at least six times independently, in the MRCA of *Angraecum sesquipedale* and *A. sororium* (P = 0.98; PP = 1), in the MRCA of *Aerangis gracillima* and *A. stelligera* (P = 0.98; PP = 1), and at tips in *Angraecum longicalcar*, *Aerangis bouarensis*, *Solenangis impraedicta*, and *Plectrelminthus caudatus*. In *S. impraedicta*, the transition to mega-sphingophily is concomitant with a migration event from continental Africa to the SWIO (electronic supplementary material, figure S3). No other particular correlation to biogeographic history was observed. Interestingly, transition rates also reveal that some lineages may have few to no capacity to revert to shorter-spurred flowers once long-spurred sphingophilous flowers have evolved (rate category B; electronic supplementary material, figure S6a).

**Figure 2.**
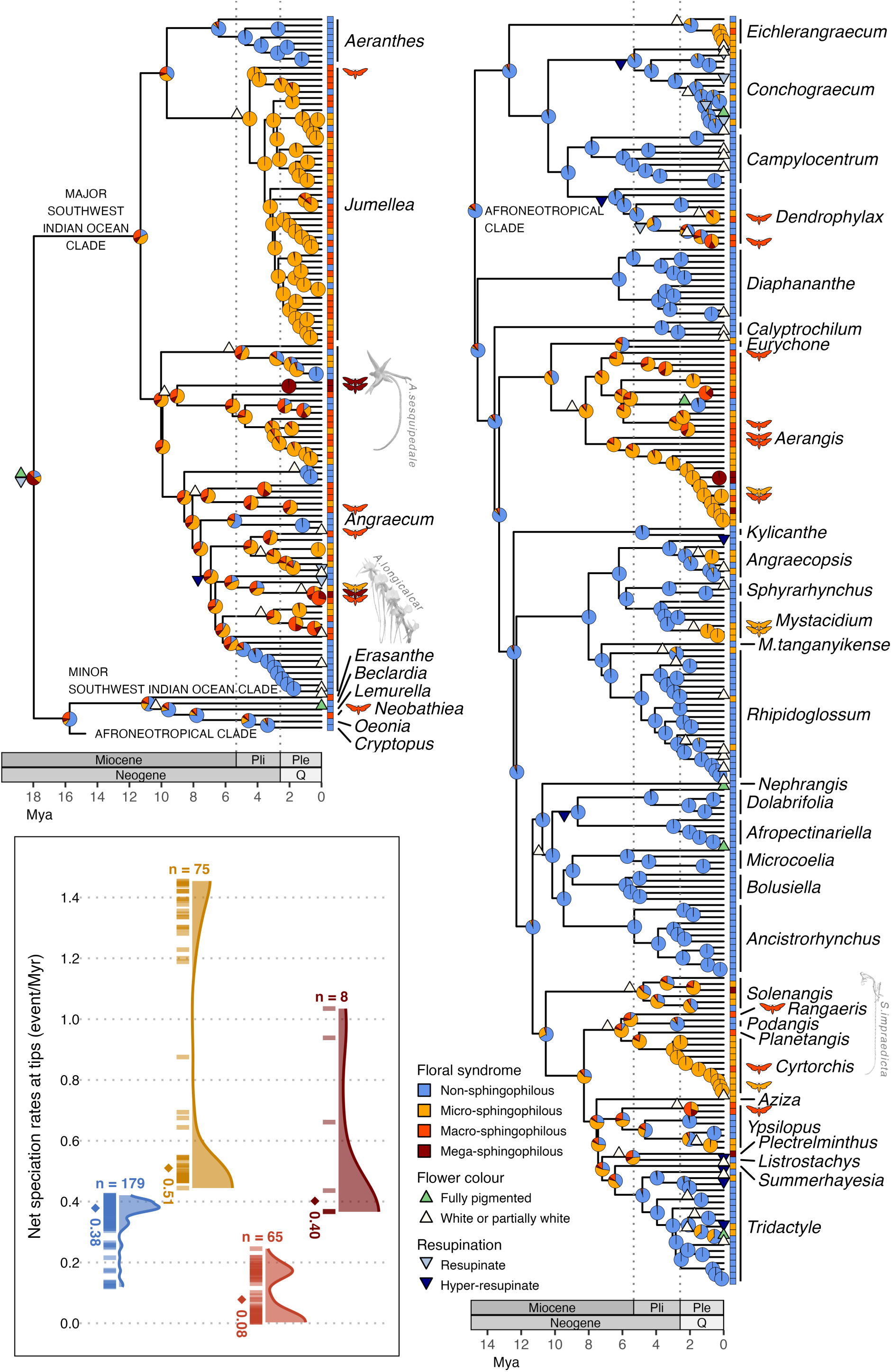
Ancestral state estimation of floral syndromes using hidden-state SSE models on the dated phylogeny of 327 species of angraecoid orchids; pie charts at nodes represent ancestral syndromes and their probabilities under an ECTD model. For flower colour and resupination, only significant state changes, i.e. state probability >0.5 while <0.5 in the ancestral node, are indicated by coloured triangles (see electronic supplementary material, figures S4–S5 for full ancestral state estimations of these traits). The character states of extant species are represented on the right side of the tree by coloured boxes. In addition, species for which a sphinx has been observed visiting or carrying pollinaria are indicated by a sphinx silhouette. Pli: Pliocene; Ple: Pleistocene; Q: Quaternary. The inset shows the net speciation rates at tips associated with each state with density plots; ‘n’ indicates the number of species per state; the median is indicated by a diamond with value below.

The transition rate from pigmented to white flowers was five times higher than the reverse transition rate (electronic supplementary material, figure S6b). Interestingly, while reversion to pigmented flowers forces transitions to non-sphingophily, the opposite is seldom observed, suggesting that transitions to non-sphingophily were mainly due to spur shortening rather than reversion to pigmented flowers (figure 2). Hyper-resupinate flowers likely appeared at least nine times and reverted to a resupinate state seven times (electronic supplementary material, figure S5), with no apparent association with transitions in spur length, flower colour, floral syndromes, or biogeographic history.

### State-dependent speciation rates

For flower colour and resupination, the CID model had the lowest AICc, indicating that speciation rates are likely independent of the state of these traits (electronic supplementary material, tables S3–S4). For syndromes, the ECTD model was selected (electronic supplementary material, table S5), indicating that at least one syndrome is significantly implicated in driving the speciation rates, but that it does not act in isolation, meaning there is at least another unmeasured trait that conjointly drives the rates, and that only their interaction can explain the speciation pattern. Indeed, while the highest rates (max = 1.45 event/Myr) were associated with the micro-sphingophilous syndrome, not all species with this syndrome had high rates, lowering the median rate of micro-sphingophilous flowers to 0.51 (figure 2). Although high-rate micro-sphingophilous flowers were found in both continental Africa and in the SWIO, the latter region hosted a higher proportion of the high-rate micro-sphingophilous flowers relative to the total number of micro-sphingophilous species in that region, raising the median to 1.03 (electronic supplementary material, figure S7). Macro-sphingophilous flowers tended to have the lowest rates of all syndromes (median = 0.08) in all regions.

## DISCUSSION

### Calibrating the sub-syndromes of sphingophily

In this study, we propose a calibration of the sub-syndromes of sphingophily based on the proboscis lengths of hawkmoth guilds and on the spur length of the flowers they pollinate. Our aim was to establish the most objective — but not absolute — thresholds possible for distinguishing sub-syndromes of sphingophily in angraecoid orchids. They may not, and were not intended to, be generalised to other groups. These thresholds are obviously dependent on the record of visits and pollinia removal by hawkmoths and may need to be adjusted in the future when new observations are made.

Although fairly linear, the relationship between proboscis length and spur length is not strict. Indeed, pollinia can be placed at various places along the proboscis or even the head of a pollinator, enabling species with different spur lengths to share the same pollinator [15]. On this note, the evolution and the impact on diversification of the position of pollinia, rather than of their orientation (resulting from the flower resupination) on the pollinator, may prove more significant. In addition, rather than total spur length, proboscis length of pollinators matches the depth to nectar in the spur (i.e. the total spur length minus the height of the nectar column). This explains why short-tongued hawkmoths, with a proboscis no longer than 6 cm, were observed pollinating the macro-sphingophilous *Dendrophylax lindenii*, whose spur ranges from 9 to 17 cm in length [23,24] (figure 1a). Nevertheless, hawkmoths with longer proboscides are expected to have driven the elongation of the spur of this species, hence its classification as macro-sphingophilous. Moreover, the relationship between the mean spur length and the depth to the nectar meniscus of 12 orchid species is fairly linear (R2 = 0.8, p-value = 8.1e-05; electronic supplementary material, figure S8), suggesting that spur length is indeed a good predictor of pollination guild. Nevertheless, other characters, such as nectar composition and floral scent chemistry, can help refine the prediction of the pollinator type [11,75]. For example, autogamous species, despite having typical sphingophilous flowers, lack scent [42,76]. Transitions to autogamy were probably caused by dispersal events to remote oceanic islands where suitable pollinators were missing [42]. Similarly, the biogeographically distinct lineages with very few dispersal events in angraecoids (electronic supplementary material, figure S2) may partly reflect limits to dispersal in conjunction with missing pollinators and/or mycorrhizal fungal symbionts [77,78].

### The evolution of flower colour

Our result that transitions from pigmented to white flowers were five times more frequent than the reverse transitions is consistent with previous findings in other groups [79]. Indeed, the evolution of white flowers typically results from loss-of-function (LOF) mutations, which are estimated to be difficultly reversible [79]. However, in white-flowered angraecoids, senescent flowers often become pigmented (see figure 1c) due to the likely presence of carotenoids, suggesting that LOF mutations have not fully affected floral pigment function. Thus, gene regulation of carotenoids may better account for observed colour variation in angraecoids. Indeed Zhang *et al*. [80] found that white individuals of the polymorphic orchid *Pleione limprichtii* were produced by differentially-expressed genes, a mechanism also at the origin of white varieties in *Primula* [81]. On the other hand, a loss of anthocyanin synthesis may be synapomorphic in angraecoids, as evidenced by the absence of pink-flowered angraecoids in contrast to sister Aeridinae, where anthocyanin-rich floral palettes abound.

### Micro-sphingophily spurred the speciation of angraecoids

Sphingophilous flowers can be regarded as highly specialised [4], which is suggested to increase speciation rates, by facilitating reproductive isolation in sympatry and reinforcement upon secondary contact [3] (but see below). However, contrary to our primary assumption, we found that micro-sphingophilous angraecoids were associated with higher speciation rates than macro-sphingophilous and mega-sphingophilous flowers (figure 2). Micro-sphingophilous angraecoids are indeed less ecologically specialised overall than the other sub-syndromes, as they are statistically visited by a higher number of hawkmoth visitor species (a mea sure of ecological specialisation) than longer-spurred species, although micro-sphingophilous flowers can also specialise in pollination by a few pollinators [10]. A possible explanation is that low speciation rates found in macro-sphingophilous and mega-sphingophilous species could result from a reduced number of available niches for these hyper-specialised flowers [82]. Indeed, the availability of a pollination niche is determined not only by the presence of a potential pollinator species, but also by its abundance, and in hawkmoths, longer proboscis lengths are correlated with lower abundance [10]. Therefore, more species-rich and individually abundant short-proboscid hawkmoths may have triggered the speciation of angraecoids within this guild more than less abundant guilds of hawkmoth with longer proboscides. Moreover, short-proboscid hawkmoths (with a proboscis length <7 cm, see figure 1) seem to be proportionally more abundant in some regions than in others, notably in Madagascar or the Kenyan drylands where they account for around three-quarters of the light-trapped hawkmoth individuals, respectively, against around half of the individuals in subtropical South Africa [10]. Such differences in the composition of hawkmoth communities may additionally explain the bimodal distribution of speciation rates within micro-sphingophilous flowers. Notably, the prevalence of high speciation rates within micro-sphingophilous flowers from the SWIO might be related to a high abundance and/or diversity of short-proboscid hawkmoth species in this region. However, to test this hypothesis, more precise data on hawkmoth abundances and diversity in the distribution range of each angraecoid species would be needed.

In addition to promoting speciation rates, specialised pollination has been hypothesised to decrease extinction rates, because hawkmoths are highly vagile insects potentially ensuring connectivity among small, fragmented populations and thus limiting local extinction [83,84]. However, diversification models often critically underestimate extinction rates [85], which is why we have chosen not to estimate them. The impact of floral specialisation on extinction rates could be tested in a Bayesian framework by incorporating a prior distribution that enforces high extinction rates as in Zenil-Ferguson *et al*. [82].

Lower speciation rates in more specialised flowers could also be indicative that hyper-specialisation is an evolutionary dead-end. This alternative hypothesis, implying low diversification rates and difficult reversals to less specialised systems, has hitherto received mixed evidence depending on the study system (e.g. [3,82,86–89]). Transition rates indicate that floral specialisation, notably on long and ultra-long proboscid sphingids, may indeed be almost irreversible, but only in some lineages of angraecoids, while conversely in other lineages reversions to less specialised pollination systems seem to have been relatively frequent (electronic supplementary material, figure S6). Therefore, it is possible that at least two evolution-ary pathways coexist in angraecoids: lineages in which floral specialisation lacks evolutionary lability and is an evolutionary dead-end, and lineages in which specialisation is labile and may have no or positive impact on diversification rates. This hypothesis also needs further investigation. Nevertheless, floral specialisation, as important as it may be, is unlikely to have influenced the speciation rates in isolation [72]. Other factors could have simultaneously influenced speciation rates, e.g. shifts in chromosome number [60], evolution of C_3_/CAM photosynthesis [9], or environmental fluctuations such as rainforest regression [90], changes in monsoon climates or orogenesis [91]. Notably, cycles of forest expansion and contraction associated with climatic changes during the Plio-Pleistocene could have increased allopatric speciation rates in forest-adapted lineages [90]. Finally, assembling phylogenetic datasets for other hawkmoth-pollinated lineages (*e.g. Adansonia*, *Clerodendrum*, *Ixora*, *Turraea*) and hawkmoths will also enable testing concurring hypotheses related to the effects of sphingophily in speciation and extinction.

## CONCLUSION

Contrary to our expectations, ecological hyper-specialisation of flowers with long spurs does not seem to accelerate speciation in angraecoid orchids. Instead, an abundance of short-proboscid hawkmoths in the SWIO may have constituted an ecological opportunity for angraecoids and enhance the speciation of less specialised, shorter-spurred, micro-sphingophilous taxa. The study of the precise placement of pollinia from different orchid species on a shared pollinator is very promising in this respect. Future research should also concentrate on refining floral syndromes by adding important features for pollinators, e.g. scent and nectar composition, which are largely unknown for most Angraecinae, as well as identifying pollinators in the field. This last point is even more important in the context of biodiversity loss and climate change, which may increase extinction rates for both hawkmoths and angraecoids.

## AUTHOR CONTRIBUTIONS

**João Farminhão:** Conceptualization, Data Curation, Formal analysis, Investigation, Writing– Original Draft. **Géromine Collobert:** Conceptualization, Data Curation, Formal analysis, Investigation, Methodology, Visualisation, Writing–Original Draft. **Benoît Perez-Lamarque:** Methodology, Writing–Review & Editing. **Simon Verlynde:** Data curation, Writing–Review & Editing. **Brigitte Ramandimbisoa:** Investigation, Writing–Review & Editing. **Esra Kaymak:** Investigation, Data Curation, Writing–Review & Editing. **Laura Azandi:** Data curation, Writing–Review & Editing. **Vincent Droissart:** Resources, Writing–Review & Editing. **Murielle Simo-Droissart:** Resources, Writing-Review & Editing. **Steve Johnson:** Methodology, Writing-Review & Editing. **Florent Martos:** Conceptualization, Resources, Supervision, Writing–Review & Editing. **Tariq Stévart:** Funding acquisition, Resources, Supervision, Writing–Review & Editing.

## ACKNOWLEDGEMENTS

The authors thank T. Givnish et D. Spalink for kindly providing the age estimates and 95% HPD of their phylogeny of Orchidaceae. We are also grateful to the American Orchid Society for support of T. Stévart, V. Droissart, M. Simo-Droissart and L. Azandi work in Central Africa. We thank the authorities of the Higher Teachers’ Training College, University of Yaoundé I, for hosting the Yaoundé shade house which produced sample material for DNA extraction. We express our sincere gratitude to Professor Bonaventure Sonké for the supervision of that living collection in Cameroon. Production of new sequences was supported by a grant from the U.S. National Science Foundation (1051547, T. Stévart as PI). Fieldwork in Gabon was undertaken under the Memorandum of Understanding between the Centre National de la Recherche Scientifique et Technologique (CENAREST) and the Missouri Botanical Garden (MBG), permit AR0011/17. We thank the former Director of IPHAMETRA (Institut de Pharmacopée et de Médecine Traditionnelle), Henri Paul Bourobou Bourobou and the former Curator of the Herbier National du Gabon, Nestor Laurier Engone Obiang, for allowing our research. PhD research of J. Farminhão was funded by the Belgian Fund for Research Training in Industry and Agriculture (FRIA) of the F.R.S-FNRS (scholarships F 3/5/5 – FRIA/FC 33848881) and by the Van Buuren-Jaumotte-Demoulin Prize, awarded by the David and Alice Van Buuren Fund. F. Martos is funded by the French national research agency (ANR-19-CE02-0002).

