## Supplementary material for "Orchids, hawkmoths and Darwin revisited: How adaptation to pollination by hawk-moths spurred the speciation of Old and New World angraecoids"

**SUPPLEMENTARY MATERIAL TO 'ORCHIDS, HAWKMOTHS AND DARWIN REVISITED: HOW SPHINGOPHILOUS TRAITS SPURRED THE SPECIATION OF OLD AND NEW WORLD ANGRAECOIDS'**

### Index

Floristic, monographic treatments and species protologues used in the survey of floral traits in angraecoids

1. Danaher MW, Ward C, Zettler LW, Covell CV. 2019 Pollinia Removal and Suspected Pollination of the Endangered Ghost Orchid, *Dendrophylax lindenii* (Orchidaceae) by Various Hawk Moths (Lepidoptera: Sphingidae): Another Mystery Dispelled. *Florida Entomologist* **102**, 671. (doi:10.1653/024.102.0401)
2. Ackerman JD. 2014 *Orchid flora of the Greater Antilles*. Bronx, New York: The New York Botanical Garden Press.
3. Azandi L, Stevart T, Sonké B, Simo-Droissart M, Avana M-L, Droissart V. 2016 Synoptic revision of the genus *Cyrtorchis* Schltr. (Angraecinae, Orchidaceae) in Central Africa, with the description of a new species restricted to submontane vegetation. *Phytotaxa* **267**, 165. (doi:10.11646/phytotaxa.267.3.1)
4. Bogarín D, Pupulin F. 2010 The Genus *Campylocentrum* (Orchidaceae: Angraecinae) in Costa Rica: A Revision. *Harvard Papers in Botany* **15**, 353–414. (doi:10.3100/025.015.0216)
5. Burkhart EP. 2003 Flora of North America North of Mexico. Volume 26. Magnoliophyta: Liliidae: Liliales and Orchidales. *Economic Botany* **57**, 658–659. (doi:10.1663/0013-0001(2003)057[0658:DFABRE]2.0.CO;2)
6. Cribb P, Hermans J. 2009 *Field guide to the Orchids of Madagascar*. Richmond, UK: Kew Pub.
7. Demissew S, Cribb P, Rasmussen FN. 2004 *Field guide to Ethiopian orchids*. Richmond, Surrey, UK: Royal Botanic Gardens, Kew.
8. Droissart V. 2009 Etude taxonomique et biogéographique des plantes endémiques d'Afrique centrale atlantique: le cas des Orchidaceae. Université libre de Bruxelles. See http://hdl.handle.net/2013/.
9. Dunsterville GCK, Garay LA. 1959 *Venezuelan Orchids Illustrated*. London: Andre Deutsch.
10. Feldmann P. 2011 *Orchidées sauvages des Antilles françaises*. Gosier, Guadeloupe: PLB Éditions.
11. Fischer E, Killman D, Delepierre G, Lebel J-P. 2010 *The orchids of Rwanda*. 1. ed. Koblenz: Dep. of Biology, Inst. for Integrated Natural Sciences, Univ. of Koblenz-Landau.
12. Foldats E. 1969 *Flora de Venezuela: Orchidaceae*. Caracas: Edicion Especial del Instituto Botanico.
13. Geerinck D, Vermeulen JJ, Pettersson B. 1992 *Flore d'Afrique Centrale: Orchidaceae, Seconde Partie*. Meise: Jardin botanique national de Belgique.
14. Hervouet J-M. 2018 *À la recherche des orchidées de Madagascar: sur les traces d'Henri Perrier de la Bâthie*. Mèze: Biotope éditions.
15. Johnson SD, Raguso RA. 2016 The long-tongued hawkmoth pollinator niche for native and invasive plants in Africa. *Ann Bot* **117**, 25–36. (doi:10.1093/aob/mcv137)
16. Jonsson L. 1981 *A monograph of the genus* Microcoelia *(Orchidaceae)*. Uppsala: Acta Universitatis Upsaliensis.
17. La Croix I. 2014 Aerangis*: exquisite African orchids to discover, identify and grow*. Portland, Oregon: Timber Press.
18. Cribb PJ. 1998 *Flora zambesiaca. Vol. 11, Pt. 2*. London: Flora Zambesiaca Managing Committee.
19. Linder HP, Kurzweil H. 1999 *Orchids of Southern Africa*. Rotterdam Brookfield: A. A. Balkema.
20. Llamacho JA, Larramendi J, Luer CA, Ackerman JD. 2005 *The orchids of Cuba = Las orquídeas de Cuba*. Lleida: Greta editores.

21. Luyt R, Johnson SD. 2001 Hawkmoth pollination of the African epiphytic orchid *Mystacidium venosum*, with special reference to flower and pollen longevity. *Plant Systematics and Evolution* **228**, 49–62. (doi:10.1007/s006060170036)
22. Luyt RP. 2002 Pollination and Evolution of the genus *Mystacidium* (Orchidaceae). University of Natal, Pietermaritzburg.
23. Martins DJ, Johnson SD. 2007 Hawkmoth pollination of aerangoid orchids in Kenya, with special reference to nectar sugar concentration gradients in the floral spurs. *American J of Botany* **94**, 650–659. (doi:10.3732/ajb.94.4.650)
24. Martins DJ, Johnson SD. 2013 Interactions between hawkmoths and flowering plants in East Africa: polyphagy and evolutionary specialization in an ecological context. *Biol J Linn Soc Lond* **110**, 199–213. (doi:10.1111/bij.12107)
25. McVaugh R. 1985 *Flora Novo-Galiciana: Orchidaceae*. Ann Arbor: The University of Michigan Press.
26. Nilsson LA, Johnsson L, Ralison L, Randrianjohany E. 1987 Angraecoid Orchids and Hawkmoths in Central Madagascar: Specialized Pollination Systems and Generalist Foragers. *Biotropica* **19**, 310. (doi:10.2307/2388628)
27. Perez-Vera F. 2003 *Les orchidées de Côte d'Ivoire*. Mèze [Paris]: Biotope IRD éd.
28. Perrier de La Bâthie H. 1939 *Flore de Madagascar: Orchidées (49e famille)*. Tananarive: Imprimerie officielle.
29. Peter CI, Venter N. 2017 Generalist, settling moth pollination in the endemic South African twig epiphyte, *Mystacidium pusillum* Harv. (Orchidaceae). *Flora* **232**, 16–21. (doi:10.1016/j.flora.2016.11.014)
30. Schlechter R. 1925 Die Orchidaceen der Insel Celebes. Part I. *Feddes Repert* **21**, 113–163. (doi:10.1002/fedr.19250210801)
31. Simo-Droissart M, Sonké B, Droissart V, Geerinck D, Lowry Ii PP, Stévart T. 2016 A taxonomic revision of *Angraecum* section *Dolabrifolia* (Orchidaceae, Angraecinae), with the description of a new species from Gabon. *Phytotaxa* **280**, 81. (doi:10.11646/phytotaxa.280.2.1)
32. Simo-Droissart M, Sonké B, Droissart V, Geerinck D, Micheneau C, Lowry PP, Plunkett GM, Hardy OJ, Stévart T. 2014 Taxonomic Revision of the Continental African Species of *Angraecum* Section *Pectinaria* (Orchidaceae). *Systematic Botany* **39**, 725–739. (doi:10.1600/036364414X682184)
33. Stewart J, Hermans J, Campbell B. 2006 *Angraecoid orchids: species from the African region*. Portland, Or: Timber Press.
34. Szlachetko DL, Olszewski TS. 2001 *Flore du Cameroun: Orchidaceae*. Yaoundé: Ministère de la recherche scientifique et technique.
35. Szlachetko DL, Sawicka M, Kras-Łapińska M. 2004 *Flore du Gabon: Orchidaceae*. Paris: Muséum national d'histoire naturelle, Département de systématique et évolution.
36. Wasserthal LT. 1997 The Pollinators of the Malagasy Star Orchids *Angraecum sesquipedale, A. sororium* and *A. compactum* and the Evolution of Extremely Long Spurs by Pollinator Shift. *Botanica Acta* **110**, 343–359. (doi:10.1111/j.1438-8677.1997.tb00650.x)
37. Verlynde S, Dubuisson J-Y, Stévart T, Simo-Droissart M, Geerinck D, Sonké B, Cawoy V, Descourvières P, Droissart V. 2013 Taxonomic revision of the genus *Bolusiella* (Orchidaceae, Angraecinae) with a new species from Cameroon, Burundi and Rwanda. *Phytotaxa* **114**, 1. (doi:10.11646/phytotaxa.114.1.1)
38. Williams LO, Allen PH, Dressler RL. 1980 *Orchids of Panama: a facsimile reprint of the Orchidaceae, flora of Panama*. St. Louis, Missouri: Missouri Botanical Garden.
39. Cribb P, Pollard BJ. 2002 New Orchid Discoveries in Western Cameroon. *Kew Bulletin* **57**, 653. (doi:10.2307/4110995)

Table S1. Sequences of 332 taxa used in the phylogenetic analyses

Taxa are listed alphabetically. Voucher specimens and GenBank accessions are indicated for each sequence. Herbarium acronyms where vouchers are housed are indicated in brackets. Newly generated sequences are indicated by "XX". Some samples collected in Rwanda are vouchered by photographs from Fischer *et al.* (2010). Missing data are indicated by "—". Information for ITS-1, matK and trnL–trnF is followed below by that of rps16, trnC–petN and ycf1.

| Taxon | ITS | | *matK* | | *trnL-trnF* | |
|---|---|---|---|---|---|---|
| | Genbank accession | Voucher [Herbarium] | Genbank accession | Voucher [Herbarium] | Genbank accession | Voucher [Herbarium] |
| *Aerangis arachnopus* (Rchb.f.) Schltr. | MH237055 | YAS 2436 [BRLU] | MK68552 4 | YAS 2436 [BRLU] | Farminhão *et al.* (2020) | YAS 2436 [BRLU] |
| *Aerangis articulata* (Rchb.f.) Schltr. | EF079444 | ex cult. Szlachetko s.n. | AY368389 | Jarrell s.n. | — | — |
| *Aerangis biloba* (Lindl.) Schltr. | MH237056 | YAS 2279 [BRLU] | MK68552 6 | YAS 2279 [BRLU] | Farminhão *et al.* (2020) | YAS 2279 [BRLU] |
| *Aerangis bouarensis* Chiron | XX | YAS 4286 [BRLU] | XX | YAS 4286 [BRLU] | XX | YAS 4286 [BRLU] |
| *Aerangis brachycarpa* (A.Rich.) T.Durand & Schinz | — | — | KU748263 | NMK: 1594.10176 | — | — |
| *Aerangis calantha* (Schltr.) Schltr. | MH237045 | YAS 2828 [BRLU] | MK68551 4 | YAS 2828 [BRLU] | — | — |
| *Aerangis citrata* (Thouars) Schltr. | DQ091600 | Whitten 1788 [FLAS] | DQ091337 | Whitten 1788 [FLAS] | — | — |
| *Aerangis collum-cygni* Summerh. | MH237057 | YAS 2779 [BRLU] | MK68552 7 | YAS 2779 [BRLU] | Farminhão *et al.* (2020) | YAS 2779 [BRLU] |
| *Aerangis confusa* J.Stewart | DQ091595 | Bytebier s.n. [EA] | DQ091332 | Bytebier s.n. [EA] | DQ091456 | Bytebier s.n. [EA] |
| *Aerangis coriacea* Summerh. | DQ091609 | Bytebier 562 [EA] | DQ091345 | Bytebier 562 [EA] | DQ091469 | Bytebier 562 [EA] |
| *Aerangis ellisii* (B.S.Williams) Schltr. | MH237255 | Chase 15080 [K] | KF557992 | Chase 15080 [K] | KF558204 | Chase 15080 [K] |
| *Aerangis fastuosa* (Rchb.f.) Schltr. | DQ091604 | Carlsward 402 [FLAS] | DQ091340 | Carlsward 402 [FLAS] | DQ091464 | Carlsward 402 [FLAS] |
| *Aerangis flexuosa* (Ridl.) Schltr. | MH237157 | Photo voucher [BRLU] | — | — | MH237541 | Photo voucher [BRLU] |
| *Aerangis gracillima* (Kraenzl.) J.C.Arends & J.Stewart | MH237060 | YAS 2302 [BRLU] | MK68553 0 | YAS 2302 [BRLU] | Farminhão *et al.* (2020) | YAS 1404 [BRLU] |
| *Aerangis gravenreuthii* (Kraenzl.) Schltr. | MH236944 | YAS 1565 [BRLU] | MK68541 5 | YAS 1565 [BRLU] | Farminhão *et al.* (2020) | YAS 1565 [BRLU] |
| *Aerangis* | DQ091607 | Carlsward 292 | Farminhão | Carlsward 292 | Farminhão *et* | Carlsward 292 |

| | | | | | | |
|---|---|---|---|---|---|---|
| *hariotiana* (Kraenzl.) P.J.Cribb & Carlsward | | [FLAS] | *et al.* (2020) | [FLAS] | *al.* (2020) | [FLAS] |
| *Aerangis hildebrandtii* (Rchb.f.) P.J.Cribb & Carlsward | DQ091608 | Kew accession 2616 [K] | DQ091344 | Kew accession 2616 [K] | Farminhão *et al.* (2020) | Kew accession 2616 [K] |
| *Aerangis kirkii* (Rchb.f.) Schltr. | DQ091596 | Bytebier 637 [EA] | DQ091333 | Bytebier 637 [EA] | Farminhão *et al.* (2020) | Bytebier 637 [EA] |
| *Aerangis kotschyana* (Rchb.f.) Schltr. | DQ091598 | Bytebier 671 [EA] | DQ091335 | Bytebier 671 [EA] | Farminhão *et al.* (2020) | Bytebier 671 [EA] |
| *Aerangis luteoalba* var. *rhodosticta* (Kraenzl.) J.Stewart | DQ091611 | Bytebier 691 [EA] | DQ091347 | Bytebier 691 [EA] | Farminhão *et al.* (2020) | Bytebier 691 [EA] |
| *Aerangis macrocentra* (Schltr.) Schltr. | MH237251 | Hermans 779 [K] | KF557923 | Hermans 779 [K] | — | — |
| *Aerangis modesta* (Hook.f.) Schltr. | DQ091603 | Carlsward 242 [FLAS] | DQ091341 | Carlsward 242 [FLAS] | Farminhão *et al.* (2020) | Carlsward 242 [FLAS] |
| *Aerangis monantha* Schltr. | XX | 618A234bis/1 [] | — | — | — | — |
| *Aerangis punctata* J.Stewart | MH237232 | No voucher (CM9) | KF557936 | No voucher (CM9) | KF558040 | No voucher (CM9) |
| *Aerangis somalensis* (Schltr.) Schltr. | DQ091612 | Bytebier 1549 [EA] | DQ091346 | Bytebier 1549 [EA] | Farminhão *et al.* (2020) | Bytebier 1549 [EA] |
| *Aerangis stelligera* Summerh. | MH237062 | YAS 2810 [BRLU] | MK685532 | YAS 2810 [BRLU] | MH237438 | YAS 2810 [BRLU] |
| *Aerangis thomsonii* (Rolfe) Schltr. | DQ091599 | Kirika 968 [EA] | DQ091336 | Kirika 968 [EA] | DQ091460 | Kirika 968 [EA] |
| *Aerangis ugandensis* Summerh. | MH237124 | Delepierre & Lebel 194 [BR] | MK685600 | Delepierre & Lebel 194 [BR] | Farminhão *et al.* (2020) | Delepierre & Lebel 194 [BR] |
| *Aerangis verdickii* (De Wild.) Schltr. | MH237125 | See p. 93 in The Orchids of Rwanda [BRLU] | MK685601 | See p. 93 in The Orchids of Rwanda [BRLU] | MH237508 | See p. 93 in The Orchids of Rwanda [BRLU] |
| *Aeranthes arachnites* (Thouars) Lindl. | DQ091759 | Carlsward 198 [FLAS] | KF557945 | Fournel 126 [REU] | KF558036 | Fournel 126 [REU] |
| *Aeranthes ecalcarata* H.Perrier | — | — | XX | 826A1362 [] | XX | 826A1362 [] |
| *Aeranthes grandiflora* Lindl. | Farminhão *et al.* (2020) | ANT18 | Farminhão *et al.* (2020) | ANT18 | Farminhão *et al.* (2020) | ANT18 |
| *Aeranthes leandriana* Bosser | Farminhão *et al.* (2020) | 13T135 [BRLU] | Farminhão *et al.* (2020) | 13T135 [BRLU] | Farminhão *et al.* (2020) | 13T135 [BRLU] |

| | | | | | | |
|---|---|---|---|---|---|---|
| *Aeranthes neoperrieri* Toill.-Gen., Ursch & Bosser | MH237219 | No voucher (CM6) | KF557953 | No voucher (CM6) | KF558059 | No voucher (CM6) |
| *Aeranthes nidus* Schltr. | Farminhão *et al.* (2020) | ANT14 | Farminhão *et al.* (2020) | ANT14 | Farminhão *et al.* (2020) | ANT14 |
| *Aeranthes peyrotii* Bosser | MH237262 | Chase 14644 [K] | KF557980 | Chase 14644 [K] | KF558061 | Chase 14644 [K] |
| *Aeranthes tenella* Bosser | MH237227 | Pailler 137 [REU] | KF557997 | Pailler 137 [REU] | KF558117 | Pailler 137 [REU] |
| *Aerides odorata* Lour. | AB217529 | TBG118480 | KF558231 | Chase 15081 [K] | KF558031 | Chase 15081 [K] |
| *Afropectinariella atlantica* (Droissart & Stévart) M.Simo & Stévart | KF672223 | JB Bambusa 130 [BRLU] | KF672284 | Van der Laan 1068 [BRLU] | KF672243 | Van der Laan 1068 [BRLU] |
| *Afropectinariella doratophylla* (Summerh.) M.Simo & Stévart | KF672204 | JB Bom Sucesso 1962.1 [BRLU] | KF672258 | JB Bom Sucesso 1962.1 [BRLU] | Farminhão *et al.* (2020) | Stévart 441 [BRLU] |
| *Afropectinariella gabonensis* (Summerh.) M.Simo & Stévart | MH236931 | YAS 1109 [BRLU] | MK68540 2 | YAS 1109 [BRLU] | Farminhão *et al.* (2020) | MBG 542 [BRLU] |
| *Afropectinariella pungens* (Schltr.) M.Simo & Stévart | KF672216 | YAS 2817 [BRLU] | KF672278 | BR 20090382-33 [BRLU] | Farminhão *et al.* (2020) | BR 20090383-34 [BRLU] |
| *Afropectinariella subulata* (Lindl.) M.Simo & Stévart | MH236911 | YAS 2480 [BRLU] | KF672285 | BR 20090388-39 [BRLU] | KF672241 | YAS 2503 [BRLU] |
| *Amesiella philippinensis* (Ames) Garay | DQ091718 | Carlsward 295 [FLAS] | EF079281 | Heidelberg BG 104625 | DQ091443 | Carlsward 295 [FLAS] |
| *Ancistrorhynchus capitatus* (Lindl.) Summerh. | MH237047 | Stévart & Pial 956 [BRLU] | MK68551 6 | Stévart & Pial 956 [BRLU] | Farminhão *et al.* (2020) | YAS 1057 [BRLU] |
| *Ancistrorhynchus cephalotes* (Rchb.f.) Summerh. | MH237118 | Nimba 198 [BRLU] | MK68559 5 | Nimba 198 [BRLU] | MH237502 | Nimba 198 [BRLU] |
| *Ancistrorhynchus clandestinus* (Lindl.) Schltr. | MH237064 | YAS 2745 [BRLU] | MK68553 4 | YAS 2831 [BRLU] | Farminhão *et al.* (2020) | YAS 2745 [BRLU] |
| *Ancistrorhynchus crystalensis* P.J.Cribb & Laan | MH237109 | Tchimbélé 18 [BRLU] | MK68558 5 | Tchimbélé 18 [BRLU] | Farminhão *et al.* (2020) | Tchimbélé 18 [BRLU] |
| *Ancistrorhynchus metteniae* (Kraenzl.) Summerh. | MH237121 | Nimba 203 [BRLU] | MK68553 5 | YAS 2214 [BRLU] | Farminhão *et al.* (2020) | Nimba 203 [BRLU] |
| *Ancistrorhynchus* | MH236973 | BR 20090367- | MK68544 | BR 20090367- | Farminhão *et* | BR 20090367- |

| | | | | | | |
|---|---|---|---|---|---|---|
| *ovatus* Summerh. | | 18 | 5 | 18 | *al.* (2020) | 18 |
| *Ancistrorhynchus recurvus* Finet | MH236935 | YAS 2785 [BRLU] | DQ091354 | No voucher | Farminhão *et al.* (2020) | YAS 2785 [BRLU] |
| *Ancistrorhynchus refractus* (Kraenzl.) Summerh. | MH236963 | BR 20090460-14 | MK68543 6 | BR 20090460-14 | Farminhão *et al.* (2020) | BR 20090460-14 |
| *Ancistrorhynchus schumannii* (Kraenzl.) Summerh. | MH236938 | YAS 895 [BRLU] | MK68554 2 | YAS 1580 [BRLU] | Farminhão *et al.* (2020) | YAS 895 [BRLU] |
| *Ancistrorhynchus serratus* Summerh. | MH237069 | YAS 2381 [BRLU] | MK68553 9 | YAS 2381 [BRLU] | Farminhão *et al.* (2020) | YAS 2381 [BRLU] |
| *Ancistrorhynchus straussii* (Schltr.) Schltr. | MH236966 | BR20090369-20 | MK68543 9 | BR20090369-20 | Farminhão *et al.* (2020) | BR20090369-20 |
| *Ancistrorhynchus tenuicaulis* Summerh. | MH237126 | See p. 95 in The Orchids of Rwanda | MK68560 2 | See p. 95 in The Orchids of Rwanda | Farminhão *et al.* (2020) | |
| *Angraecopsis elliptica* Summerh. | Farminhão *et al.* (2020) | YAS 4473 [BRLU] | Farminhão *et al.* (2020) | YAS 4473 [BRLU] | Farminhão *et al.* (2020) | YAS 4473 [BRLU] |
| *Angraecopsis gracillima* (Rolfe) Summerh. | Farminhão *et al.* (2020) | Farminhão 231 [BRLU] | Farminhão *et al.* (2020) | Droissart 978 [BRLU] | Farminhão *et al.* (2020) | Droissart 978 [BRLU] |
| *Angraecopsis ischnopus* (Schltr.) Schltr. | Farminhão *et al.* (2020) | Droissart 689 [BRLU] | Farminhão *et al.* (2020) | Droissart 689 [BRLU] | Farminhão *et al.* (2020) | Droissart 689 [BRLU] |
| *Angraecopsis parviflora* (Thouars) Schltr. | Farminhão *et al.* (2020) | YAS 3830 [BRLU] | Farminhão *et al.* (2020) | YAS 3830 [BRLU] | Farminhão *et al.* (2020) | YAS 3830 [BRLU] |
| *Angraecopsis* sp. nov. Oku | XX | YAS 3645 [BRLU] | XX | YAS 3645 [BRLU] | XX | YAS 3645 [BRLU] |
| *Angraecopsis tenerrima* Kraenzl. | MF980229 | No voucher | MG25022 2 | No voucher | MG250241 | No voucher |
| *Angraecopsis tridens* (Lindl.) Schltr. | XX | Droissart 693 [BRLU] | XX | Droissart 693 [BRLU] | XX | Droissart 693 [BRLU] |
| *Angraecum acutipetalum* Schltr. | — | — | Farminhão *et al.* (2020) | 14T21 [TAN] | Farminhão *et al.* (2020) | 14T21 [TAN] |
| *Angraecum amplexicaule* Toill.-Gen. & Bosser | — | — | KT826791 | Andriananjama nantsoa & al. 00203468 [MT] | KT826929 | Andriananjama nantsoa & al. 00203468 [MT] |
| *Angraecum appendiculatum* Frapp. ex Cordem. | MH237203 | Pailler 107 [REU] | KF557933 | Pailler 107 [REU] | KF558130 | Pailler 107 [REU] |
| *Angraecum arachnites* Schltr. | — | — | KF557990 | Hermans 4241 [K] | Farminhão *et al.* (2020) | Hermans 4241 [K] |
| *Angraecum* | Farminhão *et* | Micheneau 1 | KF558023 | Micheneau 1 | KF558140 | Micheneau 1 |

| | | | | | | |
|---|---|---|---|---|---|---|
| *borbonicum* Bosser | *al.* (2020) | [BRLU] | | [BRLU] | | [BRLU] |
| *Angraecum bracteosum* Balf.f. & S.Moore | XX | Micheneau 7 [REU] | KF557998 | Micheneau 7 [REU] | KF558266 | Micheneau 7 [REU] |
| *Angraecum breve* Schltr. | XX | AMB5173 [] | XX | AMB5173 [] | — | — |
| *Angraecum cadetii* Bosser | MH237229 | Pailler 157 [REU] | KF557986 | Pailler 157 [REU] | KF558122 | Pailler 157 [REU] |
| *Angraecum* cf. *elephantinum* | XX | AMB5135 | XX | AMB5135 | XX | AMB5135 |
| *Angraecum chaetopodum* Schltr. | — | — | KT826786. | Andriananjamanantsoa & al. 00203462 [MT] | KT826924 | Andriananjamanantsoa & al. 00203462 [MT] |
| *Angraecum clavigerum* Ridl. | — | — | Farminhão *et al.* (2020) | 42T60 [TAN] | Farminhão *et al.* (2020) | 42T60 [TAN] |
| *Angraecum compactum* Schltr. | — | — | KT826796 | Andriananjamanantsoa & al. 00203420 [MT] | KT826934 | Andriananjamanantsoa & al. 00203420 [MT] |
| *Angraecum conchiferum* Lindl. | DQ091748 | Bytebier 616 [EA] | DQ091414 | Bytebier 616 [EA] | DQ091539 | Bytebier 616 [EA] |
| *Angraecum cornigerum* Cordem. | MH237233 | Pailler 176 [REU] | KF558015 | Pailler 176 [REU] | KF558135 | Pailler 176 [REU] |
| *Angraecum corrugatum* (Cordem.) Micheneau | — | — | KF558003 | Pailler 106 [REU] | KF558106 | Pailler 106 [REU] |
| *Angraecum costatum* Frapp. ex Cordem. | — | — | KF557942 | Pailler 174 [REU] | KF558111 | Pailler 174 [REU] |
| *Angraecum cucullatum* Thouars | MH237217 | Pailler 108 [REU] | KF557927 | Pailler 108 [REU] | KF558245 | Pailler 108 [REU] |
| *Angraecum didieri* (Baill. ex Finet) Schltr. | XX | AMB741.1 [] | XX | AMB741.1 [] | XX | AMB741.1 [] |
| *Angraecum dives* Rolfe | DQ091756 | Marimoto 42 [EA] | DQ091422 | Marimoto 42 [EA] | Farminhão *et al.* (2020) | Marimoto 42 [EA] |
| *Angraecum eburneum* subsp. eburneum Bory | MH237264 | unvouchered [K] | KF558000 | unvouchered [K] | KF558261 | unvouchered [K] |
| *Angraecum eburneum* subsp. *superbum* (Thouars) H.Perrier | MH237265 | Carter 761 [K] | KF557937 | Carter 761 [K] | KF558193 | Carter 761 [K] |
| *Angraecum equitans* Schltr. | — | — | Farminhão *et al.* (2020) | Verlynde 52 [BRLU] | Farminhão *et al.* (2020) | Verlynde 52 [BRLU] |

| | | | | | | |
|---|---|---|---|---|---|---|
| *Angraecum expansum* Thouars | MH237234 | Micheneau 2 [BRLU] | KF557974 | Micheneau 2 [BRLU] | KF558087 | Micheneau 2 [BRLU] |
| *Angraecum filicornu* Thouars | — | — | KT826823 | Andriananjamanantsoa & al. 00203460 [MT] | KT826961 | Andriananjamanantsoa & al. 00203460 [MT] |
| *Angraecum florulentum* Rchb.f. | MH237267 | unvouchered [K] | KF557934 | unvouchered [K] | KF558248 | unvouchered [K] |
| *Angraecum humblotianum* Schltr. | KF672199 | Simo M. 217 [BRLU] | KF662345 | Simo M. 217 [BRLU] | KF672272 | Simo M. 217 [BRLU] |
| *Angraecum huntleyoides* Schltr. | MH237246 | Hermans 4248.1 [K] | KF557961 | Hermans 4248.1 [K] | KF558143 | Hermans 4248.1 [K] |
| *Angraecum leonis* (Rchb.f.) André | DQ091735 | Carlsward 390 [FLAS] | DQ091426 | Carlsward 390 [FLAS] | Farminhão *et al.* (2020) | Carlsward 390 [FLAS] |
| *Angraecum liliodorum* Frapp. ex Cordem. | MH237230 | No voucher | KF557982 | No voucher | KF558100 | No voucher |
| *Angraecum linearifolium* Garay | MH237252 | Hermans 3258 [K] | Farminhão *et al.* (2020) | 297T86 [BRLU] | Farminhão *et al.* (2020) | 297T86 [BRLU] |
| *Angraecum longicalcar* (Bosser) Senghas | DQ091739 | unvouchered [K] | DQ091419 | unvouchered [K] | — | — |
| *Angraecum magdalenae* Schltr. & H.Perrier | — | — | KF557995 | No voucher | KF558123 | No voucher |
| *Angraecum mauritianum* (Poir.) Frapp. | MH237235 | No voucher | KF558020 | No voucher | KF558080 | No voucher |
| *Angraecum moratii* Bosser | — | — | KT826770 | Andriananjamanantsoa & al. 00203447 [MT] | KT826908 | Andriananjamanantsoa & al. 00203447 [MT] |
| *Angraecum multiflorum* Thouars | MH237213 | Pailler 154 [REU] | KF557948 | Pailler 154 [REU] | KF558240 | Pailler 154 [REU] |
| *Angraecum nanum* Frapp. ex Cordem. | — | — | KT826798 | Andriananjamanantsoa & al. 00203325 [MT] | — | — |
| *Angraecum obesum* H.Perrier | JQ905393 | Rakotoarivelo & al. 008 [TAN] | JQ905332 | Rakotoarivelo & al. 008 [TAN] | JQ905515 | Rakotoarivelo & al. 008 [TAN] |
| *Angraecum oblongifolium* Toill.-Gen. & Bosser | — | — | KT826771 | Andriananjamanantsoa & al. 00203446 [MT] | KT826909 | Andriananjamanantsoa & al. 00203446 [MT] |
| *Angraecum obversifolium* Frapp. ex Cordem. | XX | Micheneau & Pailler 8 [REU] | KF557962 | Micheneau & Pailler 8 [REU] | KF558259 | Micheneau & Pailler 8 [REU] |

| | | | | | | |
|---|---|---|---|---|---|---|
| *Angraecum panicifolium* H.Perrier | KF672205 | Simo 215 [BRLU] | KF672277 | Simo 215 [BRLU] | KF672244 | Simo 215 [BRLU] |
| *Angraecum pectinatum* Thouars | MH237222 | No voucher (CM6h) | KF557952 | No voucher (CM6h) | KF558127 | No voucher (CM6h) |
| *Angraecum penzigianum* Schltr. | — | — | KT826772 | Andriananjama nantsoa & al. 00203466 [MT] | KT826910 | Andriananjama nantsoa & al. 00203466 [MT] |
| *Angraecum protensum* Schltr. | MH237187 | MO4617229 [NY] | MK68564 8 | MO4617229 [NY] | — | — |
| *Angraecum pseudodidieri* H.Perrier | Farminhão *et al.* (2020) | No voucher (CM6) | KF558002 | No voucher (CM6) | KF558032 | No voucher (CM6) |
| *Angraecum ramosum* Thouars | MH237205 | Micheneau & Pailler 73 [REU] | Farminhão *et al.* (2020) | Micheneau & Pailler 73 [REU] | KF558037 | Micheneau & Pailler 73 [REU] |
| *Angraecum rutenbergianum* Kraenzl. | DQ091743 | Carlsward 300 [FLAS] | DQ091423 | Carlsward 300 [FLAS] | DQ091548 | Carlsward 300 [FLAS] |
| *Angraecum sacciferum* Lindl. | — | — | KF558028 | Bytebier 2226 [NBG] | KF558101 | Bytebier 2226 [NBG] |
| *Angraecum sedifolium* Schltr. | MH237253 | Hermans 3802 [K] | XX | AMB5167 [] | XX | AMB5167 [] |
| *Angraecum serpens* (H.Perrier) Bosser | — | — | KT826773 | Andriananjama nantsoa & al. 00203438 [MT] | KT826911 | Andriananjama nantsoa & al. 00203438 [MT] |
| *Angraecum sesquipedale* Thouars | MH237223 | No voucher (CM6j) | KF557996 | No voucher (CM6j) | KF558109 | No voucher (CM6j) |
| *Angraecum sororium* Schltr. | — | — | KT826825 | Andriananjama nantsoa & al. 00203400 [MT] | KT826963 | Andriananjama nantsoa & al. 00203400 [MT] |
| *Angraecum* sp. | XX | AMB5175 [] | XX | AMB5175 [] | XX | AMB5175 [] |
| *Angraecum striatum* Thouars | XX | Micheneau 4 [REU] | KF557991 | Micheneau 4 [REU] | KF558214 | Micheneau 4 [REU] |
| *Angraecum tenuifolium* Frapp. ex Cordem. | MH237215 | Pailler 116 [REU] | KF557968 | Pailler 116 [REU] | KF558035 | Pailler 116 [REU] |
| *Angraecum teretifolium* Ridl. | MH237212 | No voucher (CM3f) | Farminhão *et al.* (2020) | No voucher (CM3f) | KF558132 | No voucher (CM3f) |
| *Angraecum triangulifolium* Senghas | — | — | KT826794 | Andriananjama nantsoa & al. 00203442 [MT] | KT826932 | Andriananjama nantsoa & al. 00203442 [MT] |
| *Angraecum viguieri* Schltr. | — | — | Farminhão *et al.* (2020) | Verlynde 51 [BRLU] | Farminhão *et al.* (2020) | Verlynde 51 [BRLU] |

| | | | | | | |
|---|---|---|---|---|---|---|
| *Aziza trilobata* (Summerh.) Farminhão & D'haijère | Farminhão *et al.* (2020b) | Farminhão 11 [BRLU] | Farminhão *et al.* (2020b) | Farminhão 11 [BRLU] | — | — |
| *Beclardia macrostachya* (Thouars) A.Rich. | MH237245 | Fournel 276 [REU] | Farminhão *et al.* (2020) | 27T13 | KF558039 | Fournel 276 [REU] |
| *Bolusiella fractiflexa* Droissart, Stévart & Verlynde | MH237001 | Delepierre & Lebel 216 [BR] | MK685411 | YAS 1357 [BRLU] | Farminhão *et al.* (2020) | YAS 1357 [BRLU] |
| *Bolusiella iridifolia* (Rolfe) Schltr. | DQ091665 | Bytebier 1113 [EA] | MK685474 | Photo voucher (see p. 106 in The Orchids of Rwanda) | Farminhão *et al.* (2020) | Photo voucher (see p. 106 in The Orchids of Rwanda) |
| *Bolusiella maudiae* (Bolus) Schltr. | DQ091664 | Bytebier 485 [EA] | DQ091355 | Bytebier 485 [EA] | Farminhão *et al.* (2020) | Bytebier 485 [EA] |
| *Bolusiella talbotii* (Rendle) Summerh. | MH237122 | Nimba 40 [BRLU] | MK685598 | Nimba 40 [BRLU] | Farminhão *et al.* (2020) | Nimba 40 [BRLU] |
| *Bolusiella zenkeri* (Kraenzl.) Schltr. | MH237037 | YAS 2442 [BRLU] | Farminhão *et al.* (2020) | MBG 601 [BRLU] | Farminhão *et al.* (2020) | YAS 2442 [BRLU] |
| *Calyptrochilum aurantiacum* (P.J.Cribb & Laan) Stévart, M.Simo & Droissart | MH237088 | YAS 2773 [BRLU] | MK685561 | YAS 2773 [BRLU] | Farminhão *et al.* (2020) | YAS 2773 [BRLU] |
| *Calyptrochilum christyanum* (Rchb.f.) Summerh. | MH237039 | YAS 1799 [BRLU] | MK685508 | YAS 1799 [BRLU] | Farminhão *et al.* (2020) | YAS 1799 [BRLU] |
| *Calyptrochilum emarginatum* (Afzel. ex Sw.) Schltr. | MH237043 | YAS 2735 [BRLU] | MK685512 | YAS 2735 [BRLU] | Farminhão *et al.* (2020) | YAS 2735 [BRLU] |
| *Campylocentrum brenesii* Schltr. | LT706522 | M. Blanco 2139 [USJ] | MK685658 | M. Blanco 2139 [USJ] | MH237589 | M. Blanco 2139 [USJ] |
| *Campylocentrum fasciola* (Lindl.) Cogn. | DQ091564 | Carlsward 301 [FLAS] | DQ091321 | Carlsward 301 [FLAS] | — | — |
| *Campylocentrum jamaicense* (Rchb.f. & Wullschl.) Benth. ex Fawc. | AY147219 | Carlsward 12 (FLAS) | AF506346 | Ackerman 3341 [UPRRP] | AF506325 | Ackerman 3341 [UPRRP] |
| *Campylocentrum minutum* C.Schweinf. | MH237188 | MO4904973 [NY] | — | — | MH237576 | MO4904973 [NY] |
| *Campylocentrum neglectum* (Rchb.f. & Warm.) Cogn. | AF506297 | Carlsward 272 [FLAS] | AF506345 | Carlsward 272 [FLAS] | Farminhão *et al.* (2020) | Carlsward 272 [FLAS] |
| *Campylocentrum pachyrrhizum* (Rchb.f.) Rolfe | AF506301 | Ackerman s.n. [UPRRP] | AF506350 | Ackerman s.n. [UPRRP] | — | — |

| | | | | | | |
|---|---|---|---|---|---|---|
| *Campylocentrum poeppigii* (Rchb.f.) Rolfe | LT718679 | Bogarín 2218 [CR] | AF506351 | Carnevali 4507 [CICY] | — | — |
| *Campylocentrum stenanthum* Schltr. | AF506298 | Carlsward 180 [FLAS] | AF506347 | Carlsward 180 [FLAS] | — | — |
| *Campylocentrum tyrridion* Garay & Dunst. ex Foldats | AF506305 | Carnevali 5145 [FLAS] | MK68565 9 | Carnevali 5145 [FLAS] | MH237591 | Carnevali 5145 [FLAS] |
| *Cleisostoma arietinum* (Rchb.f.) Garay | DQ091694 | Carlsward 211 [SEL] | KJ33550 | Z.J.Liu 6991 [] | — | — |
| *Conchograecum affine* (Schltr.) Szlach., Grochocka, Oledrz. & Mytnik | MH237105 | BTO 95 [BRLU] | MK68558 1 | BTO 95 [BRLU] | Farminhão *et al.* (2020) | BTO 95 [BRLU] |
| *Conchograecum angustum* (Rolfe) Szlach., Grochocka, Oledrz. & Mytnik | MH236914 | YAS 1226 [BRLU] | MK68539 0 | YAS 1226 [BRLU] | Farminhão *et al.* (2020) | YAS 1226 [BRLU] |
| *Conchograecum claessensii* (De Wild.) Szlach., Grochocka, Oledrz. & Mytnik | MH237067 | YAS 3136 [BRLU] | MK68553 7 | YAS 3136 [BRLU] | Farminhão *et al.* (2020) | YAS 3136 [BRLU] |
| *Conchograecum cribbianum* (Szlach. & Olszewski) Szlach., Grochocka, Oledrz. & Mytnik | XX | MBG249 [BRLU] | XX | MBG249 [BRLU] | XX | MBG249 [BRLU] |
| *Conchograecum cultriforme* (Summerh.) Szlach., Grochocka, Oledrz. & Mytnik | AF506321 | Carlsward 298 [FLAS] | AF506364 | Carlsward 298 [FLAS] | Farminhão *et al.* (2020) | Carlsward 298 [FLAS] |
| *Conchograecum erectum* (Summerh.) Szlach., Grochocka, Oledrz. & Mytnik | DQ091566 | Bytebier 801 [EA] | DQ091323 | Bytebier 801 [EA] | Farminhão *et al.* (2020) | Bytebier 801 [EA] |
| *Conchograecum geerinckianum* (Stévart & Jecmenica) Szlach., Grochocka, Oledrz. & Mytnik | XX | SIB762 [BRLU] | XX | SIB762 [BRLU] | XX | SIB762 [BRLU] |
| *Conchograecum lanceolatum* (Ječmenica, Stévart & Droissart) Szlach., Grochocka, Oledrz. & Mytnik | MH237163 | BTO 23 [BRLU] | MK68563 0 | BTO 23 [BRLU] | Farminhão *et al.* (2020) | BTO 23 [BRLU] |
| *Conchograecum* | Farminhão *et* | YAS 2836 | MK68554 | YAS 2836 | Farminhão *et* | YAS 2836 |

| | | | | | | |
|---|---|---|---|---|---|---|
| *moandense* (De Wild.) Szlach., Grochocka, Oledrz. & Mytnik | *al.* (2020) | [BRLU] | 3 | [BRLU] | *al.* (2020) | [BRLU] |
| *Conchograecum multinominatum* (Rendle) Szlach., Grochocka, Oledrz. & Mytnik | MH236916 | YAS 696 [BRLU] | MK68539 1 | YAS 696 [BRLU] | Farminhão *et al.* (2020) | YAS 696 [BRLU] |
| *Conchograecum oliveirae* (Stévart & Jecmenica) Szlach., Grochocka, Oledrz. & Mytnik | Farminhão *et al.* (2020) | Bom Sucesso 1617 [BRLU] | Farminhão *et al.* (2020) | Bom Sucesso 1617 [BRLU] | Farminhão *et al.* (2020) | Bom Sucesso 1617 [BRLU] |
| *Conchograecum reygaertii* (De Wild.) Szlach., Grochocka, Oledrz. & Mytnik | MH237072 | YAS 2829 [BRLU] | MK68554 5 | YAS 2829 [BRLU] | Farminhão *et al.* (2020) | YAS 2829 [BRLU] |
| *Conchograecum sanfordii* (P.J.Cribb & B.J.Pollard) Szlach., Grochocka, Oledrz. & Mytnik | MH237072 | YAS 1136 [BRLU] | MK68554 5 | YAS 1136 [BRLU] | Farminhão *et al.* (2020) | YAS 1136 [BRLU] |
| *Conchograecum umbrosum* (P.J.Cribb) Szlach., Grochocka, Oledrz. & Mytnik | MH237184 | MO3262138 [NY] | — | — | MH237573 | MO3262138 [NY] |
| *Cryptopus paniculatus* H.Perrier | MH237248 | Hermans 5392 [K] | DQ091327 | Hermans 5392 [K] | Farminhão *et al.* (2020) | 2073A4951 [TAN] |
| *Cyrtorchis arcuata* (Lindl.) Schltr. | MH236958 | BR 20090398-49 | MK68543 0 | BR 20090398-49 | Farminhão *et al.* (2020) | BR 20090398-49 |
| *Cyrtorchis arcuata* subsp. *whytei* (Rolfe) Summerh. | Farminhão *et al.* (2020) | YAS 3599 [BRLU] | Farminhão *et al.* (2020) | YAS 3599 [BRLU] | Farminhão *et al.* (2020) | YAS 3599 [BRLU] |
| *Cyrtorchis aschersonii* (Kraenzl.) Schltr. | MH237100 | YAS 2434 [BRLU] | MK68557 4 | YAS 2434 [BRLU] | Farminhão *et al.* (2020) | YAS 2434 [BRLU] |
| *Cyrtorchis chailluana* (Hook.f.) Schltr. | MH236968 | BR 19750114 | MK68544 1 | BR 19750114 | MK721986 | BR 19750114 |
| *Cyrtorchis hamata* (Rolfe) Schltr. | Farminhão *et al.* (2020) | YAS 4671 [BRLU] | Farminhão *et al.* (2020) | YAS 4671 [BRLU] | Farminhão *et al.* (2020) | YAS 4671 [BRLU] |
| *Cyrtorchis henriquesiana* (Ridl.) Schltr. | MH237104 | BTO 99 [BRLU] | MK68558 0 | BTO 99 [BRLU] | MH237488. | BTO 99 [BRLU] |
| *Cyrtorchis letouzeyi* Szlach. & Olszewski | MH237050 | YAS 253 [BRLU] | MK68551 9 | YAS 253 [BRLU] | MH237425 | YAS 253 [BRLU] |

| | | | | | | |
|---|---|---|---|---|---|---|
| *Cyrtorchis monteiroae* (Rchb.f.) Schltr. | MH237155 | Bom Sucesso 2142.1 [BRLU] | MK685625 | Bom Sucesso 2142.1 [BRLU] | Farminhão *et al.* (2020) | Bom Sucesso 2142.1 [BRLU] |
| *Cyrtorchis praetermissa* Summerh. | MH237135 | Photo voucher (see pp. 168-169 in The Orchids of Rwanda | MK685611 | Photo voucher (see pp. 168-169 in The Orchids of Rwanda | Farminhão *et al.* (2020) | Photo voucher (see pp. 168-169 in The Orchids of Rwanda |
| *Cyrtorchis ringens* (Rchb.f.) Summerh. | MH237137 | Photo voucher (see pp. 166-167 in The Orchids of Rwanda | MK685613 | Photo voucher (see pp. 166-167 in The Orchids of Rwanda | Farminhão *et al.* (2020) | Photo voucher (see pp. 166-167 in The Orchids of Rwanda |
| *Dendrophylax alcoa* Dod | AF506307 | Ackerman 2773 [UPRRP] | — | — | — | — |
| *Dendrophylax barrettiae* Fawc. & Rendle | AF506308 | Carlsward 199 [FLAS] | AF506353 | Carlsward 199 [FLAS] | AF506330 | Carlsward 199 [FLAS] |
| *Dendrophylax fawcettii* Rolfe | AF506309 | Whitten 1939 [FLAS] | AF506354 | Whitten 1939 [FLAS] | AF506331 | Whitten 1939 [FLAS] |
| *Dendrophylax filiformis* (Sw.) Benth. ex Fawc. | AF506296 | Whitten 1842 [FLAS] | AF506344 | Whitten 1842 [FLAS] | — | — |
| *Dendrophylax funalis* (Sw.) Benth. ex Rolfe | AY147221 | Carlsward 199 [FLAS] | Farminhão *et al.* (2020) | Carlsward 302 [FLAS] | KF558067 | Carlsward 302 [FLAS] |
| *Dendrophylax lindenii* (Lindl.) Benth. ex Rolfe | MH237174 | unvouchered | AF506362 | photo voucher [FLAS] | AF506338 | photo voucher [FLAS] |
| *Dendrophylax megarhizus* Molgo & Carnevali | AF506314 | Carnevali 5907 [CICY] | AF506357 | Carnevali 5907 [CICY] | AF506335 | Carnevali 5907 [CICY] |
| *Dendrophylax porrectus* (Rchb.f.) Carlsward & Whitten | AY147223 | Carlsward 329 [FLAS] | JN176110 | Carlsward 330 [FLAS] | — | — |
| *Dendrophylax sallei* (Rchb.f.) Benth. ex Rolfe | AY147225 | Whitten 1945 [JBSD] | AY147239 | Whitten 1945 [JBSD] | AY147234 | Whitten 1945 [JBSD] |
| *Dendrophylax varius* (Aubl.) Urb. | AY147222 | Whitten 1960 [JBSD] | AY147236 | Whitten 1960 [JBSD] | AY147230 | Whitten 1960 [JBSD] |
| *Diaphananthe bidens* (Afzel. ex Sw.) Schltr. | Farminhão *et al.* (2020) | YAS 2340 [BRLU] | Farminhão *et al.* (2020) | YAS 2340 [BRLU] | Farminhão *et al.* (2020) | YAS 2152 [BRLU] |
| *Diaphananthe fragrantissima* (Rchb.f.) Schltr. | DQ091618 | Kirika 536 [EA] | MK685548 | YAS 1862 [BRLU] | Farminhão *et al.* (2020) | Kirika 536 [EA] |
| *Diaphananthe ichneumonea* (Lindl.) P.J.Cribb & Carlsward | MH236949 | YAS 889 [BRLU] | MK685420 | YAS 889 [BRLU] | Farminhão *et al.* (2020) | YAS 889 [BRLU] |

| | | | | | | |
|---|---|---|---|---|---|---|
| *Diaphananthe lebelii* (Eb.Fisch. & Killmann) Descourv. & Stévart | MH237006 | HOLOTYPE | Farminhão *et al.* (2020) | HOLOTYPE | Farminhão *et al.* (2020) | HOLOTYPE |
| *Diaphananthe lecomtei* (Finet) P.J.Cribb & Carlsward | MH237044 | YAS 2336 [BRLU] | MK685513 | YAS 2336 [BRLU] | Farminhão *et al.* (2020) | YAS 2336 [BRLU] |
| *Diaphananthe lorifolia* Summerh. | DQ091619 | Bytebier 346 [EA] | MK685655 | Bytebier 346 [EA] | Farminhão *et al.* (2020) | Bytebier 346 [EA] |
| *Diaphananthe odoratissima* (Rchb.f.) P.J.Cribb & Carlsward | MH237042 | YAS 1217 [BRLU] | MK685511 | YAS 1217 [BRLU] | Farminhão *et al.* (2020) | YAS 1217 [BRLU] |
| *Diaphananthe pellucida* (Lindl.) Schltr. | Farminhão *et al.* (2020) | YAS 1317 [BRLU] | DQ091377 | Carlsward 241 [FLAS] | — | — |
| *Diaphananthe sarcophylla* (Schltr. ex Prain) P.J.Cribb & Carlsward | DQ091621 | Bytebier 339 [EA] | MK685603 | Photo voucher (see p. 152 in The Orchids of Rwanda) | Farminhão *et al.* (2020) | Photo voucher (see p. 152 in The Orchids of Rwanda) |
| *Diaphananthe sarcorhynchoides* J.B.Hall | MH237110 | MBG 61 [BRLU] | MK685586 | MBG 61 [BRLU] | Farminhão *et al.* (2020) | MBG 61 [BRLU] |
| *Diaphananthe spiralis* (Stévart & Droissart) P.J.Cribb & Carlsward | MH237052 | YAS 1842 [BRLU] | MK685521 | YAS 1842 [BRLU] | MH237427 | YAS 1842 [BRLU] |
| *Diaphananthe vesicata* (Lindl.) P.J.Cribb & Carlsward | MH237063 | YAS 312 [BRLU] | KF557976 | Chase 14645 [K] | KF558083 | Chase 14645 [K] |
| *Dolabrifolia aporoides* (Summerh.) Szlach. & Romowicz | KF672202 | Bom Sucesso 946 [BRLU] | MK685444 | BR 20090370-21 | KF672242 | Bom Sucesso 946 [BRLU] |
| *Dolabrifolia bancoensis* (Burg) Szlach. & Romowicz | MH236909 | YAS 2187 [BRLU] | MK685385 | YAS 2187 [BRLU] | Farminhão *et al.* (2020) | YAS 2187 [BRLU] |
| *Dolabrifolia biteaui* (M.Simo & Stévart) M.Simo | KX060051 | BR20090387-38 | KX060071 | BR20090387-38 | Farminhão *et al.* (2020) | BTO 19 [BRLU] |
| *Dolabrifolia disticha* (Lindl.) Szlach. & Romowicz | KX060062 | YAS 62 [BRLU] | KF672265 | YAS 924 [BRLU] | KX060102 | YAS 62 [BRLU] |
| *Dolabrifolia podochiloides* (Schltr.) Szlach. & Romowicz | KX060045 | MBG 656 [BRLU] | KX060063 | Bambusa 159 [BRLU] | KX060085 | MBG 656 [BRLU] |

| | | | | | | |
|---|---|---|---|---|---|---|
| *Eichlerangraecum angustipetalum* comb. ined. | MH236910 | YAS 112 [BRLU] | MK685387 | YAS 112 [BRLU] | XX | YAS 112 [BRLU] |
| *Eichlerangraecum birrimense* (Rolfe) Szlach., Mytnik & Grochocka | — | — | MK685544 | YAS 399 [BRLU] | Farminhão *et al.* (2020) | YAS 399 [BRLU] |
| *Eichlerangraecum eichlerianum* var. *eichlerianum* (Kraenzl.) Szlach., Mytnik & Grochocka | MH237170 | BTO170 [BRLU] | MK685622 | SIB 1406 [BRLU] | MH237556 | BTO170 [BRLU] |
| *Eichlerangraecum eichlerianum* var. *curvicalcaratum* (Szlach. & Olszewski) Szlach., Mytnik & Grochocka | MH236907 | YAS 944 [BRLU] | MK685383 | YAS 944 [BRLU] | MH237272 | YAS 944 [BRLU] |
| *Eichlerangraecum infundibulare* (Lindl.) Szlach., Mytnik & Grochocka | MH237268 | No voucher (CM22961) | KF557977 | No voucher (CM22961) | KF558077 | No voucher (CM22961) |
| *Erasanthe henrici* (Schltr.) P.J.Cribb, Hermans & D.L.Roberts | MH237260 | No voucher (CM22946) | Farminhão *et al.* (2020) | No voucher (CM22946) | KF558102 | No voucher (CM22946) |
| *Eurychone galeandrae* (Rchb.f.) Schltr. | DQ091614 | Carlsward 293 [FLAS] | DQ091349 | Carlsward 293 [FLAS] | DQ091473 | Carlsward 293 [FLAS] |
| *Eurychone rothschildiana* (O'Brien) Schltr. | MH237117 | Traoré 141 [BRLU] | MK685593 | Traoré 141 [BRLU] | Farminhão *et al.* (2020) | Traoré 141 [BRLU] |
| *Jumellea alionae* P.J.Cribb | JQ905444 | Rakotoarivelo & al. 200 [TAN] | JQ905383 | Rakotoarivelo & al. 200 [TAN] | JQ905566 | |
| *Jumellea ambongensis* Schltr. | JQ905390 | Rakotoarivelo & al. 131 [TAN] | JQ905329 | Rakotoarivelo & al. 131 [TAN] | JQ905512 | Rakotoarivelo & al. 131 [TAN] |
| *Jumellea amplifolia* Schltr. | MH237225 | No voucher (CM6l) | KF558018 | No voucher (CM6l) | KF558110 | No voucher (CM6l) |
| *Jumellea anjouanensis* (Finet) H.Perrier | JQ905431 | Rakotoarivelo *et al.* 203 [TAN] | JQ905370 | Rakotoarivelo *et al.* 203 [TAN] | JQ905492 | Rakotoarivelo *et al.* 203 [TAN] |
| *Jumellea arachnantha* (Rchb.f.) Schltr. | JQ905439 | Rakotoarivelo *et al.* 136 [TAN] | JQ905378 | Rakotoarivelo *et al.* 136 [TAN] | JQ905500 | Rakotoarivelo *et al.* 136 [TAN] |
| *Jumellea arborescens* H.Perrier | JQ905398 | Rakotoarivelo & al. 202 [TAN] | JQ905337 | Rakotoarivelo & al. 202 [TAN] | JQ905520 | Rakotoarivelo & al. 202 [TAN] |

| | | | | | | |
|---|---|---|---|---|---|---|
| *Jumellea bathiei* Schltr. | JQ905400 | Rakotoarivelo & al. 035 [TAN] | JQ905339 | Rakotoarivelo & al. 035 [TAN] | JQ905522 | Rakotoarivelo & al. 035 [TAN] |
| *Jumellea bosseri* Pailler | JQ905401 | Pailler 270 [REU] | JQ905340 | Pailler 270 [REU] | JQ905523 | Pailler 270 [REU] |
| *Jumellea brachycentra* Schltr. | JQ905402 | Rakotoarivelo & al. 241 [TAN] | JQ905341 | Rakotoarivelo & al. 241 [TAN] | JQ905524 | Rakotoarivelo & al. 241 [TAN] |
| *Jumellea brevifolia* H.Perrier | JQ905403 | Rakotoarivelo & al. 300 [TAN] | JQ905342 | Rakotoarivelo & al. 300 [TAN] | JQ905525 | Rakotoarivelo & al. 300 [TAN] |
| *Jumellea comorensis* (Rchb.f.) Schltr. | JQ905404 | Rakotoarivelo & al. 040 [REU] | JQ905343 | Rakotoarivelo & al. 040 [REU] | JQ905526 | Rakotoarivelo & al. 040 [REU] |
| *Jumellea confusa* (Schltr.) Schltr. | JQ905440 | Rakotoarivelo & al. 088 [TAN] | JQ905379 | Rakotoarivelo & al. 088 [TAN] | JQ905562 | Rakotoarivelo & al. 088 [TAN] |
| *Jumellea densefoliata* Senghas | JQ905405 | Rakotoarivelo & al. 109 [TAN] | JQ905344 | Rakotoarivelo & al. 109 [TAN] | JQ905527 | Rakotoarivelo & al. 109 [TAN] |
| *Jumellea divaricata* (Frapp. ex Cordem.) Schltr. | JQ905406 | Pailler 267 [REU] | JQ905345 | Pailler 267 [REU] | JQ905528 | Pailler 267 [REU] |
| *Jumellea exilis* (Cordem.) Schltr. | JQ905407 | Pailler 216 [REU] | JQ905346 | Pailler 216 [REU] | JQ905529 | Pailler 216 [REU] |
| *Jumellea fragrans* (Thouars) Schltr. | MH237206 | Micheneau & Pailler 10 [REU] | KF557918 | Micheneau & Pailler 10 [REU] | KF558224 | Micheneau & Pailler 10 [REU] |
| *Jumellea francoisii* Schltr. | JQ905411 | Rakotoarivelo & al. 230 [TAN] | JQ905350 | Rakotoarivelo & al. 230 [TAN] | JQ905533 | Rakotoarivelo & al. 230 [TAN] |
| *Jumellea hyalina* H.Perrier | JQ905415 | Rakotoarivelo & al. 006 [TAN] | JQ905354 | Rakotoarivelo & al. 006 [TAN] | JQ905537 | Rakotoarivelo & al. 006 [TAN] |
| *Jumellea ibityana* Schltr. | JQ905416 | Rakotoarivelo & al. 031 [TAN] | JQ905355 | Rakotoarivelo & al. 031 [TAN] | JQ905538 | Rakotoarivelo & al. 031 [TAN] |
| *Jumellea imerinensis* Schltr. | JQ905408 | Rakotoarivelo *et al.* 317 [TAN] | JQ905347 | Rakotoarivelo *et al.* 317 [TAN] | JQ905469 | Rakotoarivelo *et al.* 317 [TAN] |
| *Jumellea jumelleana* (Schltr.) Summerh. | JQ905417 | Rakotoarivelo & al. 099 [TAN] | JQ905356 | Rakotoarivelo & al. 099 [TAN] | JQ905539 | Rakotoarivelo & al. 099 [TAN] |
| *Jumellea lignosa* (Schltr.) Schltr. | JQ905418 | Rakotoarivelo & al. 036 [TAN] | JQ905357 | Rakotoarivelo & al. 036 [TAN] | JQ905540 | Rakotoarivelo & al. 036 [TAN] |
| *Jumellea linearipetala* H.Perrier | MH237226 | No voucher (CM6m) | KF557971 | No voucher (CM6m) | KF558103 | No voucher (CM6m) |
| *Jumellea longivaginans* | JQ905414 | Rakotoarivelo *et al.* 150 | JQ905362 | Rakotoarivelo *et al.* 150 | JQ905475 | Rakotoarivelo *et al.* 150 |

| | | | | | | |
|---|---|---|---|---|---|---|
| H.Perrier | | [TAN] | | [TAN] | | [TAN] |
| *Jumellea majalis* (Schltr.) Schltr. | JQ905427 | Rakotoarivelo & al. 307 [TAN] | JQ905366 | Rakotoarivelo & al. 307 [TAN] | JQ905549 | Rakotoarivelo & al. 307 [TAN] |
| *Jumellea major* Schltr. | JQ905424 | Rakotoarivelo *et al.* 322 [TAN] | JQ905363 | Rakotoarivelo *et al.* 322 [TAN] | JQ905485 | Rakotoarivelo *et al.* 322 [TAN] |
| *Jumellea maxillarioides* (Ridl.) Schltr. | DQ091757 | unvouchered | Farminhão *et al.* (2020) | 1T53 [TAN] | Farminhão *et al.* (2020) | 1T53 [TAN] |
| *Jumellea pachyceras* Schltr. | JQ905428 | Rakotoarivelo *et al.* 311 [TAN] | JQ905367 | Rakotoarivelo *et al.* 311 [TAN] | JQ905489 | Rakotoarivelo *et al.* 311 [TAN] |
| *Jumellea pailleri* F.Rakotoar. | JQ905429 | Rakotoarivelo & al. 060 [REU] | JQ905368 | Rakotoarivelo & al. 060 [REU] | JQ905551 | Rakotoarivelo & al. 060 [REU] |
| *Jumellea papangensis* H.Perrier | Farminhão *et al.* (2020) | 84A41 [TAN] | KF558022 | Chase 17913 [K] | KF558079 | Chase 17913 [K] |
| *Jumellea peyrotii* Bosser | JQ905430 | Rakotoarivelo *et al.* 144 [TAN] | JQ905369 | Rakotoarivelo *et al.* 144 [TAN] | JQ905491 | Rakotoarivelo *et al.* 144 [TAN] |
| *Jumellea punctata* H.Perrier | XX | 424A443 [] | XX | AMB4870 [] | XX | 424A443 [] |
| *Jumellea recta* (Thouars) Schltr. | JQ905434 | Pailler 205 [REU] | KF557928 | Fournel 114 [REU] | KF558141 | Fournel 114 [REU] |
| *Jumellea recurva* (Thouars) Schltr. | JQ905433 | Pailler 204 [REU] | JQ905372 | Pailler 204 [REU] | JQ905555 | Pailler 204 [REU] |
| *Jumellea rigida* Schltr. | JQ905437 | Rakotoarivelo & al. 220 [TAN] | JQ905376 | Rakotoarivelo & al. 220 [TAN] | JQ905559 | Rakotoarivelo & al. 220 [TAN] |
| *Jumellea rossii* Senghas | JQ905438 | Pailler 293 [REU] | JQ905377 | Pailler 293 [REU] | JQ905560 | Pailler 293 [REU] |
| *Jumellea similis* Schltr. | JQ905442 | Rakotoarivelo & al. 100 [TAN] | JQ905381 | Rakotoarivelo & al. 100 [TAN] | JQ905564 | Rakotoarivelo & al. 100 [TAN] |
| *Jumellea spathulata* (Ridl.) Schltr. | JQ905441 | Rakotoarivelo *et al.* 209 [TAN] | JQ905380 | Rakotoarivelo *et al.* 209 [TAN] | JQ905502 | Rakotoarivelo *et al.* 209 [TAN] |
| *Jumellea stenoglossa* H.Perrier | JQ905445 | Pailler 239 [TAN] | JQ905384 | Pailler 239 [TAN] | Farminhão *et al.* (2020) | 87T124 [TAN] |
| *Jumellea stenophylla* (Frapp. ex Cordem.) Schltr. | MH237241 | Micheneau & Pailler 92 [REU] | KF558026 | Micheneau & Pailler 92 [REU] | KF558033 | Micheneau & Pailler 92 [REU] |
| *Jumellea tenuibracteata* (H.Perrier ex Hermans) F.P.Rakotoar. & Pailler | JQ905421 | Rakotoarivelo & al. 321 [TAN] | JQ905360 | Rakotoarivelo & al. 321 [TAN] | JQ905543 | Rakotoarivelo & al. 321 [TAN] |

| | | | | | | |
|---|---|---|---|---|---|---|
| *Jumellea teretifolia* Schltr. | JQ905447 | Rakotoarivelo & al. 160 [TAN] | JQ905386 | Rakotoarivelo & al. 160 [TAN] | JQ905568 | Rakotoarivelo & al. 160 [TAN] |
| *Jumellea triquetra* (Thouars) Schltr. | XX | Micheneau & Pailler 11 [REU] | KF557924 | Micheneau & Pailler 11 [REU] | KF558216 | Micheneau & Pailler 11 [REU] |
| *Jumellea unguicularis* Schltr. | JQ905450 | Rakotoarivelo & al. 216 [TAN] | JQ905389 | Rakotoarivelo & al. 216 [TAN] | JQ905571 | Rakotoarivelo & al. 216 [TAN] |
| *Jumellea walleri* (Rolfe) la Croix | Farminhão *et al.* (2020) | No voucher (CM6) | KF557931 | No voucher (CM6) | KF558148 | No voucher (CM6) |
| *Jumellea zaratananae* Schltr. | JQ905449 | Rakotoarivelo & al. 326 [TAN] | JQ905388 | Rakotoarivelo & al. 326 [TAN] | JQ905570 | Rakotoarivelo & al. 326 [TAN] |
| *Kylicanthe cornuata* Descourv., Stévart & Droissart | MH237081 | YAS 2498 [BRLU] | MK68555 4 | YAS 2498 [BRLU] | Farminhão *et al.* (2020) | YAS 2498 [BRLU] |
| *Kylicanthe quintasii* (Rolfe) Farminhão, Stévart & Droissart | Farminhão *et al.* (2020) | Stévart 4708 [BRLU] | MK68562 6 | Bom Sucesso 1981.1 [BRLU] | Farminhão *et al.* (2020) | Stévart 4708 [BRLU] |
| *Lemurella pallidiflora* Bosser | DQ091591 | Kew accession 4958 [K] | DQ091330 | Kew accession 4958 [K] | — | — |
| *Listrostachys pertusa* (Lindl.) Rchb.f. | MH237083 | YAS 2522 [BRLU] | MK68555 6 | YAS 2522 [BRLU] | MK722001 | YAS 2522 [BRLU] |
| *Microcoelia aphylla* (Thouars) Summerh. | DQ091651 | Carlsward 341 (BRLU) | DQ091400 | Carlsward 341 (BRLU) | Farminhão *et al.* (2020) | Carlsward 341 (BRLU) |
| *Microcoelia bulbocalcarata* L.Jonss. | MH237154 | Bom Sucesso 917 [BRLU] | MK68562 4 | Bom Sucesso 917 [BRLU] | Farminhão *et al.* (2020) | Bom Sucesso 917 [BRLU] |
| *Microcoelia caespitosa* (Rolfe) Summerh. | — | — | Farminhão *et al.* (2020) | YAS 7220 [BRLU] | — | — |
| *Microcoelia koehleri* (Schltr.) Summerh. | — | — | Farminhão *et al.* (2020) | BR 19910194-71 | — | — |
| *Mystacidium aliceae* Bolus | MF980236 | Martos 775 [NU] | MG25022 7 | Martos 775 [NU] | MG250247 | Martos 775 [NU] |
| *Mystacidium brayboniae* Summerh. | DQ091572 | Carlsward 179 (FLAS) | DQ091361 | Carlsward 179 (FLAS) | DQ091486 | Carlsward 179 (FLAS) |
| *Mystacidium capense* (L.f.) Schltr. | MF980235 | Martos 769 [NU] | MG25022 6 | Martos 769 [NU] | Farminhão *et al.* (2020) | Whitten 1781 [FLAS] |
| *Mystacidium flanaganii* (Bolus) Bolus | MF980245 | Young 1326 [NU] | MG25023 4 | Young 1326 [NU] | MG250254 | Young 1326 [NU] |
| *Mystacidium* | MF980228 | Bytebier 2227 | MG25022 | Bytebier 2227 | MG250240 | Bytebier 2227 |

| | | | | | | |
|---|---|---|---|---|---|---|
| *gracile* Harv. | | [NU] | 1 | [NU] | | [NU] |
| *Mystacidium pusillum* Harv. | MF980233 | Martos 740 [NU] | MG25022 4 | Martos 740 [NU] | — | — |
| *Mystacidium venosum* Harv. ex Rolfe | MF980232 | Martos 733 [NU] | MG25022 3 | Martos 733 [NU] | Farminhão *et al.* (2020) | Hermans 5084 (K] |
| *Mystacidium tanganyikense* Summerh. | MF980224 | Ashton unvouchered | MG25021 9 | Ashton unvouchered | MG250236 | Ashton unvouchered |
| *Neobathiea grandidieriana* (Rchb.f.) Garay | DQ091589 | Carlsward 395 [FLAS] | DQ091329 | Carlsward 395 [FLAS] | Farminhão *et al.* (2020) | Carlsward 395 [FLAS] |
| *Nephrangis filiformis* (Kraenzl.) Summerh. | MH237086 | YAS 2916 [BRLU] | MK68555 9 | YAS 2916 [BRLU] | Farminhão *et al.* (2020) | SIB 1690 [BRLU] |
| *Oeonia rosea* Ridl. | DQ091586 | Whitten 1813 [FLAS] | DQ091328 | Whitten 1813 [FLAS] | Farminhão *et al.* (2020) | Whitten 1813 [FLAS] |
| *Oeoniella polystachys* (Thouars) Schltr. | DQ091736 | Carlsward 221 [FLAS] | DQ091432 | Carlsward 221 [FLAS] | Farminhão *et al.* (2020) | Carlsward 221 [FLAS] |
| *Phalaenopsis wilsonii* Rolfe | DQ091672 | Carlsward 331 [FLAS] | AB217751 | TBG144214 | — | — |
| *Planetangis longicaudata* (Rolfe) Stévart & Farminhão | MH237161 | BTO181 [BRLU] | — | — | MH237546 | BTO181 [BRLU] |
| *Plectrelminthus caudatus* (Lindl.) Summerh. | MH237090 | YAS 2803 [BRLU] | MK68556 3 | YAS 2803 [BRLU] | Farminhão *et al.* (2020) | YAS 2803 [BRLU] |
| *Podangis dactyloceras* (Rchb.f.) Schltr. | MH237089 | YAS 2652 [BRLU] | MK68556 2 | YAS 2652 [BRLU] | Farminhão *et al.* (2020) | YAS 2652 [BRLU] |
| *Podangis muscicola* (Rchb.f.) Farminhão & D'haijère | DQ091630 | Carlsward 169 [SEL] | DQ091387 | Carlsward 169 [SEL] | Farminhão *et al.* (2020) | Carlsward 169 [SEL] |
| *Podangis rhipsalisocia* (Rchb.f.) P.J.Cribb & Carlsward | MH237092 | YAS 2011 [BRLU] | MH68556 6 | YAS 2011 [BRLU] | Farminhão *et al.* (2020) | YAS 2011 [BRLU] |
| *Rhipidoglossum arbonnieri* (Geerinck) Eb.Fisch., Killmann, J.-P.Lebel & Delep. | MH237133 | See p. 374 in The Orchids of Rwanda [BRLU] | MK68561 0 | See p. 374 in The Orchids of Rwanda [BRLU] | MH237517 | See p. 374 in The Orchids of Rwanda [BRLU] |
| *Rhipidoglossum brachyceras* (Summerh.) Farminhão & Stévart | DQ091577 | Bytebier 361 [EA] | DQ091365 | Bytebier 361 [EA] | Farminhão *et al.* (2020) | Bytebier 361 [EA] |

| | | | | | | |
|---|---|---|---|---|---|---|
| *Rhipidoglossum brevifolium* Summerh. | Farminhão *et al.* (2020) | Farminhão *et al.* 95 [BRLU] | MK685662 | Stévart 2887 [BRLU] | Farminhão *et al.* (2020) | Farminhão *et al.* 95 [BRLU] |
| *Rhipidoglossum burttii* (Summerh.) Summerh. | Farminhão *et al.* (2020) | Farminhão & Dumbo 228 [BRLU] | MK685617 | Photo voucher (see p. 279 in The Orchids of Rwanda) | Farminhão *et al.* (2020) | Farminhão & Dumbo 228 [BRLU] |
| *Rhipidoglossum caffrum* (Bolus) Farminhão & Stévart | MF980234 | Martos 752 [NU] | MG250225 | Martos 752 [NU] | MG250245 | Martos 752 [NU] |
| *Rhipidoglossum confusum* (P.J.Cribb) Farminhão & Stévart | DQ091578 | Kew accession 3936 [K] | DQ091366 | Kew accession 3936 [K] | Farminhão *et al.* (2020) | Kew accession 3936 [K] |
| *Rhipidoglossum curvatum* (Rolfe) Garay | XX | YAS 5581 [BRLU] | XX | YAS 2334 [BRLU] | XX | YAS 1277 [BRLU] |
| *Rhipidoglossum delepierreanum* (J.-P.Lebel & Geerinck) Eb.Fisch., Killmann, J.-P.Lebel & Delep. | MH237005 | Photo voucher (see p. 377 in The Orchids of Rwanda) | MK685475 | Photo voucher (see p. 377 in The Orchids of Rwanda) | Farminhão *et al.* (2020) | Photo voucher (see p. 377 in The Orchids of Rwanda) |
| *Rhipidoglossum densiflorum* Summerh. | MH237099 | YAS 2154 [BRLU] | MK685573 | YAS 2154 [BRLU] | Farminhão *et al.* (2020) | YAS 2154 [BRLU] |
| *Rhipidoglossum eggelingii* (Summerh.) Farminhão & Stévart | Farminhão *et al.* (2020) | Farminhão & Dumbo 232 [BRLU] | MK685609 | Photo voucher (see pp. 372-373 in The Orchids of Rwanda | Farminhão *et al.* (2020) | Farminhão & Dumbo 253 [BRLU] |
| *Rhipidoglossum globulosocalcaratum* (De Wild.) Summerh. | MH237093 | YAS 2837 [BRLU] | MK685567 | YAS 2837 [BRLU] | Farminhão *et al.* (2020) | YAS 2837 [BRLU] |
| *Rhipidoglossum kamerunense* (Schltr.) Garay | Farminhão *et al.* (2020) | YAS 3723 [BRLU] | MK685646 | Nkonmeneck 300 [SEL] | Farminhão *et al.* (2020) | YAS 3723 [BRLU] |
| *Rhipidoglossum millarii* (Bolus) Farminhão & Stévart | DQ091579 | Carlsward 346 [FLAS] | MK685651 | Carlsward 346 [FLAS] | Farminhão *et al.* (2020) | Carlsward 346 [FLAS] |
| *Rhipidoglossum pendulum* (la Croix & P.J.Cribb) Farminhão & Stévart | MH237158 | Bom Sucesso 2155.1 [BRLU] | MK685627 | Bom Sucesso 2155.1 [BRLU] | Farminhão *et al.* (2020) | Bom Sucesso 2155.1 [BRLU] |
| *Rhipidoglossum polydactylum* (Kraenzl.) Garay | Farminhão *et al.* (2020) | Droissart & Stévart 699 [BRLU] | Farminhão *et al.* (2020) | Droissart & Stévart 699 [BRLU] | Farminhão *et al.* (2020) | Droissart & Stévart 699 [BRLU] |

| | | | | | | |
|---|---|---|---|---|---|---|
| *Rhipidoglossum pulchellum* var. *geniculatum* (Summerh.) Garay | MH237096 | YAS 3043 [BRLU] | MK685570 | YAS 3043 [BRLU] | Farminhão *et al.* (2020) | YAS 5423 [BRLU] |
| *Rhipidoglossum pusillum* (Summerh.) Farminhão & Stévart | Farminhão *et al.* (2020) | Photo voucher (see p. 97 in The Orchids of Rwanda) | MK685614 | Photo voucher (see p. 97 in The Orchids of Rwanda) | — | — |
| *Rhipidoglossum rutilum* (Rchb.f.) Schltr. | Farminhão *et al.* (2020) | YAS 2399 [BRLU] | MK685552 | YAS 2183 [BRLU] | Farminhão *et al.* (2020) | YAS 2399 [BRLU] |
| *Rhipidoglossum stolzii* (Schltr.) Garay | XX | BR5629 [BR] | — | — | XX | BR5629 [BR] |
| *Rhipidoglossum subsimplex* (Summerh.) Garay | MH237195 | Bytebier 546 [EA] | MK685653 | Bytebier 546 [EA] | Farminhão *et al.* (2020) | Bytebier 546 [EA] |
| *Rhipidoglossum thomense* (la Croix & P.J.Cribb) Farminhão & Stévart | MH237009 | Stévart 246 [BRLU] | MK685477 | Stévart 246 [BRLU] | Farminhão *et al.* (2020) | Stévart 246 [BRLU] |
| *Rhipidoglossum xanthopollinium* (Rchb.f.) Schltr. | MH237191 | Carlsward 384 [FLAS] | MK685649 | Carlsward 384 [FLAS] | Farminhão *et al.* (2020) | Carlsward 384 [FLAS] |
| *Solenangis clavata* (Rolfe) Schltr. | DQ091666 | Carlsward 397 [FLAS] | DQ091409 | Carlsward 397 [FLAS] | DQ091534 | Carlsward 397 [FLAS] |
| *Solenangis liberica* (Mansf.) R.Rice | — | — | — | — | XX | D'haijère 7 [BRLU] |
| *Solenangis saotomensis* R.Rice | XX | Stévart 274 [BRLU] | — | — | — | — |
| *Solenangis scandens* (Schltr.) Schltr. | Farminhão *et al.* (2020) | YAS 7294 [BRLU] | MK722038 | YAS 3308 [BRLU] | MK722019 | Yaoundé 3308 |
| *Solenangis impraedicta* sp. nov. | XX | RBR418 [BRLU] | XX | RBR418 [BRLU] | XX | RBR418 [BRLU] |
| *Solenangis wakefieldii* (Rolfe) P.J.Cribb & J.Stewart | DQ091667 | Bytebier 627 [EA] | DQ091410 | Bytebier 627 [EA] | DQ091535 | Bytebier 627 [EA] |
| *Sphyrarhynchus amaniensis* (Summerh.) Bytebier | DQ091568 | No voucher (B252) | DQ091357 | No voucher (B252) | Farminhão *et al.* (2020) | No voucher (B252) |
| *Sphyrarhynchus brevilobus* (Summerh.) Bytebier | DQ091569 | Bytebier 307 [EA] | DQ091358 | Bytebier 307 [EA] | Farminhão *et al.* (2020) | Bytebier 307 [EA] |
| *Sphyrarhynchus schliebenii* Mansf. | DQ091570 | Bytebier 393 [EA] | DQ091359 | Bytebier 393 [EA] | — | — |

| | | | | | | |
|---|---|---|---|---|---|---|
| *Summerhayesia laurentii* (De Wild.) P.J.Cribb | Farminhão *et al.* (2020) | BTO 220 | Farminhão *et al.* (2020) | YAS 7965 [BRLU] | Farminhão *et al.* (2020) | BTO 220 |
| *Tridactyle anthomaniaca* (Rchb.f.) Summerh. | MH236990 | Y3679 RH (cultivated at YAS) | MK685470 | YAS 2462 [BRLU] | MK721977 | Y3679 RH (cultivated at YAS) |
| *Tridactyle aurantiopunctata* P.J.Cribb & Stévart | KF672201 | Stévart 656 [BRLU] | MK685498 | Stévart 290 [BRLU] | Farminhão *et al.* (2020) | Stévart 290 [BRLU] |
| *Tridactyle bicaudata* (Lindl.) Schltr. | Farminhão *et al.* (2020) | Farminhão *et al.* 196 [BRLU] | KF672263 | KIS 135 (cultivated at Kisantu shade house) | Farminhão *et al.* (2020) | Bytebier 348 [EA] |
| *Tridactyle brevicalcarata* Summerh. | MH236967 | BR 20090424-75 | MK685636 | BTO 168 [BRLU] | Farminhão *et al.* (2020) | BTO 168 [BRLU] |
| *Tridactyle clavata* (Summerh.) R.Rice | MH237147 | Photo voucher (see p. 201 in The Orchids of Rwanda) | MK685620 | Photo voucher (see p. 201 in The Orchids of Rwanda) | Farminhão *et al.* (2020) | Photo voucher (see p. 201 in The Orchids of Rwanda) |
| *Tridactyle crassifolia* Summerh. | MH237176 | Nkongmeneck 2076 [SEL] | MK685642 | Nkongmeneck 2076 [SEL] | MK721976 | Nkongmeneck 2076 [SEL] |
| *Tridactyle exellii* P.J.Cribb & Stévart | MH237026 | Stévart 297 [BRLU] | MK685493 | Stévart 297 [BRLU] | MK721985 | Stévart 297 [BRLU] |
| *Tridactyle filifolia* (Schltr.) Schltr. | MH237196 | Bytebier 707 [EA] | MK685654 | Bytebier 707 [EA] | MK721989 | Bytebier 707 [EA] |
| *Tridactyle gentilii* (De Wild.) Schltr. | MH237101 | KIS 116 (cultivated at Kisantu) | MK685577 | KIS 116 (cultivated at Kisantu) | Farminhão *et al.* (2020) | KIS 116 (cultivated at Kisantu) |
| *Tridactyle latifolia* Summerh. | MH237024 | Primo and Stevart 94 [BRLU] | MK685491 | Primo and Stevart 94 [BRLU] | MH237398 | Primo and Stevart 94 [BRLU] |
| *Tridactyle laurentii* (De Wild.) Schltr. | MH236995 | YAS 2489 [BRLU] | MK685453 | No voucher | MK721996 | No voucher |
| *Tridactyle ligulifolia* (Summerh.) R.Rice | MH237130 | Photo voucher (see p. 202 in The Orchids of Rwanda) | MK685607 | Photo voucher (see p. 202 in The Orchids of Rwanda) | Farminhão *et al.* (2020) | Photo voucher (see p. 202 in The Orchids of Rwanda) |
| *Tridactyle minutifolia* Stévart & D'haijère | MH236984 | G.Brice 252 [BRLU] | MK685480 | Stévart 3609 [BRLU] | Farminhão *et al.* (2020) | G.Brice 252 [BRLU] |
| *Tridactyle muriculata* (Rendle) Schltr. | MH236996 | YAS 2189 [BRLU] | MK685468 | YAS 2189 [BRLU] | MK721974 | YAS 2189 [BRLU] |
| *Tridactyle nalaensis* (De Wild.) Schltr. | MH236971 | BR 20090372-23 | MK685634 | BT091 [BRLU] | — | — |
| *Tridactyle scottellii* (Rendle) Schltr. | Farminhão *et al.* (2020) | Photo voucher (BTO 231) | Farminhão *et al.* (2020) | Photo voucher (BTO 231) | Farminhão *et al.* (2020) | Ndong Bokung *et al.* 337 [BRLU] |

| | | | | | | |
|---|---|---|---|---|---|---|
| *Tridactyle thomensis* P.J.Cribb & Stévart | MH237034 | Stévart 1217 [BRLU] | MK722035 | Bom Sucesso 2040.1 | MK721983 | Bom Sucesso 2040.1 |
| *Tridactyle tridactylites* (Rolfe) Schltr. | MH237033 | Stévart 1229 [BRLU] | MK685501 | Stévart 1229 [BRLU] | Farminhão *et al.* (2020) | Stévart 1229 [BRLU] |
| *Tridactyle truncatiloba* Summerh. | MH237035 | Stévart *et al.* 1624 [BRLU] | MK685504 | Stévart *et al.* 1624 [BRLU] | Farminhão *et al.* (2020) | Stévart *et al.* 1679 [BRLU] |
| *Vanda falcata* (Thunb.) Beer | DQ091684 | Carlsward 163 [SEL] | DQ091318 | Carlsward 163 [SEL] | — | — |
| *Ypsilopus amaniensis* (Kraenzl.) D'haijère & Stévart | DQ091634 | Bytebier & Kirika 26 [EA] | DQ091386 | Bytebier & Kirika 26 [EA] | MK721981 | Bytebier & Kirika 26 [EA] |
| *Ypsilopus erectus* (P.J.Cribb) P.J.Cribb & J.Stewart | MK714122 | Grieve 1244 [EA] | MK722036 | Grieve 1244 [EA] | MK721991 | Grieve 1244 [EA] |
| *Ypsilopus furcistipes* (Summerh.) D'haijère & Stévart | DQ091635 | Bytebier 1731 [EA] | DQ091392 | Bytebier 1731 [EA] | MK721994 | Bytebier 1731 [EA] |
| *Ypsilopus longifolius* (Kraenzl.) Summerh. | DQ091636 | Bytebier 609 [EA] | DQ091393 | Bytebier 609 [EA] | Farminhão *et al.* (2020) | Bytebier 609 [EA] |
| *Ypsilopus schliebenii* (Mansf.) D'haijère & Stévart | MH236965 | BR 20090389-40 | MK685438 | BR 20090389-40 | — | — |
| *Ypsilopus tanneri* (P.J.Cribb) D'haijère & Stévart | DQ091632 | PCP 198 [EA] | DQ091394 | PCP 198 [EA] | MK721993 | PCP 198 [EA] |
| *Ypsilopus tricuspis* (Bolus) D'haijère & Stévart | XX | Farminhão 306 [BRLU] | XX | Farminhão 306 [BRLU] | XX | Farminhão 306 [BRLU] |
| *Ypsilopus viridiflorus* P.J.Cribb & J.Stewart | DQ091633 | Bytebier 402 [EA] | DQ091395 | Bytebier 402 [EA] | — | — |

| Taxon | *rps16* | | *trnC-petN* | | *ycf1* | |
|---|---|---|---|---|---|---|
| | Genbank accession | Voucher [Herbarium] | Genbank accession | Voucher [Herbarium] | Genbank accession | Voucher [Herbarium] |
| *Aerangis arachnopus* (Rchb.f.) Schltr. | Farminhão *et al.* (2020) | YAS 2436 [BRLU] | MH237430 | YAS 2436 [BRLU] | Farminhão *et al.* (2020) | YAS 2436 [BRLU] |
| *Aerangis articulata* (Rchb.f.) Schltr. | — | — | — | — | — | — |
| *Aerangis biloba* (Lindl.) Schltr. | — | — | MH237432 | YAS 2279 [BRLU] | — | — |

| | | | | | | |
|---|---|---|---|---|---|---|
| *Aerangis bouarensis* Chiron | XX | YAS 4286 [BRLU] | — | — | — | — |
| *Aerangis brachycarpa* (A.Rich.) T.Durand & Schinz | — | — | — | — | — | — |
| *Aerangis calantha* (Schltr.) Schltr. | — | — | MH237420 | YAS 2828 [BRLU] | — | — |
| *Aerangis citrata* (Thouars) Schltr. | — | — | DQ091461 | Whitten 1788 [FLAS] | EU490715 | Whitten 1788 [FLAS] |
| *Aerangis collum-cygni* Summerh. | Farminhão *et al.* (2020) | YAS 2779 [BRLU] | MH237433 | YAS 2779 [BRLU] | Farminhão *et al.* (2020) | YAS 2779 [BRLU] |
| *Aerangis confusa* J.Stewart | XX | Bytebier s.n. [EA] | — | — | XX | Bytebier s.n. [EA] |
| *Aerangis coriacea* Summerh. | XX | Bytebier 562 [EA] | — | — | XX | Bytebier 562 [EA] |
| *Aerangis ellisii* (B.S.Williams) Schltr. | KF558043 | Chase 15080 [K] | — | — | — | — |
| *Aerangis fastuosa* (Rchb.f.) Schltr. | XX | Carlsward 402 [FLAS] | — | — | XX | Carlsward 402 [FLAS] |
| *Aerangis flexuosa* (Ridl.) Schltr. | — | — | — | — | — | — |
| *Aerangis gracillima* (Kraenzl.) J.C.Arends & J.Stewart | Farminhão *et al.* (2020) | YAS 1404 [BRLU] | MH237312 | YAS 1404 [BRLU] | Farminhão *et al.* (2020) | YAS 2302 [BRLU] |
| *Aerangis gravenreuthii* (Kraenzl.) Schltr. | Farminhão *et al.* (2020) | YAS 1565 [BRLU] | MH237310 | YAS 1565 [BRLU] | Farminhão *et al.* (2020) | YAS 1565 [BRLU] |
| *Aerangis hariotiana* (Kraenzl.) P.J.Cribb & Carlsward | Farminhão *et al.* (2020) | Carlsward 292 [FLAS] | Farminhão *et al.* (2020) | Carlsward 292 [FLAS] | Farminhão *et al.* (2020) | Carlsward 292 [FLAS] |
| *Aerangis hildebrandtii* (Rchb.f.) P.J.Cribb & Carlsward | — | Kew accession 2616 [K] | DQ091468 | Kew accession 2616 [K] | Farminhão *et al.* (2020) | — |
| *Aerangis kirkii* (Rchb.f.) Schltr. | — | Bytebier 637 [EA] | DQ091457 | Bytebier 637 [EA] | — | — |
| *Aerangis kotschyana* (Rchb.f.) Schltr. | — | Bytebier 671 [EA] | DQ091459 | Bytebier 671 [EA] | — | — |
| *Aerangis luteoalba* var. *rhodosticta* (Kraenzl.) J.Stewart | — | — | DQ091471 | Bytebier 691 [EA] | — | — |
| *Aerangis macrocentra* (Schltr.) Schltr. | KF558085 | Hermans 779 [K] | — | — | — | — |
| *Aerangis modesta* | — | — | DQ09146 | Carlsward 242 | — | — |

| | | | | | | |
|---|---|---|---|---|---|---|
| (Hook.f.) Schltr. | | | 5 | [FLAS] | | |
| *Aerangis monantha* Schltr. | — | — | — | — | XX | 618A234bis/1 [] |
| *Aerangis punctata* J.Stewart | Farminhão *et al.* (2020) | No voucher (CM9) | KF558209 | No voucher (CM9) | Farminhão *et al.* (2020) | No voucher (CM9) |
| *Aerangis somalensis* (Schltr.) Schltr. | — | — | DQ09147 0 | Bytebier 1549 [EA] | — | — |
| *Aerangis stelligera* Summerh. | — | — | — | — | — | — |
| *Aerangis thomsonii* (Rolfe) Schltr. | — | — | — | — | XX | Kirika 968 [EA] |
| *Aerangis ugandensis* Summerh. | — | — | MH23750 7 | Delepierre & Lebel 194 [BR] | — | — |
| *Aerangis verdickii* (De Wild.) Schltr. | XX | See p. 93 in The Orchids of Rwanda [BRLU] | — | — | — | — |
| *Aeranthes arachnites* (Thouars) Lindl. | Farminhão *et al.* (2020) | Fournel 126 [REU] | KF558232 | Fournel 126 [REU] | Farminhão *et al.* (2020) | Fournel 126 [REU] |
| *Aeranthes ecalcarata* H.Perrier | XX | 826A1362 [] | XX | 826A1362 [] | XX | 826A1362 [] |
| *Aeranthes grandiflora* Lindl. | Farminhão *et al.* (2020) | ANT18 | Farminhão *et al.* (2020) | ANT18 | Farminhão *et al.* (2020) | ANT18 |
| *Aeranthes leandriana* Bosser | Farminhão *et al.* (2020) | 13T135 [BRLU] | Farminhão *et al.* (2020) | 13T135 [BRLU] | Farminhão *et al.* (2020) | 13T135 [BRLU] |
| *Aeranthes neoperrieri* Toill.-Gen., Ursch & Bosser | Farminhão *et al.* (2020) | No voucher (CM6) | KF558199 | No voucher (CM6) | Farminhão *et al.* (2020) | No voucher (CM6) |
| *Aeranthes nidus* Schltr. | Farminhão *et al.* (2020) | ANT14 | Farminhão *et al.* (2020) | ANT14 | Farminhão *et al.* (2020) | ANT14 |
| *Aeranthes peyrotii* Bosser | Farminhão *et al.* (2020) | Chase 14644 [K] | KF558197 | Chase 14644 [K] | — | — |
| *Aeranthes tenella* Bosser | Farminhão *et al.* (2020) | Pailler 137 [REU] | KF558162 | Pailler 137 [REU] | — | — |
| *Aerides odorata* Lour. | — | — | KF557954 | Chase 15081 [K] | — | — |
| *Afropectinariella atlantica* (Droissart & Stévart) M.Simo & Stévart | KF672312 | JB Bambusa 130 [BRLU] | KF662343 | Van der Laan 1068 [BRLU] | KF672326 | JB Bambusa 130 [BRLU] |

| | | | | | | |
|---|---|---|---|---|---|---|
| *Afropectinariella doratophylla* (Summerh.) M.Simo & Stévart | KF672297 | JB Bom Sucesso 1962.1 [BRLU] | MH237287 | Stévart 441 [BRLU] | KF672318 | JB Bom Sucesso 1962.1 [BRLU] |
| *Afropectinariella gabonensis* (Summerh.) M.Simo & Stévart | Farminhão *et al.* (2020) | MBG 542 [BRLU] | MH237297 | YAS 1109 [BRLU] | Farminhão *et al.* (2020) | YAS 1109 [BRLU] |
| *Afropectinariella pungens* (Schltr.) M.Simo & Stévart | KF672304 | YAS 2817 [BRLU] | MH237325 | BR 20090383-34 [BRLU] | KF672338 | YAS 2817 [BRLU] |
| *Afropectinariella subulata* (Lindl.) M.Simo & Stévart | KF672314 | YAS 2503 [BRLU] | KF672285 | BR 20090388-39 | KF672324 | BR 20090388-39 |
| *Amesiella philippinensis* (Ames) Garay | — | — | — | — | — | — |
| *Ancistrorhynchus capitatus* (Lindl.) Summerh. | Farminhão *et al.* (2020) | Stévart & Pial 956 [BRLU] | MH237422 | Stévart & Pial 956 [BRLU] | Farminhão *et al.* (2020) | YAS 1057 [BRLU] |
| *Ancistrorhynchus cephalotes* (Rchb.f.) Summerh. | Farminhão *et al.* (2020) | Nimba 198 [BRLU] | Farminhão *et al.* (2020) | Nimba 198 [BRLU] | Farminhão *et al.* (2020) | Nimba 198 [BRLU] |
| *Ancistrorhynchus clandestinus* (Lindl.) Schltr. | Farminhão *et al.* (2020) | YAS 2831 [BRLU] | MH237440 | YAS 2831 [BRLU] | Farminhão *et al.* (2020) | YAS 2745 [BRLU] |
| *Ancistrorhynchus crystalensis* P.J.Cribb & Laan | Farminhão *et al.* (2020) | Tchimbélé 18 [BRLU] | MH237493 | Tchimbélé 18 [BRLU] | Farminhão *et al.* (2020) | Tchimbélé 18 [BRLU] |
| *Ancistrorhynchus metteniae* (Kraenzl.) Summerh. | Farminhão *et al.* (2020) | YAS 2214 [BRLU] | MH237504 | Nimba 203 [BRLU] | Farminhão *et al.* (2020) | YAS 2214 [BRLU] |
| *Ancistrorhynchus ovatus* Summerh. | Farminhão *et al.* (2020) | BR 20090367-18 | MH237341 | BR 20090367-18 | Farminhão *et al.* (2020) | BR 20090367-18 |
| *Ancistrorhynchus recurvus* Finet | Farminhão *et al.* (2020) | YAS 2785 [BRLU] | MH237301 | No voucher | Farminhão *et al.* (2020) | YAS 2785 [BRLU] |
| *Ancistrorhynchus refractus* (Kraenzl.) Summerh. | Farminhão *et al.* (2020) | BR 20090460-14 | MH237332 | BR 20090460-14 | Farminhão *et al.* (2020) | BR 20090460-14 |
| *Ancistrorhynchus schumannii* (Kraenzl.) Summerh. | Farminhão *et al.* (2020) | YAS 895 [BRLU] | MH237304 | YAS 895 [BRLU] | Farminhão *et al.* (2020) | YAS 895 [BRLU] |
| *Ancistrorhynchus serratus* Summerh. | Farminhão *et al.* (2020) | YAS 2381 [BRLU] | MH237445 | YAS 2381 [BRLU] | — | — |
| *Ancistrorhynchus straussii* (Schltr.) Schltr. | Farminhão *et al.* (2020) | BR20090369-20 | MH237335 | BR20090369-20 | Farminhão *et al.* (2020) | BR20090369-20 |

| | | | | | | |
|---|---|---|---|---|---|---|
| *Ancistrorhynchus tenuicaulis* Summerh. | Farminhão *et al.* (2020) | See p. 95 in The Orchids of Rwanda | MH23750 9 | See p. 95 in The Orchids of Rwanda | Farminhão *et al.* (2020) | See p. 95 in The Orchids of Rwanda |
| *Angraecopsis elliptica* Summerh. | Farminhão *et al.* (2020) | YAS 4473 [BRLU] | Farminhão *et al.* (2020) | YAS 4473 [BRLU] | Farminhão *et al.* (2020) | Stévart 5075 [BRLU] |
| *Angraecopsis gracillima* (Rolfe) Summerh. | Farminhão *et al.* (2020) | Farminhão 231 [BRLU] | Farminhão *et al.* (2020) | Farminhão 231 [BRLU] | Farminhão *et al.* (2020) | Farminhão 231 [BRLU] |
| *Angraecopsis ischnopus* (Schltr.) Schltr. | Farminhão *et al.* (2020) | Droissart 689 [BRLU] | Farminhão *et al.* (2020) | Droissart 689 [BRLU] | Farminhão *et al.* (2020) | Droissart 689 [BRLU] |
| *Angraecopsis parviflora* (Thouars) Schltr. | — | — | Farminhão *et al.* (2020) | YAS 3830 [BRLU] | Farminhão *et al.* (2020) | YAS 7877 [BRLU] |
| *Angraecopsis* sp. nov. Oku | — | — | — | — | — | — |
| *Angraecopsis tenerrima* Kraenzl. | — | — | — | — | MG250268 | No voucher |
| *Angraecopsis tridens* (Lindl.) Schltr. | XX | Droissart 693 [BRLU] | XX | Droissart 693 [BRLU] | XX | Droissart 693 [BRLU] |
| *Angraecum acutipetalum* Schltr. | Farminhão *et al.* (2020) | 14T21 [TAN] | Farminhão *et al.* (2020) | 14T21 [TAN] | Farminhão *et al.* (2020) | 14T21 [TAN] |
| *Angraecum amplexicaule* Toill.-Gen. & Bosser | KT826861 | Andriananjamana ntsoa & al. 00203468 [MT] | — | — | — | — |
| *Angraecum appendiculatum* Frapp. ex Cordem. | Farminhão *et al.* (2020) | Pailler 107 [REU] | KF558173 | Pailler 107 [REU] | Farminhão *et al.* (2020) | Pailler 107 [REU] |
| *Angraecum arachnites* Schltr. | — | — | KF558250 | Hermans 4241 [K] | — | — |
| *Angraecum borbonicum* Bosser | Farminhão *et al.* (2020) | Micheneau 1 [BRLU] | KF558268 | Micheneau 1 [BRLU] | Farminhão *et al.* (2020) | Micheneau 1 [BRLU] |
| *Angraecum bracteosum* Balf.f. & S.Moore | KF558045 | Micheneau 7 [REU] | — | — | XX | Micheneau 7 [REU] |
| *Angraecum breve* Schltr. | — | — | — | — | XX | AMB5173 [] |
| *Angraecum cadetii* Bosser | Farminhão *et al.* (2020) | Pailler 157 [REU] | KF55822 | Pailler 157 [REU] | Farminhão *et al.* (2020) | Pailler 157 [REU] |
| *Angraecum* cf. *elephantinum* | XX | AMB5135 | — | — | XX | AMB5135 |
| *Angraecum chaetopodum* Schltr. | KT826856 | Andriananjamana ntsoa & al. 00203462 [MT] | — | — | — | — |
| *Angraecum* | Farminhão | 42T60 [TAN] | Farminhão | 42T60 [TAN] | Farminhão *et* | 42T60 [TAN] |

| | | | | | | |
|---|---|---|---|---|---|---|
| *clavigerum* Ridl. | *et al.* (2020) | | *et al.* (2020) | | *al.* (2020) | |
| *Angraecum compactum* Schltr. | KT826866 | Andriananjamanantsoa & al. 00203420 [MT] | — | — | — | — |
| *Angraecum conchiferum* Lindl. | XX | Bytebier 616 [EA] | — | — | — | — |
| *Angraecum cornigerum* Cordem. | Farminhão *et al.* (2020) | Pailler 176 [REU] | KF558247 | Pailler 176 [REU] | Farminhão *et al.* (2020) | Pailler 176 [REU] |
| *Angraecum corrugatum* (Cordem.) Micheneau | Farminhão *et al.* (2020) | Pailler 106 [REU] | KF558194 | Pailler 106 [REU] | Farminhão *et al.* (2020) | Pailler 106 [REU] |
| *Angraecum costatum* Frapp. ex Cordem. | Farminhão *et al.* (2020) | Pailler 174 [REU] | KF558180 | Pailler 174 [REU] | Farminhão *et al.* (2020) | Pailler 174 [REU] |
| *Angraecum cucullatum* Thouars | KF558051 | Pailler 108 [REU] | — | — | XX | Pailler 108 [REU] |
| *Angraecum didieri* (Baill. ex Finet) Schltr. | XX | AMB741.1 [] | XX | AMB5505 [] | XX | AMB741.1 [] |
| *Angraecum dives* Rolfe | Farminhão *et al.* (2020) | Marimoto 42 [EA] | DQ091547 | Marimoto 42 [EA] | — | — |
| *Angraecum eburneum* subsp. eburneum Bory | KF558128 | unvouchered [K] | XX | unvouchered [K] | XX | unvouchered [K] |
| *Angraecum eburneum* subsp. *superbum* (Thouars) H.Perrier | KF558104 | Carter 761 [K] | XX | Carter 761 [K] | XX | Carter 761 [K] |
| *Angraecum equitans* Schltr. | Farminhão *et al.* (2020) | Verlynde 52 [BRLU] | Farminhão *et al.* (2020) | Verlynde 52 [BRLU] | Farminhão *et al.* (2020) | Verlynde 52 [BRLU] |
| *Angraecum expansum* Thouars | Farminhão *et al.* (2020) | Micheneau 2 [BRLU] | KF558151 | Micheneau 2 [BRLU] | Farminhão *et al.* (2020) | Micheneau 2 [BRLU] |
| *Angraecum filicornu* Thouars | KT826892 | Andriananjamanantsoa & al. 00203460 [MT] | — | — | — | — |
| *Angraecum florulentum* Rchb.f. | KF558088 | unvouchered [K] | — | — | — | — |
| *Angraecum humblotianum* Schltr. | KF672255 | Simo M. 217 [BRLU] | KF672303 | Simo M. 217 [BRLU] | KF672321 | Simo M. 217 [BRLU] |
| *Angraecum huntleyoides* Schltr. | Farminhão *et al.* (2020) | Hermans 4248.1 [K] | KF558191 | Hermans 4248.1 [K] | Farminhão *et al.* (2020) | Hermans 4248.1 [K] |
| *Angraecum leonis* (Rchb.f.) André | Farminhão *et al.* | Carlsward 390 [FLAS] | DQ091551 | Carlsward 390 [FLAS] | KY558779 | Carlsward 390 [FLAS] |

| | | | | | | |
|---|---|---|---|---|---|---|
| | (2020) | | | | | |
| *Angraecum liliodorum* Frapp. ex Cordem. | Farminhão *et al.* (2020) | No voucher | KF558153 | No voucher | Farminhão *et al.* (2020) | No voucher |
| *Angraecum linearifolium* Garay | Farminhão *et al.* (2020) | 297T86 [BRLU] | Farminhão *et al.* (2020) | 297T86 [BRLU] | Farminhão *et al.* (2020) | 297T86 [BRLU] |
| *Angraecum longicalcar* (Bosser) Senghas | — | — | — | — | — | — |
| *Angraecum magdalenae* Schltr. & H.Perrier | Farminhão *et al.* (2020) | No voucher | KF558219 | No voucher | Farminhão *et al.* (2020) | No voucher |
| *Angraecum mauritianum* (Poir.) Frapp. | Farminhão *et al.* (2020) | No voucher | KF558195 | No voucher | Farminhão *et al.* (2020) | No voucher |
| *Angraecum moratii* Bosser | KT826840 | Andriananjamanantsoa & al. 00203447 [MT] | — | — | — | — |
| *Angraecum multiflorum* Thouars | KF558053 | Pailler 154 [REU] | XX | Pailler 154 [REU] | XX | Pailler 154 [REU] |
| *Angraecum nanum* Frapp. ex Cordem. | KT826868 | Andriananjamanantsoa & al. 00203325 [MT] | — | — | — | — |
| *Angraecum obesum* H.Perrier | JQ905454 | Rakotoarivelo & al. 008 [TAN] | — | — | JQ905279 | Rakotoarivelo & al. 008 [TAN] |
| *Angraecum oblongifolium* Toill.-Gen. & Bosser | KT826841 | Andriananjamanantsoa & al. 00203446 [MT] | — | — | — | — |
| *Angraecum obversifolium* Frapp. ex Cordem. | KF558125 | Micheneau & Pailler 8 [REU] | — | — | XX | Micheneau & Pailler 8 [REU] |
| *Angraecum panicifolium* H.Perrier | KF672307 | Simo 215 [BRLU] | MH237346 | Simo 215 [BRLU] | KF672322 | Simo 215 [BRLU] |
| *Angraecum pectinatum* Thouars | Farminhão *et al.* (2020) | No voucher (CM6h) | KF558154 | No voucher (CM6h) | Farminhão *et al.* (2020) | No voucher (CM6h) |
| *Angraecum penzigianum* Schltr. | KT826842 | Andriananjamanantsoa & al. 00203466 [MT] | — | — | — | — |
| *Angraecum protensum* Schltr. | XX | MO4617229 [NY] | — | — | — | — |
| *Angraecum pseudodidieri* H.Perrier | Farminhão *et al.* (2020) | No voucher (CM6) | KF558200 | No voucher (CM6) | Farminhão *et al.* (2020) | No voucher (CM6) |
| *Angraecum ramosum* Thouars | Farminhão *et al.* | Micheneau & Pailler 73 [REU] | KF558170 | Micheneau & Pailler 73 | Farminhão *et al.* (2020) | Micheneau & Pailler 73 |

| | | | | | | |
|---|---|---|---|---|---|---|
| | (2020) | | | [REU] | | [REU] |
| *Angraecum rutenbergianum* Kraenzl. | XX | Carlsward 300 [FLAS] | — | — | XX | Carlsward 300 [FLAS] |
| *Angraecum sacciferum* Lindl. | Farminhão *et al.* (2020) | Bytebier 2226 [NBG] | KF558181 | Bytebier 2226 [NBG] | Farminhão *et al.* (2020) | Bytebier 2226 [NBG] |
| *Angraecum sedifolium* Schltr. | XX | AMB5167 [] | XX | AMB5167 [] | — | — |
| *Angraecum serpens* (H.Perrier) Bosser | KT826843 | Andriananjamanantsoa & al. 00203438 [MT] | — | — | — | — |
| *Angraecum sesquipedale* Thouars | — | — | KF558179 | No voucher (CM6j) | — | — |
| *Angraecum sororium* Schltr. | KT826894 | Andriananjamanantsoa & al. 00203400 [MT] | — | — | — | — |
| *Angraecum* sp. | XX | AMB5175 [] | XX | AMB5175 [] | XX | AMB5175 [] |
| *Angraecum striatum* Thouars | KF558058 | Micheneau 4 [REU] | — | — | XX | Micheneau 4 [REU] |
| *Angraecum tenuifolium* Frapp. ex Cordem. | Farminhão *et al.* (2020) | Pailler 116 [REU] | KF558254 | Pailler 116 [REU] | Farminhão *et al.* (2020) | Pailler 116 [REU] |
| *Angraecum teretifolium* Ridl. | Farminhão *et al.* (2020) | No voucher (CM3f) | KF558190 | No voucher (CM3f) | Farminhão *et al.* (2020) | No voucher (CM3f) |
| *Angraecum triangulifolium* Senghas | KT826864 | Andriananjamanantsoa & al. 00203442 [MT] | — | — | — | — |
| *Angraecum viguieri* Schltr. | Farminhão *et al.* (2020) | Verlynde 51 [BRLU] | Farminhão *et al.* (2020) | Verlynde 51 [BRLU] | Farminhão *et al.* (2020) | Verlynde 51 [BRLU] |
| *Aziza trilobata* (Summerh.) Farminhão & D'haijère | — | — | — | — | Farminhão *et al.* (2020b) | Farminhão 11 [BRLU] |
| *Beclardia macrostachya* (Thouars) A.Rich. | — | — | Farminhão *et al.* (2020) | 27T13 | Farminhão *et al.* (2020) | 27T13 |
| *Bolusiella fractiflexa* Droissart, Stévart & Verlynde | Farminhão *et al.* (2020) | YAS 1357 [BRLU] | MH237306 | YAS 1357 [BRLU] | Farminhão *et al.* (2020) | YAS 1357 [BRLU] |
| *Bolusiella iridifolia* (Rolfe) Schltr. | Farminhão *et al.* (2020) | Bytebier 1113 [EA] | DQ091481 | Bytebier 1113 [EA] | Farminhão *et al.* (2020) | Photo voucher (see p. 106 in The Orchids of Rwanda) |
| *Bolusiella maudiae* (Bolus) Schltr. | Farminhão *et al.* (2020) | Bytebier 485 [EA] | DQ091480 | Bytebier 485 [EA] | Farminhão *et al.* (2020) | Bytebier 485 [EA] |

| | | | | | | |
|---|---|---|---|---|---|---|
| *Bolusiella talbotii* (Rendle) Summerh. | Farminhão *et al.* (2020) | YAS 2303 [BRLU] | MH237505 | Nimba 40 [BRLU] | Farminhão *et al.* (2020) | YAS 2303 [BRLU] |
| *Bolusiella zenkeri* (Kraenzl.) Schltr. | Farminhão *et al.* (2020) | YAS 2442 [BRLU] | MH237412 | YAS 2442 [BRLU] | Farminhão *et al.* (2020) | YAS 2442 [BRLU] |
| *Calyptrochilum aurantiacum* (P.J.Cribb & Laan) Stévart, M.Simo & Droissart | Farminhão *et al.* (2020) | YAS 2773 [BRLU] | MH237468 | YAS 2773 [BRLU] | Farminhão *et al.* (2020) | YAS 2773 [BRLU] |
| *Calyptrochilum christyanum* (Rchb.f.) Summerh. | — | — | MH237414 | YAS 1799 [BRLU] | Farminhão *et al.* (2020) | YAS 1799 [BRLU] |
| *Calyptrochilum emarginatum* (Afzel. ex Sw.) Schltr. | Farminhão *et al.* (2020) | YAS 2735 [BRLU] | MH237418 | YAS 2735 [BRLU] | Farminhão *et al.* (2020) | YAS 2735 [BRLU] |
| *Campylocentrum brenesii* Schltr. | XX | M. Blanco 2139 [USJ] | XX | M. Blanco 2139 [USJ] | KY558829 | M. Blanco 2139 [USJ] |
| *Campylocentrum fasciola* (Lindl.) Cogn. | — | — | DQ091445 | Carlsward 301 [FLAS] | KY558818 | Bogarín10415 [CR] |
| *Campylocentrum jamaicense* (Rchb.f. & Wullschl.) Benth. ex Fawc. | XX | Ackerman 3341 [UPRRP] | — | — | KY558801 | Ackerman 3341 [UPRRP] |
| *Campylocentrum minutum* C.Schweinf. | XX | MO4904973 [NY] | XX | MO4904973 [NY] | XX | MO4904973 [NY] |
| *Campylocentrum neglectum* (Rchb.f. & Warm.) Cogn. | — | — | AF506324 | Carlsward 272 [FLAS] | KY558804 | Carlsward 272 [FLAS] |
| *Campylocentrum pachyrrhizum* (Rchb.f.) Rolfe | — | — | AF506328 | Ackerman s.n. [UPRRP] | KY558825 | Ackerman s.n. [UPRRP] |
| *Campylocentrum poeppigii* (Rchb.f.) Rolfe | — | — | LT724158 | Bogarín 2218 [CR] | KY558810 | Bogarín 2218 [CR] |
| *Campylocentrum stenanthum* Schltr. | — | — | AY147227 | Carlsward 180 [FLAS] | KY558838 | Carlsward 180 [FLAS] |
| *Campylocentrum tyrridion* Garay & Dunst. ex Foldats | XX | Carnevali 5145 [FLAS] | XX | Carnevali 5145 [FLAS] | KY558820 | Carnevali 5145 [FLAS] |
| *Cleisostoma arietinum* (Rchb.f.) Garay | — | — | KJ733628 | Z.J.Liu 6991 [] | — | — |
| *Conchograecum affine* (Schltr.) Szlach., Grochocka, Oledrz. & Mytnik | Farminhão *et al.* (2020) | BTO 95 [BRLU] | MH237489 | BTO 95 [BRLU] | Farminhão *et al.* (2020) | BTO 95 [BRLU] |

| | | | | | | |
|---|---|---|---|---|---|---|
| *Conchograecum angustum* (Rolfe) Szlach., Grochocka, Oledrz. & Mytnik | Farminhão *et al.* (2020) | YAS 1226 [BRLU] | MH237279 | YAS 1226 [BRLU] | Farminhão *et al.* (2020) | YAS 1226 [BRLU] |
| *Conchograecum claessensii* (De Wild.) Szlach., Grochocka, Oledrz. & Mytnik | Farminhão *et al.* (2020) | YAS 3136 [BRLU] | MH237443 | YAS 3136 [BRLU] | Farminhão *et al.* (2020) | YAS 3136 [BRLU] |
| *Conchograecum cribbianum* (Szlach. & Olszewski) Szlach., Grochocka, Oledrz. & Mytnik | XX | MBG249 [BRLU] | XX | MBG249 [BRLU] | XX | MBG249 [BRLU] |
| *Conchograecum cultriforme* (Summerh.) Szlach., Grochocka, Oledrz. & Mytnik | Farminhão *et al.* (2020) | Carlsward 298 [FLAS] | AF506340 | Carlsward 298 [FLAS] | Farminhão *et al.* (2020) | Carlsward 298 [FLAS] |
| *Conchograecum erectum* (Summerh.) Szlach., Grochocka, Oledrz. & Mytnik | Farminhão *et al.* (2020) | Bytebier 801 [EA] | DQ091447 | Bytebier 801 [EA] | Farminhão *et al.* (2020) | Bytebier 801 [EA] |
| *Conchograecum geerinckianum* (Stévart & Jecmenica) Szlach., Grochocka, Oledrz. & Mytnik | XX | SIB762 [BRLU] | XX | SIB762 [BRLU] | XX | SIB762 [BRLU] |
| *Conchograecum lanceolatum* (Ječmenica, Stévart & Droissart) Szlach., Grochocka, Oledrz. & Mytnik | Farminhão *et al.* (2020) | BTO 23 [BRLU] | MH237548 | BTO 23 [BRLU] | Farminhão *et al.* (2020) | BTO 23 [BRLU] |
| *Conchograecum moandense* (De Wild.) Szlach., Grochocka, Oledrz. & Mytnik | Farminhão *et al.* (2020) | YAS 2836 [BRLU] | MH237275 | YAS 654 [BRLU] | Farminhão *et al.* (2020) | YAS 2836 [BRLU] |
| *Conchograecum multinominatum* (Rendle) Szlach., Grochocka, Oledrz. & Mytnik | Farminhão *et al.* (2020) | YAS 696 [BRLU] | MH237281 | YAS 696 [BRLU] | Farminhão *et al.* (2020) | YAS 696 [BRLU] |
| *Conchograecum oliveirae* (Stévart & Jecmenica) Szlach., Grochocka, Oledrz. & Mytnik | Farminhão *et al.* (2020) | Bom Sucesso 1617 [BRLU] | Farminhão *et al.* (2020) | Bom Sucesso 1617 [BRLU] | Farminhão *et al.* (2020) | Bom Sucesso 1617 [BRLU] |
| *Conchograecum reygaertii* (De Wild.) Szlach., | Farminhão *et al.* (2020) | YAS 2829 [BRLU] | MH237451 | YAS 2829 [BRLU] | Farminhão *et al.* (2020) | YAS 2829 [BRLU] |

| | | | | | | |
|---|---|---|---|---|---|---|
| Grochocka, Oledrz. & Mytnik | | | | | | |
| *Conchograecum sanfordii* (P.J.Cribb & B.J.Pollard) Szlach., Grochocka, Oledrz. & Mytnik | Farminhão *et al.* (2020) | YAS 1136 [BRLU] | MH237451 | YAS 1136 [BRLU] | Farminhão *et al.* (2020) | YAS 1136 [BRLU] |
| *Conchograecum umbrosum* (P.J.Cribb) Szlach., Grochocka, Oledrz. & Mytnik | — | — | — | — | — | — |
| *Cryptopus paniculatus* H.Perrier | Farminhão *et al.* (2020) | 2073A4951 [TAN] | KF558264 | Hermans 5392 [K] | Farminhão *et al.* (2020) | 2073A4951 [TAN] |
| *Cyrtorchis arcuata* (Lindl.) Schltr. | Farminhão *et al.* (2020) | BR 20090398-49 | MH237326 | BR 20090398-49 | Farminhão *et al.* (2020) | BR 20090398-49 |
| *Cyrtorchis arcuata* subsp. *whytei* (Rolfe) Summerh. | Farminhão *et al.* (2020) | YAS 3599 [BRLU] | — | — | Farminhão *et al.* (2020) | YAS 3599 [BRLU] |
| *Cyrtorchis aschersonii* (Kraenzl.) Schltr. | Farminhão *et al.* (2020) | YAS 2434 [BRLU] | MH237481 | YAS 2434 [BRLU] | Farminhão *et al.* (2020) | YAS 2434 [BRLU] |
| *Cyrtorchis chailluana* (Hook.f.) Schltr. | MK697535 | BR 19750114 | DQ091506 | Carlsward 156 [SEL] | MK722018 | Carlsward 156 [SEL] |
| *Cyrtorchis hamata* (Rolfe) Schltr. | Farminhão *et al.* (2020) | Nimba 222 | Farminhão *et al.* (2020) | YAS 4671 [BRLU] | Farminhão *et al.* (2020) | YAS 4671 [BRLU] |
| *Cyrtorchis henriquesiana* (Ridl.) Schltr. | Azandi *et al.* (in press) | BTO 99 [BRLU] | Azandi *et al.* (in press) | BTO 99 [BRLU] | Azandi *et al.* (in press) | BTO 99 [BRLU] |
| *Cyrtorchis letouzeyi* Szlach. & Olszewski | Azandi *et al.* (in press) | YAS 253 [BRLU] | Azandi *et al.* (in press) | YAS 253 [BRLU] | Azandi *et al.* (in press) | YAS 253 [BRLU] |
| *Cyrtorchis monteiroae* (Rchb.f.) Schltr. | Farminhão *et al.* (2020) | Bom Sucesso 2142.1 [BRLU] | MH237539 | Bom Sucesso 2142.1 [BRLU] | Farminhão *et al.* (2020) | Bom Sucesso 2142.1 [BRLU] |
| *Cyrtorchis praetermissa* Summerh. | Farminhão *et al.* (2020) | Photo voucher (see pp. 168-169 in The Orchids of Rwanda | MH237519 | Photo voucher (see pp. 168-169 in The Orchids of Rwanda | Farminhão *et al.* (2020) | No voucher (Carlsward *et al.*) |
| *Cyrtorchis ringens* (Rchb.f.) Summerh. | Farminhão *et al.* (2020) | Photo voucher (see pp. 166-167 in The Orchids of Rwanda | MH237521 | Photo voucher (see pp. 166-167 in The Orchids of Rwanda | Farminhão *et al.* (2020) | Photo voucher (see pp. 166-167 in The Orchids of Rwanda |
| *Dendrophylax alcoa* Dod | — | — | — | — | — | — |
| *Dendrophylax* | — | — | — | — | JN176133 | Carlsward 199 |

| | | | | | | |
|---|---|---|---|---|---|---|
| *barrettiae* Fawc. & Rendle | | | | | | [FLAS] |
| *Dendrophylax fawcettii* Rolfe | — | — | — | — | JN176135 | Whitten 3265 [FLAS] |
| *Dendrophylax filiformis* (Sw.) Benth. ex Fawc. | — | — | AF506323 | Whitten 1842 [FLAS] | — | — |
| *Dendrophylax funalis* (Sw.) Benth. ex Rolfe | — | — | KF558238 | Carlsward 302 [FLAS] | KY558782 | Carlsward 302 [FLAS] |
| *Dendrophylax lindenii* (Lindl.) Benth. ex Rolfe | — | — | — | — | JN176130 | photo voucher [FLAS] |
| *Dendrophylax megarhizus* Molgo & Carnevali | — | — | — | — | JN176112 | Carnevali 5907 [CICY] |
| *Dendrophylax porrectus* (Rchb.f.) Carlsward & Whitten | — | — | AY147232 | Carlsward 329 [FLAS] | JN176123 | Carlsward 329 [FLAS] |
| *Dendrophylax sallei* (Rchb.f.) Benth. ex Rolfe | — | — | — | — | JN176128 | Whitten 1945 [JBSD] |
| *Dendrophylax varius* (Aubl.) Urb. | — | — | — | — | JN176129 | Whitten 1960 [JBSD] |
| *Diaphananthe bidens* (Afzel. ex Sw.) Schltr. | — | — | Farminhão *et al.* (2020) | YAS 2340 [BRLU] | Farminhão *et al.* (2020) | YAS 2152 [BRLU] |
| *Diaphananthe fragrantissima* (Rchb.f.) Schltr. | Farminhão *et al.* (2020) | Kirika 536 [EA] | MH23745 5 | YAS 1862 [BRLU] | Farminhão *et al.* (2020) | Kirika 536 [EA] |
| *Diaphananthe ichneumonea* (Lindl.) P.J.Cribb & Carlsward | Farminhão *et al.* (2020) | Calrsward 286 [FLAS] | MH23731 5 | YAS 889 [BRLU] | Farminhão *et al.* (2020) | YAS 889 [BRLU] |
| *Diaphananthe lebelii* (Eb.Fisch. & Killmann) Descourv. & Stévart | Farminhão *et al.* (2020) | HOLOTYPE | MH23737 7 | HOLOTYPE | Farminhão *et al.* (2020) | HOLOTYPE |
| *Diaphananthe lecomtei* (Finet) P.J.Cribb & Carlsward | — | — | MH23741 9 | YAS 2336 [BRLU] | Farminhão *et al.* (2020) | YAS 2336 [BRLU] |
| *Diaphananthe lorifolia* Summerh. | Farminhão *et al.* (2020) | Bytebier 346 [EA] | DQ09150 1 | Bytebier 346 [EA] | Farminhão *et al.* (2020) | Bytebier 346 [EA] |
| *Diaphananthe odoratissima* (Rchb.f.) P.J.Cribb & Carlsward | Farminhão *et al.* (2020) | No voucher (Carlsward *et al.*) | MH23741 7 | YAS 1217 [BRLU] | Farminhão *et al.* (2020) | YAS 1217 [BRLU] |

| | | | | | | |
|---|---|---|---|---|---|---|
| *Diaphananthe pellucida* (Lindl.) Schltr. | — | — | Farminhão *et al.* (2020) | YAS 1317 [BRLU] | Farminhão *et al.* (2020) | BTO 171 [BRLU] |
| *Diaphananthe sarcophylla* (Schltr. ex Prain) P.J.Cribb & Carlsward | Farminhão *et al.* (2020) | Bytebier 339 [EA] | DQ091503 | Bytebier 339 [EA] | Farminhão *et al.* (2020) | Bytebier 339 [EA] |
| *Diaphananthe sarcorhynchoides* J.B.Hall | Farminhão *et al.* (2020) | MBG 61 [BRLU] | MH237494 | MBG 61 [BRLU] | Farminhão *et al.* (2020) | MBG 61 [BRLU] |
| *Diaphananthe spiralis* (Stévart & Droissart) P.J.Cribb & Carlsward | XX | YAS 1842 [BRLU] | — | — | XX | YAS 1842 [BRLU] |
| *Diaphananthe vesicata* (Lindl.) P.J.Cribb & Carlsward | Farminhão *et al.* (2020) | Chase 14645 [K] | MH237439 | YAS 312 [BRLU] | Farminhão *et al.* (2020) | Chase 14645 [K] |
| *Dolabrifolia aporoides* (Summerh.) Szlach. & Romowicz | Farminhão *et al.* (2020) | YAS 2246 [BRLU] | MH237340 | BR 20090370-21 | Farminhão *et al.* (2020) | BR 20090370-21 |
| *Dolabrifolia bancoensis* (Burg) Szlach. & Romowicz | KX060119 | YAS 2207 [BRLU] | MH237274 | YAS 2187 [BRLU] | KX060158 | YAS 2207 [BRLU] |
| *Dolabrifolia biteaui* (M.Simo & Stévart) M.Simo | Farminhão *et al.* (2020) | BTO 19 [BRLU] | Farminhão *et al.* (2020) | BTO 19 [BRLU] | KX060149 | BR20090387-38 |
| *Dolabrifolia disticha* (Lindl.) Szlach. & Romowicz | KX060120 | YAS 62 [BRLU] | KX060127 | YAS 2309 [BRLU] | KX060147 | YAS 2309 [BRLU] |
| *Dolabrifolia podochiloides* (Schltr.) Szlach. & Romowicz | KX060103 | Bambusa 159 [BRLU] | KX060123 | MBG 656 [BRLU] | KX060143 | MBG 656 [BRLU] |
| *Eichlerangraecum angustipetalum* com. ined. | XX | YAS 112 [BRLU] | XX | YAS 112 [BRLU] | XX | YAS 112 [BRLU] |
| *Eichlerangraecum birrimense* (Rolfe) Szlach., Mytnik & Grochocka | Farminhão *et al.* (2020) | YAS 399 [BRLU] | MH237450 | YAS 399 [BRLU] | Farminhão *et al.* (2020) | YAS 399 [BRLU] |
| *Eichlerangraecum eichlerianum* var. *eichlerianum* (Kraenzl.) Szlach., Mytnik & Grochocka | Farminhão *et al.* (2020b) | BTO170 [BRLU] | XX | BTO170 [BRLU] | Farminhão *et al.* (2020b) | BTO170 [BRLU] |
| *Eichlerangraecum eichlerianum* var. *curvicalcaratum* | XX | YAS 944 [BRLU] | XX | YAS 944 [BRLU] | XX | YAS 944 [BRLU] |

| | | | | | | |
|---|---|---|---|---|---|---|
| (Szlach. & Olszewski) Szlach., Mytnik & Grochocka | | | | | | |
| *Eichlerangraecum infundibulare* (Lindl.) Szlach., Mytnik & Grochocka | Farminhão *et al.* (2020) | No voucher (CM22961) | KF558243 | No voucher (CM22961) | Farminhão *et al.* (2020) | No voucher (CM22961) |
| *Erasanthe henrici* (Schltr.) P.J.Cribb, Hermans & D.L.Roberts | Farminhão *et al.* (2020) | No voucher (CM22946) | KF558175 | No voucher (CM22946) | Farminhão *et al.* (2020) | No voucher (CM22946) |
| *Eurychone galeandrae* (Rchb.f.) Schltr. | — | — | — | — | — | — |
| *Eurychone rothschildiana* (O'Brien) Schltr. | Farminhão *et al.* (2020) | Traoré 141 [BRLU] | MH237501 | Traoré 141 [BRLU] | Farminhão *et al.* (2020) | Traoré 141 [BRLU] |
| *Jumellea alionae* P.J.Cribb | JQ905505 | Rakotoarivelo & al. 200 [TAN] | — | — | JQ905322 | Rakotoarivelo & al. 200 [TAN] |
| *Jumellea ambongensis* Schltr. | JQ905451 | Rakotoarivelo & al. 131 [TAN] | — | — | JQ905276 | Rakotoarivelo & al. 131 [TAN] |
| *Jumellea amplifolia* Schltr. | Farminhão *et al.* (2020) | No voucher (CM6l) | KF558236 | No voucher (CM6l) | Farminhão *et al.* (2020) | No voucher (CM6l) |
| *Jumellea anjouanensis* (Finet) H.Perrier | — | — | JQ905553 | Rakotoarivelo *et al.* 203 [TAN] | JQ905311 | Rakotoarivelo *et al.* 203 [TAN] |
| *Jumellea arachnantha* (Rchb.f.) Schltr. | — | — | JQ905561 | Rakotoarivelo *et al.* 136 [TAN] | JQ905317 | Rakotoarivelo *et al.* 136 [TAN] |
| *Jumellea arborescens* H.Perrier | JQ905459 | Rakotoarivelo & al. 202 [TAN] | — | — | — | — |
| *Jumellea bathiei* Schltr. | JQ905461 | Rakotoarivelo & al. 035 [TAN] | — | — | JQ905283 | Rakotoarivelo & al. 035 [TAN] |
| *Jumellea bosseri* Pailler | JQ905462 | Pailler 270 [REU] | — | — | — | — |
| *Jumellea brachycentra* Schltr. | JQ905463 | Rakotoarivelo & al. 241 [TAN] | — | — | JQ905284 | Rakotoarivelo & al. 241 [TAN] |
| *Jumellea brevifolia* H.Perrier | JQ905464 | Rakotoarivelo & al. 300 [TAN] | — | — | JQ905285 | Rakotoarivelo & al. 300 [TAN] |
| *Jumellea comorensis* (Rchb.f.) Schltr. | JQ905465 | Rakotoarivelo & al. 040 [REU] | — | — | JQ905286 | Rakotoarivelo & al. 040 [REU] |

| | | | | | | |
|---|---|---|---|---|---|---|
| *Jumellea confusa* (Schltr.) Schltr. | JQ905501 | Rakotoarivelo & al. 088 [TAN] | — | — | JQ905318 | Rakotoarivelo & al. 088 [TAN] |
| *Jumellea densefoliata* Senghas | JQ905466 | Rakotoarivelo & al. 109 [TAN] | — | — | JQ905287 | Rakotoarivelo & al. 109 [TAN] |
| *Jumellea divaricata* (Frapp. ex Cordem.) Schltr. | JQ905467 | Pailler 267 [REU] | — | — | JQ905288 | Pailler 267 [REU] |
| *Jumellea exilis* (Cordem.) Schltr. | JQ905468 | Pailler 216 [REU] | — | — | JQ905289 | Pailler 216 [REU] |
| *Jumellea fragrans* (Thouars) Schltr. | KF558091 | Micheneau & Pailler 10 [REU] | — | — | — | — |
| *Jumellea francoisii* Schltr. | JQ905472 | Rakotoarivelo & al. 230 [TAN] | — | — | JQ905293 | Rakotoarivelo & al. 230 [TAN] |
| *Jumellea hyalina* H.Perrier | JQ905476 | Rakotoarivelo & al. 006 [TAN] | — | — | JQ905297 | Rakotoarivelo & al. 006 [TAN] |
| *Jumellea ibityana* Schltr. | JQ905477 | Rakotoarivelo & al. 031 [TAN] | — | — | JQ905298 | Rakotoarivelo & al. 031 [TAN] |
| *Jumellea imerinensis* Schltr. | — | — | JQ905530 | Rakotoarivelo *et al.* 317 [TAN] | JQ905290 | Rakotoarivelo *et al.* 317 [TAN] |
| *Jumellea jumelleana* (Schltr.) Summerh. | JQ905478 | Rakotoarivelo & al. 099 [TAN] | — | — | JQ905299 | Rakotoarivelo & al. 099 [TAN] |
| *Jumellea lignosa* (Schltr.) Schltr. | JQ905479 | Rakotoarivelo & al. 036 [TAN] | — | — | JQ905300 | Rakotoarivelo & al. 036 [TAN] |
| *Jumellea linearipetala* H.Perrier | Farminhão *et al.* (2020) | No voucher (CM6m) | KF558160 | No voucher (CM6m) | Farminhão *et al.* (2020) | No voucher (CM6m) |
| *Jumellea longivaginans* H.Perrier | — | — | JQ905536 | Rakotoarivelo *et al.* 150 [TAN] | JQ905296 | Rakotoarivelo *et al.* 150 [TAN] |
| *Jumellea majalis* (Schltr.) Schltr. | JQ905488 | Rakotoarivelo & al. 307 [TAN] | — | — | JQ905307 | Rakotoarivelo & al. 307 [TAN] |
| *Jumellea major* Schltr. | — | — | JQ905546 | Rakotoarivelo *et al.* 322 [TAN] | JQ905304 | Rakotoarivelo *et al.* 322 [TAN] |
| *Jumellea maxillarioides* (Ridl.) Schltr. | Farminhão *et al.* (2020) | 1T53 [TAN] | Farminhão *et al.* (2020) | 1T53 [TAN] | Farminhão *et al.* (2020) | 1T53 [TAN] |
| *Jumellea pachyceras* Schltr. | — | — | JQ905550 | Rakotoarivelo *et al.* 311 [TAN] | JQ905308 | Rakotoarivelo *et al.* 311 [TAN] |
| *Jumellea pailleri* F.Rakotoar. | JQ905490 | Rakotoarivelo & al. 060 [REU] | — | — | JQ905309 | Rakotoarivelo & al. 060 |

| | | | | | | |
|---|---|---|---|---|---|---|
| | | | | | | [REU] |
| *Jumellea papangensis* H.Perrier | Farminhão *et al.* (2020) | Chase 17913 [K] | KF558251 | Chase 17913 [K] | Farminhão *et al.* (2020) | Chase 17913 [K] |
| *Jumellea peyrotii* Bosser | — | — | JQ905552 | Rakotoarivelo *et al.* 144 [TAN] | JQ905310 | Rakotoarivelo *et al.* 144 [TAN] |
| *Jumellea punctata* H.Perrier | XX | AMB4870 [] | — | — | XX | AMB4870 [] |
| *Jumellea recta* (Thouars) Schltr. | Farminhão *et al.* (2020) | Fournel 114 [REU] | KF558246 | Fournel 114 [REU] | Farminhão *et al.* (2020) | Fournel 114 [REU] |
| *Jumellea recurva* (Thouars) Schltr. | JQ905494 | Pailler 204 [REU] | — | — | JQ905313 | Pailler 204 [REU] |
| *Jumellea rigida* Schltr. | JQ905498 | Rakotoarivelo & al. 220 [TAN] | — | — | JQ905315 | Rakotoarivelo & al. 220 [TAN] |
| *Jumellea rossii* Senghas | JQ905499 | Pailler 293 [REU] | — | — | JQ905316 | Pailler 293 [REU] |
| *Jumellea similis* Schltr. | JQ905503 | Rakotoarivelo & al. 100 [TAN] | — | — | JQ905320 | Rakotoarivelo & al. 100 [TAN] |
| *Jumellea spathulata* (Ridl.) Schltr. | — | — | JQ905563 | Rakotoarivelo *et al.* 209 [TAN] | JQ905319 | Rakotoarivelo *et al.* 209 [TAN] |
| *Jumellea stenoglossa* H.Perrier | — | — | Farminhão *et al.* (2020) | 87T124 [TAN] | JQ905323 | Pailler 239 [TAN] |
| *Jumellea stenophylla* (Frapp. ex Cordem.) Schltr. | Farminhão *et al.* (2020) | Micheneau & Pailler 92 [REU] | KF558165 | Micheneau & Pailler 92 [REU] | Farminhão *et al.* (2020) | Micheneau & Pailler 92 [REU] |
| *Jumellea tenuibracteata* (H.Perrier ex Hermans) F.P.Rakotoar. & Pailler | JQ905482 | Rakotoarivelo & al. 321 [TAN] | — | — | — | — |
| *Jumellea teretifolia* Schltr. | JQ905508 | Rakotoarivelo & al. 160 [TAN] | — | — | JQ905325 | Rakotoarivelo & al. 160 [TAN] |
| *Jumellea triquetra* (Thouars) Schltr. | KF558112 | Micheneau & Pailler 11 [REU] | — | — | XX | Micheneau & Pailler 11 [REU] |
| *Jumellea unguicularis* Schltr. | JQ905511 | Rakotoarivelo & al. 216 [TAN] | — | — | JQ905328 | Rakotoarivelo & al. 216 [TAN] |
| *Jumellea walleri* (Rolfe) la Croix | Farminhão *et al.* (2020) | No voucher (CM6) | KF558257 | No voucher (CM6) | Farminhão *et al.* (2020) | No voucher (CM6) |
| *Jumellea zaratananae* Schltr. | JQ905510 | Rakotoarivelo & al. 326 [TAN] | — | — | JQ905327 | Rakotoarivelo & al. 326 |

| | | | | | | |
|---|---|---|---|---|---|---|
| | | | | | | [TAN] |
| *Kylicanthe cornuata* Descourv., Stévart & Droissart | Farminhão *et al.* (2020) | YAS 2498 [BRLU] | MH23746 1 | YAS 2498 [BRLU] | Farminhão *et al.* (2020) | YAS 2498 [BRLU] |
| *Kylicanthe quintasii* (Rolfe) Farminhão, Stévart & Droissart | Farminhão *et al.* (2020) | Bom Sucesso 1981.1 [BRLU] | MH23759 4 | Stévart 4708 [BRLU] | Farminhão *et al.* (2020) | Stévart 4708 [BRLU] |
| *Lemurella pallidiflora* Bosser | — | — | DQ09145 4 | Kew accession 4958 [K] | — | — |
| *Listrostachys pertusa* (Lindl.) Rchb.f. | MK697536 | YAS 2522 [BRLU] | MH23746 3 | YAS 2522 [BRLU] | MK722033 | YAS 2522 [BRLU] |
| *Microcoelia aphylla* (Thouars) Summerh. | Farminhão *et al.* (2020) | Carlsward 341 (BRLU] | DQ09152 5 | Carlsward 341 (BRLU] | EU490751 | Carlsward 341 (BRLU] |
| *Microcoelia bulbocalcarata* L.Jonss. | Farminhão *et al.* (2020) | Bom Sucesso 917 [BRLU] | DQ09152 1 | No voucher (Carlsward *et al.* 2006) | Farminhão *et al.* (2020) | Bom Sucesso 917 [BRLU] |
| *Microcoelia caespitosa* (Rolfe) Summerh. | — | — | — | — | — | — |
| *Microcoelia koehleri* (Schltr.) Summerh. | — | — | — | — | Farminhão *et al.* (2020) | BR 19910194-71 |
| *Mystacidium aliceae* Bolus | — | — | — | — | MG250275 | Martos 775 [NU] |
| *Mystacidium braybontiae* Summerh. | XX | Carlsward 179 (FLAS) | XX | Carlsward 179 (FLAS) | XX | Carlsward 179 (FLAS) |
| *Mystacidium capense* (L.f.) Schltr. | Farminhão *et al.* (2020) | Whitten 1781 [FLAS] | DQ09148 7 | Whitten 1781 [FLAS] | Farminhão *et al.* (2020) | Whitten 1781 [FLAS] |
| *Mystacidium flanaganii* (Bolus) Bolus | — | — | — | — | MG250276 | Young 1326 [NU] |
| *Mystacidium gracile* Harv. | — | — | — | — | MG250260 | Bytebier 2227 [NU] |
| *Mystacidium pusillum* Harv. | — | — | MG25024 4 | Martos 740 [NU] | MG250273 | Martos 740 [NU] |
| *Mystacidium venosum* Harv. ex Rolfe | Farminhão *et al.* (2020) | Hermans 5084 (K] | MG25024 3 | Martos 733 [NU] | MG250270 | Hermans 5084 (K] |
| *Mystacidium tanganyikense* Summerh. | — | — | — | — | — | — |
| *Neobathiea grandidieriana* (Rchb.f.) Garay | Farminhão *et al.* (2020) | Carlsward 395 [FLAS] | DQ09145 3 | Carlsward 395 [FLAS] | Farminhão *et al.* (2020) | Carlsward 395 [FLAS] |
| *Nephrangis* | Farminhão | YAS 2916 | MH23755 | SIB 1690 | Farminhão *et* | SIB 1690 |

| | | | | | | |
|---|---|---|---|---|---|---|
| *filiformis* (Kraenzl.) Summerh. | *et al.* (2020) | [BRLU] | 4 | [BRLU] | *al.* (2020) | [BRLU] |
| *Oeonia rosea* Ridl. | Farminhão *et al.* (2020) | Whitten 1813 [FLAS] | DQ091452 | Whitten 1813 [FLAS] | Farminhão *et al.* (2020) | Whitten 1813 [FLAS] |
| *Oeoniella polystachys* (Thouars) Schltr. | Farminhão *et al.* (2020) | Carlsward 221 [FLAS] | DQ091557 | Carlsward 221 [FLAS] | EU490756 | Carlsward 221 [FLAS] |
| *Phalaenopsis wilsonii* Rolfe | — | — | AF533475 | C.Tsai 1645 [TNM] | EU490763 | Carlsward 331 [FLAS] |
| *Planetangis longicaudata* (Rolfe) Stévart & Farminhão | Farminhão *et al.* (2020b) | BTO181 [BRLU] | — | — | Farminhão *et al.* (2020b) | BTO181 [BRLU] |
| *Plectrelminthus caudatus* (Lindl.) Summerh. | — | — | MH237470 | YAS 2803 [BRLU] | Farminhão *et al.* (2020) | YAS 2803 [BRLU] |
| *Podangis dactyloceras* (Rchb.f.) Schltr. | Farminhão *et al.* (2020) | YAS 2652 [BRLU] | MH237469 | YAS 2652 [BRLU] | Farminhão *et al.* (2020) | YAS 2652 [BRLU] |
| *Podangis muscicola* (Rchb.f.) Farminhão & D'haijère | Farminhão *et al.* (2020) | Carlsward 169 [SEL] | DQ091513 | Carlsward 169 [SEL] | EU490774 | Carlsward 169 [SEL] |
| *Podangis rhipsalisocia* (Rchb.f.) P.J.Cribb & Carlsward | Farminhão *et al.* (2020) | YAS 2011 [BRLU] | MH237473 | YAS 2011 [BRLU] | Farminhão *et al.* (2020) | YAS 2011 [BRLU] |
| *Rhipidoglossum arbonnieri* (Geerinck) Eb.Fisch., Killmann, J.-P.Lebel & Delep. | — | — | — | — | XX | See p. 374 in The Orchids of Rwanda [BRLU] |
| *Rhipidoglossum brachyceras* (Summerh.) Farminhão & Stévart | Farminhão *et al.* (2020) | Bytebier 361 [EA] | DQ091490 | Bytebier 361 [EA] | MG250257 | Bytebier 361 [EA] |
| *Rhipidoglossum brevifolium* Summerh. | Farminhão *et al.* (2020) | Farminhão *et al.* 95 [BRLU] | Farminhão *et al.* (2020) | Farminhão *et al.* 95 [BRLU] | Farminhão *et al.* (2020) | Farminhão *et al.* 95 [BRLU] |
| *Rhipidoglossum burttii* (Summerh.) Summerh. | Farminhão *et al.* (2020) | Photo voucher (see p. 279 in The Orchids of Rwanda) | Farminhão *et al.* (2020) | Farminhão & Dumbo 228 [BRLU] | Farminhão *et al.* (2020) | Farminhão & Dumbo 228 [BRLU] |
| *Rhipidoglossum caffrum* (Bolus) Farminhão & Stévart | — | — | — | — | MG250274 | Martos 752 [NU] |
| *Rhipidoglossum confusum* | Farminhão *et al.* | Kew accession 3936 [K] | Farminhão *et al.* | Farminhão *et al.* 96 [BRLU] | MG250269 | Kew accession 3936 [K] |

| | | | | | | |
|---|---|---|---|---|---|---|
| (P.J.Cribb) Farminhão & Stévart | (2020) | | (2020) | | | |
| *Rhipidoglossum curvatum* (Rolfe) Garay | XX | YAS 1277 [BRLU] | — | — | XX | YAS 1277 [BRLU] |
| *Rhipidoglossum delepierreanum* (J.-P.Lebel & Geerinck) Eb.Fisch., Killmann, J.-P.Lebel & Delep. | Farminhão *et al.* (2020) | Photo voucher (see p. 377 in The Orchids of Rwanda) | MH237376 | Photo voucher (see p. 377 in The Orchids of Rwanda) | Farminhão *et al.* (2020) | Photo voucher (see p. 377 in The Orchids of Rwanda) |
| *Rhipidoglossum densiflorum* Summerh. | Farminhão *et al.* (2020) | YAS 2154 [BRLU] | MH237480 | YAS 2154 [BRLU] | Farminhão *et al.* (2020) | YAS 2154 [BRLU] |
| *Rhipidoglossum eggelingii* (Summerh.) Farminhão & Stévart | Farminhão *et al.* (2020) | Farminhão & Dumbo 253 [BRLU] | Farminhão *et al.* (2020) | Farminhão & Dumbo 232 [BRLU] | Farminhão *et al.* (2020) | Farminhão & Dumbo 253 [BRLU] |
| *Rhipidoglossum globulosocalcaratum* (De Wild.) Summerh. | Farminhão *et al.* (2020) | YAS 2837 [BRLU] | MH237474 | YAS 2837 [BRLU] | Farminhão *et al.* (2020) | YAS 2837 [BRLU] |
| *Rhipidoglossum kamerunense* (Schltr.) Garay | Farminhão *et al.* (2020) | Nkonmeneck 300 [SEL] | MH237181 | Nkonmeneck 300 [SEL] | MG250264 | Nkonmeneck 300 [SEL] |
| *Rhipidoglossum millarii* (Bolus) Farminhão & Stévart | Farminhão *et al.* (2020) | Carlsward 346 [FLAS] | MH237581 | Carlsward 346 [FLAS] | MG250267 | Carlsward 346 [FLAS] |
| Rhipidoglossum pendulum (la Croix & P.J.Cribb) Farminhão & Stévart | Farminhão *et al.* (2020) | Bom Sucesso 2155.1 [BRLU] | MH237381 | Farminhão & Stévart 2 [BRLU] | Farminhão *et al.* (2020) | Bom Sucesso 2155.1 [BRLU] |
| *Rhipidoglossum polydactylum* (Kraenzl.) Garay | Farminhão *et al.* (2020) | Droissart & Stévart 699 [BRLU] | Farminhão *et al.* (2020) | Droissart & Stévart 699 [BRLU] | Farminhão *et al.* (2020) | Droissart & Stévart 699 [BRLU] |
| *Rhipidoglossum pulchellum* var. *geniculatum* (Summerh.) Garay | Farminhão *et al.* (2020) | YAS 3043 [BRLU] | MH237477 | YAS 3043 [BRLU] | Farminhão *et al.* (2020) | YAS 3043 [BRLU] |
| *Rhipidoglossum pusillum* (Summerh.) Farminhão & Stévart | Farminhão *et al.* (2020) | Photo voucher (see p. 97 in The Orchids of Rwanda) | MH237525 | Photo voucher (see p. 97 in The Orchids of Rwanda) | Farminhão *et al.* (2020) | Photo voucher (see p. 97 in The Orchids of Rwanda) |
| *Rhipidoglossum rutilum* (Rchb.f.) Schltr. | Farminhão *et al.* (2020) | YAS 2399 [BRLU] | Farminhão *et al.* (2020) | YAS 2399 [BRLU] | Farminhão *et al.* (2020) | YAS 2399 [BRLU] |
| *Rhipidoglossum* | XX | BR5629 [BR] | XX | BR5629 [BR] | XX | BR5629 [BR] |

| | | | | | | |
|---|---|---|---|---|---|---|
| *stolzii* (Schltr.) Garay | | | | | | |
| *Rhipidoglossum subsimplex* (Summerh.) Garay | Farminhão *et al.* (2020) | Bytebier 546 [EA] | MH23758 3 | Bytebier 546 [EA] | MG250259 | Bytebier 546 [EA] |
| *Rhipidoglossum thomense* (la Croix & P.J.Cribb) Farminhão & Stévart | Farminhão *et al.* (2020) | Stévart 246 [BRLU] | MH23738 0 | Stévart 246 [BRLU] | Farminhão *et al.* (2020) | Stévart 246 [BRLU] |
| *Rhipidoglossum xanthopollinium* (Rchb.f.) Schltr. | Farminhão *et al.* (2020) | Carlsward 384 [FLAS] | MH23757 9 | Carlsward 384 [FLAS] | EU490775 | Carlsward 384 [FLAS] |
| *Solenangis clavata* (Rolfe) Schltr. | XX | Texier 1972 [BRLU] | DQ09153 4 | Carlsward 397 [FLAS] | Farminhão *et al.* (2020) | Texier *et al.* 1972 [BRLU] |
| *Solenangis liberica* (Mansf.) R.Rice | — | — | — | — | XX | D'haijère 7 [BRLU] |
| *Solenangis saotomensis* R.Rice | XX | Stévart 274 [BRLU] | — | — | XX | Stévart 274 [BRLU] |
| *Solenangis scandens* (Schltr.) Schltr. | Farminhão *et al.* (2020) | Yaoundé 3308 | MK72204 4 | Yaoundé 3308 | MK721987 | Yaoundé 3308 |
| *Solenangis impraedicta* sp. nov. | XX | RBR418 [BRLU] | XX | RBR418 [BRLU] | XX | RBR418 [BRLU] |
| *Solenangis wakefieldii* (Rolfe) P.J.Cribb & J.Stewart | — | — | — | — | — | — |
| *Sphyrarhynchus amaniensis* (Summerh.) Bytebier | Farminhão *et al.* (2020) | No voucher (B252) | DQ09148 2 | No voucher (B252) | Farminhão *et al.* (2020) | Farminhão *et al.* 160 [BRLU] |
| *Sphyrarhynchus brevilobus* (Summerh.) Bytebier | Farminhão *et al.* (2020) | Bytebier 307 [EA] | DQ09148 3 | Bytebier 307 [EA] | MG250256 | Bytebier 307 [EA] |
| *Sphyrarhynchus schliebenii* Mansf. | — | — | DQ09148 4 | Bytebier 393 [EA] | MG250258 | Bytebier 393 [EA] |
| *Summerhayesia laurentii* (De Wild.) P.J.Cribb | — | — | Farminhão *et al.* (2020) | BTO 220 | Farminhão *et al.* (2020) | Nimba 965 [BRLU] |
| *Tridactyle anthomaniaca* (Rchb.f.) Summerh. | MK697519 | Y3679 RH (cultivated at YAS) | MH23735 9 | Y3679 RH (cultivated at YAS) | MK722009 | YAS 2462 [BRLU] |
| *Tridactyle aurantiopunctata* P.J.Cribb & Stévart | KF672290 | Stévart 656 [BRLU] | KF662356 | Stévart 656 [BRLU] | Farminhão *et al.* (2020) | Stévart 290 [BRLU] |
| *Tridactyle bicaudata* (Lindl.) Schltr. | KF672305 | KIS 135 (cultivated at Kisantu shade | MH23758 6 | Bytebier 348 [EA] | Farminhão *et al.* (2020) | Bytebier 348 [EA] |

| | | | | | | |
|---|---|---|---|---|---|---|
| | | house) | | | | |
| *Tridactyle brevicalcarata* Summerh. | MK697526 | BR 20090424-75 | MH23755 5 | BTO 168 [BRLU] | MK722037 | BR 20090424-75 |
| *Tridactyle clavata* (Summerh.) R.Rice | Farminhão *et al.* (2020) | Photo voucher (see p. 201 in The Orchids of Rwanda) | MH23753 2 | Photo voucher (see p. 201 in The Orchids of Rwanda) | Farminhão *et al.* (2020) | Photo voucher (see p. 201 in The Orchids of Rwanda) |
| *Tridactyle crassifolia* Summerh. | MK697517 | Nkongmeneck 2076 [SEL] | MH23756 4 | Nkongmeneck 2076 [SEL] | MK722008 | Nkongmeneck 2076 [SEL] |
| *Tridactyle exellii* P.J.Cribb & Stévart | MK697533 | Stévart 297 [BRLU] | MH23740 0 | Stévart 297 [BRLU] | MK722017 | Stévart 297 [BRLU] |
| *Tridactyle filifolia* (Schltr.) Schltr. | MK697513 | Bytebier 707 [EA] | MH23758 4 | Bytebier 707 [EA] | MK722021 | Bytebier 707 [EA] |
| *Tridactyle gentilii* (De Wild.) Schltr. | Farminhão *et al.* (2020) | Stévart & Pial 280 [BRLU] | MH23739 6 | Stévart & Pial 280 [BRLU] | Farminhão *et al.* (2020) | Stévart & Pial 280 [BRLU] |
| *Tridactyle latifolia* Summerh. | MK721988 | Primo and Stevart 94 [BRLU] | MK69750 9 | Primo and Stevart 94 [BRLU] | MK722020 | Primo and Stevart 94 [BRLU] |
| *Tridactyle laurentii* (De Wild.) Schltr. | MK697523 | YAS 2489 [BRLU] | MH23736 5 | YAS 2489 [BRLU] | MK722027 | No voucher |
| *Tridactyle ligulifolia* (Summerh.) R.Rice | Farminhão *et al.* (2020) | Photo voucher (see p. 202 in The Orchids of Rwanda) | MH23751 4 | Photo voucher (see p. 202 in The Orchids of Rwanda) | Farminhão *et al.* (2020) | Photo voucher (see p. 202 in The Orchids of Rwanda) |
| *Tridactyle minutifolia* Stévart & D'haijère | Farminhão *et al.* (2020) | Stévart 3609 [BRLU] | MH23738 4 | Stévart 3609 [BRLU] | Farminhão *et al.* (2020) | G.Brice 252 [BRLU] |
| *Tridactyle muriculata* (Rendle) Schltr. | MK697512 | YAS 2189 [BRLU] | MH23736 6 | YAS 2189 [BRLU] | MK22006 | YAS 2189 [BRLU] |
| *Tridactyle nalaensis* (De Wild.) Schltr. | MK697534 | BT091 [BRLU] | MH23755 3 | BT091 [BRLU] | — | — |
| *Tridactyle scottellii* (Rendle) Schltr. | — | — | Farminhão *et al.* (2020) | Photo voucher (BTO 231) | Farminhão *et al.* (2020) | Ndong Bokung *et al.* 337 [BRLU] |
| *Tridactyle thomensis* P.J.Cribb & Stévart | Farminhão *et al.* (2020) | Stévart 1217 [BRLU] | MH23740 4 | Stévart 296 [BRLU] | Farminhão *et al.* (2020) | Stévart 1217 [BRLU] |
| *Tridactyle tridactylites* (Rolfe) Schltr. | Farminhão *et al.* (2020) | Stévart 1229 [BRLU] | MH23740 7 | Stévart 1229 [BRLU] | Farminhão *et al.* (2020) | Stévart 1229 [BRLU] |
| *Tridactyle truncatiloba* Summerh. | Farminhão *et al.* (2020) | Stévart *et al.* 1624 [BRLU] | MH23741 0 | Stévart *et al.* 1624 [BRLU] | Farminhão *et al.* (2020) | Stévart *et al.* 1679 [BRLU] |
| *Vanda falcata* (Thunb.) Beer | — | — | DQ09144 2 | Carlsward 163 [SEL] | — | — |
| *Ypsilopus* | MK697524 | Bytebier & Kirika | DQ09138 | Bytebier & | MK722013 | Bytebier & |

| | | | | | | |
|---|---|---|---|---|---|---|
| *amaniensis* (Kraenzl.) D'haijère & Stévart | | 26 [EA] | 6 | Kirika 26 [EA] | | Kirika 26 [EA] |
| *Ypsilopus erectus* (P.J.Cribb) P.J.Cribb & J.Stewart | MK697515 | Grieve 1244 [EA] | MK72204 2 | Grieve 1244 [EA] | MK722022 | Grieve 1244 [EA] |
| *Ypsilopus furcistipes* (Summerh.) D'haijère & Stévart | MK697520 | Bytebier 1731 [EA] | DQ09151 8 | Bytebier 1731 [EA] | MK722025 | Bytebier 1731 [EA] |
| *Ypsilopus longifolius* (Kraenzl.) Summerh. | Farminhão *et al.* (2020) | Bytebier 609 [EA] | DQ09151 9 | Bytebier 609 [EA] | Farminhão *et al.* (2020) | Bytebier 609 [EA] |
| *Ypsilopus schliebenii* (Mansf.) D'haijère & Stévart | — | — | MH23733 4 | BR 20090389-40 | — | — |
| *Ypsilopus tanneri* (P.J.Cribb) D'haijère & Stévart | MK697518 | PCP 198 [EA] | DQ09152 0 | PCP 198 [EA] | MK722024 | PCP 198 [EA] |
| *Ypsilopus tricuspis* (Bolus) D'haijère & Stévart | XX | Farminhão 306 [BRLU] | — | — | XX | Farminhão 306 [BRLU] |
| *Ypsilopus viridiflorus* P.J.Cribb & J.Stewart | MK721971 | Bytebier 402 [EA] | — | — | MK722003 | Bytebier 402 [EA] |

Table S2. AICc comparison of BioGeoBEARS models

d, e, j: values for dispersal, extinction and founder-event speciation parameters, respectively. δAICc: difference of AICc between the model and the model with minimal AICc. AICc weights: proportion of the total predictive power of all models explained by the model.

| Model | Number of parameters | d | e | j | Log-likelihood | AIC | AICc | δAICc | AICc weigths |
|---|---|---|---|---|---|---|---|---|---|
| DEC*+J | 3 | 0 | 0.14 | 0 | -70.09 | 146.18 | 146.25 | 0 | 0.67 |
| DEC* | 2 | 0.01 | 0.14 | 0 | -72.84 | 149.68 | 149.71 | 3.46 | 0.12 |
| BAYAREALIKE*+J | 3 | 0 | 0.27 | 0 | -72.04 | 150.08 | 150.15 | 3.9 | 0.09 |
| DIVALIKE* | 2 | 0.01 | 0.13 | 0 | -73.11 | 150.23 | 150.26 | 4.01 | 0.09 |
| DIVALIKE*+J | 3 | 0.01 | 0.53 | 0 | -73.18 | 152.36 | 152.43 | 6.18 | 0.03 |
| BAYAREALIKE* | 2 | 0.01 | 0.28 | 0 | -78.31 | 160.62 | 160.66 | 14.41 | 0 |

Table S3. AICc comparison of hidden-state SSE models for the flower colour in Angraecinae

CR: Constant-rate model, diversification rates are constant across the phylogeny; ETD: examined trait-dependent model, diversification rates are dependent on the states of the observed character; CID: character-independent model, diversification rates are dependent on a concealed (unobserved) character; ECTD: examined and concealed trait-dependent model, diversification rates are partly dependent on the examined trait states but in conjunction with hidden states. Greek letters τ (tau) and ε (epsilon) indicate the number of parameters for turnover and extinction fraction respectively. δAICc: difference of AICc between the model and the model with minimal AICc. AICc weights: proportion of the total predictive power of all models explained by the model.

| Model | τ | ε | Hidden states | Log-likelihood | AIC | AICc | δAICc | AICc weights |
|---|---|---|---|---|---|---|---|---|
| CID4 | 4 | 1 | 4 | -866.33 | 1748.65 | 1749.11 | 0 | 0.77 |
| ECTD | 4 | 1 | 2 | -865.55 | 1751.09 | 1751.79 | 2.68 | 0.2 |
| CID2 | 2 | 1 | 2 | -871.86 | 1755.73 | 1755.99 | 6.88 | 0.02 |
| ETD | 2 | 1 | 1 | -879.71 | 1769.41 | 1769.6 | 20.5 | 0 |
| CR | 1 | 1 | 1 | -880.87 | 1769.73 | 1769.85 | 20.75 | 0 |

Table S4. AICc comparison of hidden-state SSE models for the resupination in Angraecinae

CR: Constant-rate model, diversification rates are constant across the phylogeny; ETD: examined trait-dependent model, diversification rates are dependent on the states of the observed character; CID: character-independent model, diversification rates are dependent on a concealed (unobserved) character; ECTD: examined and concealed trait-dependent model, diversification rates are partly dependent on the examined trait states but in conjunction with hidden states. Greek letters τ (tau) and ε (epsilon) indicate the number of parameters for turnover and extinction fraction respectively. δAICc: difference of AICc between the model and the model with minimal AICc. AICc weights: proportion of the total predictive power of all models explained by the model.

| Model | τ | ε | Hidden states | Log-likelihood | AIC | AICc | δAICc | AICc weights |
|---|---|---|---|---|---|---|---|---|
| CID4 | 4 | 1 | 4 | -788.57 | 1593.13 | 1593.59 | 0 | 0.97 |
| CID2 | 2 | 1 | 2 | -794.1 | 1600.21 | 1600.47 | 6.88 | 0.03 |
| ECTD | 4 | 1 | 2 | -793.82 | 1607.64 | 1608.33 | 14.75 | 0 |
| CR | 1 | 1 | 1 | -803.11 | 1614.21 | 1614.34 | 20.75 | 0 |
| ETD | 2 | 1 | 1 | -803.03 | 1616.05 | 1616.24 | 22.65 | 0 |

Table S5. AICc comparison of hidden-state SSE models for the floral syndrome in Angraecinae

CR: Constant-rate model, diversification rates are constant across the phylogeny; ETD: examined trait-dependent model, diversification rates are dependent on the states of the observed character; CID: character-independent model, diversification rates are dependent on a concealed (unobserved) character; ECTD: examined and concealed trait-dependent model, diversification rates are partly dependent on the examined trait states but in conjunction with hidden states. Greek letters τ (tau) and ε (epsilon) indicate the number of parameters for turnover and extinction fraction respectively. δAICc: difference of AICc between the model and the model with minimal AICc. AICc weights: proportion of the total predictive power of all models explained by the model.

| Model | τ | ε | Hidden states | log(likelihood) | AIC | AICc | δAICc | AICc weights |
|---|---|---|---|---|---|---|---|---|
| ECTD | 8 | 1 | 2 | -995.83 | 2043.66 | 2048.34 | 0 | 0.99 |
| ETD | 4 | 1 | 1 | -1016 | 2058 | 2059.16 | 10.82 | 0 |
| CID4 | 4 | 1 | 4 | -1016.25 | 2060.5 | 2061.84 | 13.51 | 0 |
| CID8 | 8 | 1 | 8 | -1016.29 | 2068.57 | 2070.79 | 22.45 | 0 |
| CR | 1 | 1 | 1 | -1030.62 | 2081.24 | 2081.93 | 33.59 | 0 |

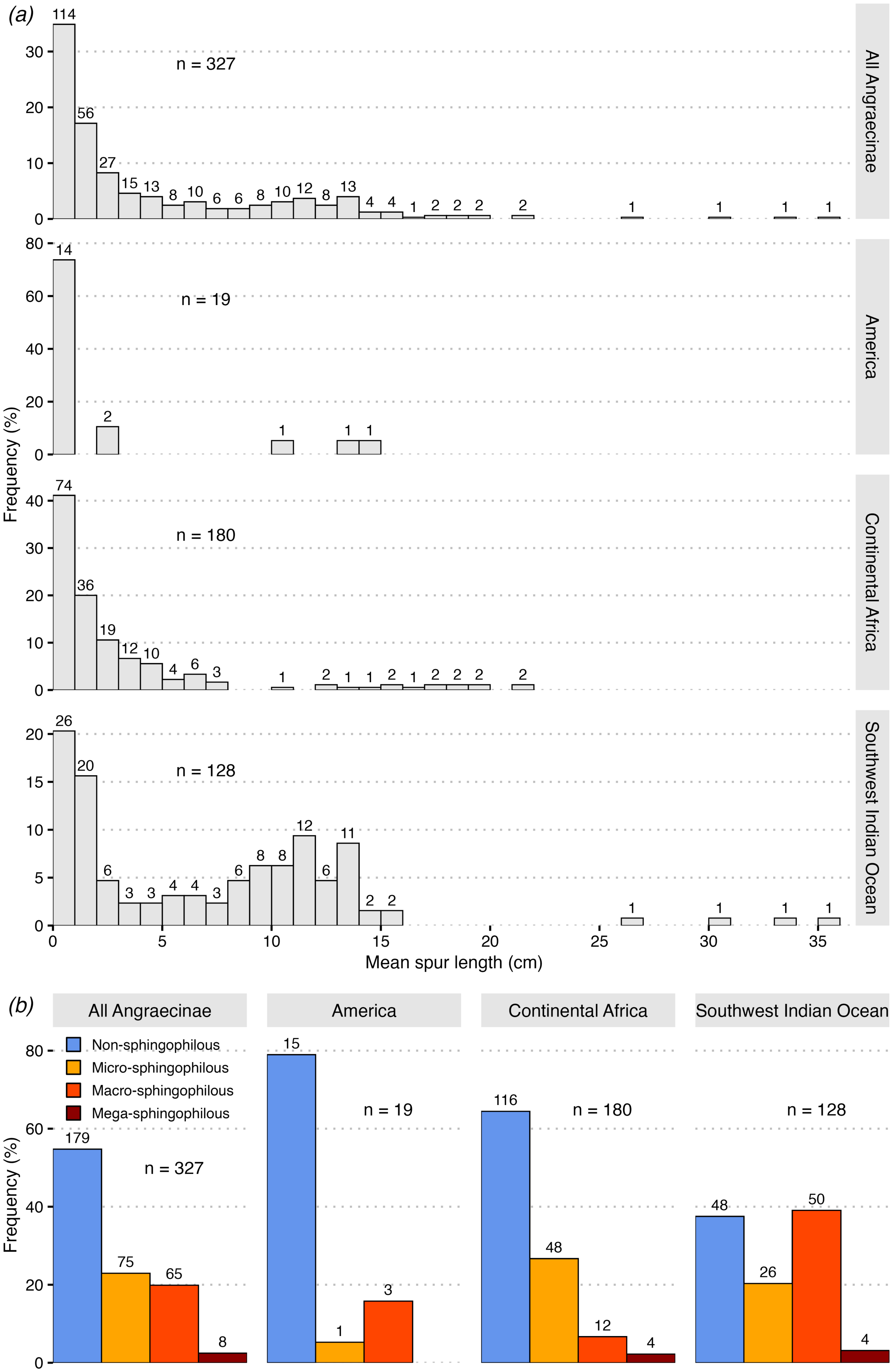

(a)
All Angraecinae
n = 327
America
n = 19
Continental Africa
n = 180
Southwest Indian Ocean
n = 128
Frequency (%)
Mean spur length (cm)
(b)
All Angraecinae
America
Continental Africa
Southwest Indian Ocean
Non-sphingophilous
Micro-sphingophilous
Macro-sphingophilous
Mega-sphingophilous
n = 327
n = 19
n = 180
n = 128
Frequency (%)

Figure S1. Frequency of mean spur lengths and proposed floral syndromes for 327 Angraecinae species and in three geographic areas

(a) Frequency of mean spur lengths, discretised in intervals of one centimeter with values on the left-side included (e.g. values between 0 and 1 include values equal to 0 but not values equal to 1, which are rather included in the interval 1–2). (b) Frequency of floral syndromes. For both (a) and (b), the number of species in each category is indicated above the bars, and n = the total number of species; species ranging across several regions are duplicated.

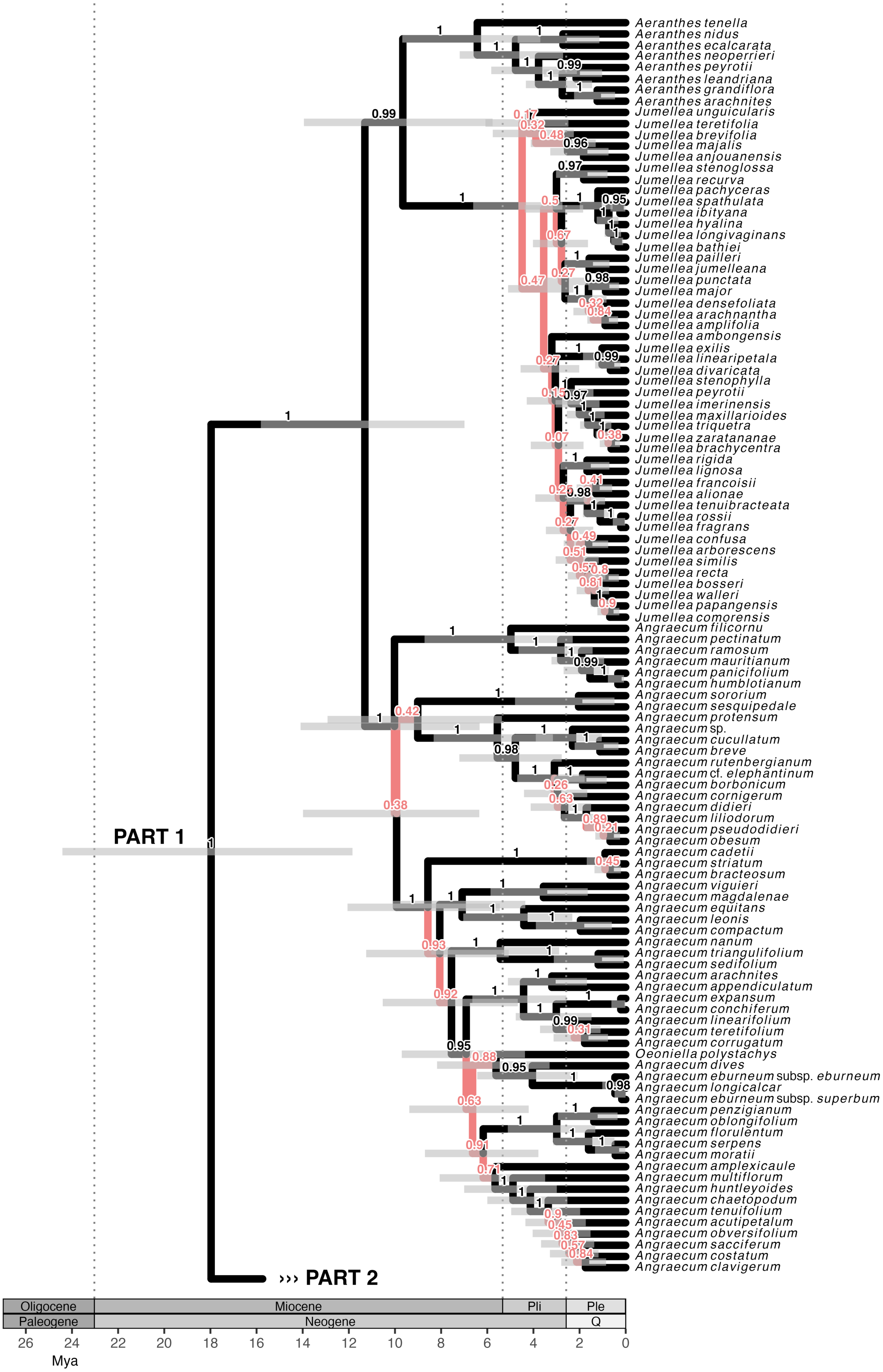
Aeranthes tenella
Aeranthes nidus
Aeranthes ecalcarata
Aeranthes neoperrieri
Aeranthes peyrotii
Aeranthes leandriana
Aeranthes grandiflora
Aeranthes arachnites
Jumellea unguicularis
Jumellea teretifolia
Jumellea brevifolia
Jumellea majalis
Jumellea anjouanensis
Jumellea stenoglossa
Jumellea recurva
Jumellea pachyceras
Jumellea spathulata
Jumellea ibityana
Jumellea hyalina
Jumellea longivaginans
Jumellea bathiei
Jumellea pailleri
Jumellea jumelleana
Jumellea punctata
Jumellea major
Jumellea densefoliata
Jumellea arachnantha
Jumellea amplifolia
Jumellea ambongensis
Jumellea exilis
Jumellea linearipetala
Jumellea divaricata
Jumellea stenophylla
Jumellea peyrotii
Jumellea imerinensis
Jumellea maxillarioides
Jumellea triquetra
Jumellea zaratananae
Jumellea brachycentra
Jumellea rigida
Jumellea lignosa
Jumellea francoisii
Jumellea alionae
Jumellea tenuibracteata
Jumellea rossii
Jumellea fragrans
Jumellea confusa
Jumellea arborescens
Jumellea similis
Jumellea recta
Jumellea bosseri
Jumellea walleri
Jumellea papangensis
Jumellea comorensis
Angraecum filicornu
Angraecum pectinatum
Angraecum ramosum
Angraecum mauritianum
Angraecum panicifolium
Angraecum humblotianum
Angraecum sororium
Angraecum sesquipedale
Angraecum protensum
Angraecum sp.
Angraecum cucullatum
Angraecum breve
Angraecum rutenbergianum
Angraecum cf. elephantinum
Angraecum borbonicum
Angraecum cornigerum
Angraecum didieri
Angraecum liliodorum
Angraecum pseudodidieri
Angraecum obesum
Angraecum cadetii
Angraecum striatum
Angraecum bracteosum
Angraecum viguieri
Angraecum magdalenae
Angraecum equitans
Angraecum leonis
Angraecum compactum
Angraecum nanum
Angraecum triangulifolium
Angraecum sedifolium
Angraecum arachnites
Angraecum appendiculatum
Angraecum expansum
Angraecum conchiferum
Angraecum linearifolium
Angraecum teretifolium
Angraecum corrugatum
Oeoniella polystachys
Angraecum dives
Angraecum eburneum subsp. eburneum
Angraecum longicalcar
Angraecum eburneum subsp. superbum
Angraecum penzigianum
Angraecum oblongifolium
Angraecum florulentum
Angraecum serpens
Angraecum moratii
Angraecum amplexicaule
Angraecum multiflorum
Angraecum huntleyoides
Angraecum chaetopodum
Angraecum tenuifolium
Angraecum acutipetalum
Angraecum obversifolium
Angraecum sacciferum
Angraecum costatum
Angraecum clavigerum
PART 1
››› PART 2
Oligocene
Miocene
Pli
Ple
Paleogene
Neogene
Q
26 24 22 20 18 16 14 12 10 8 6 4 2 0
Mya

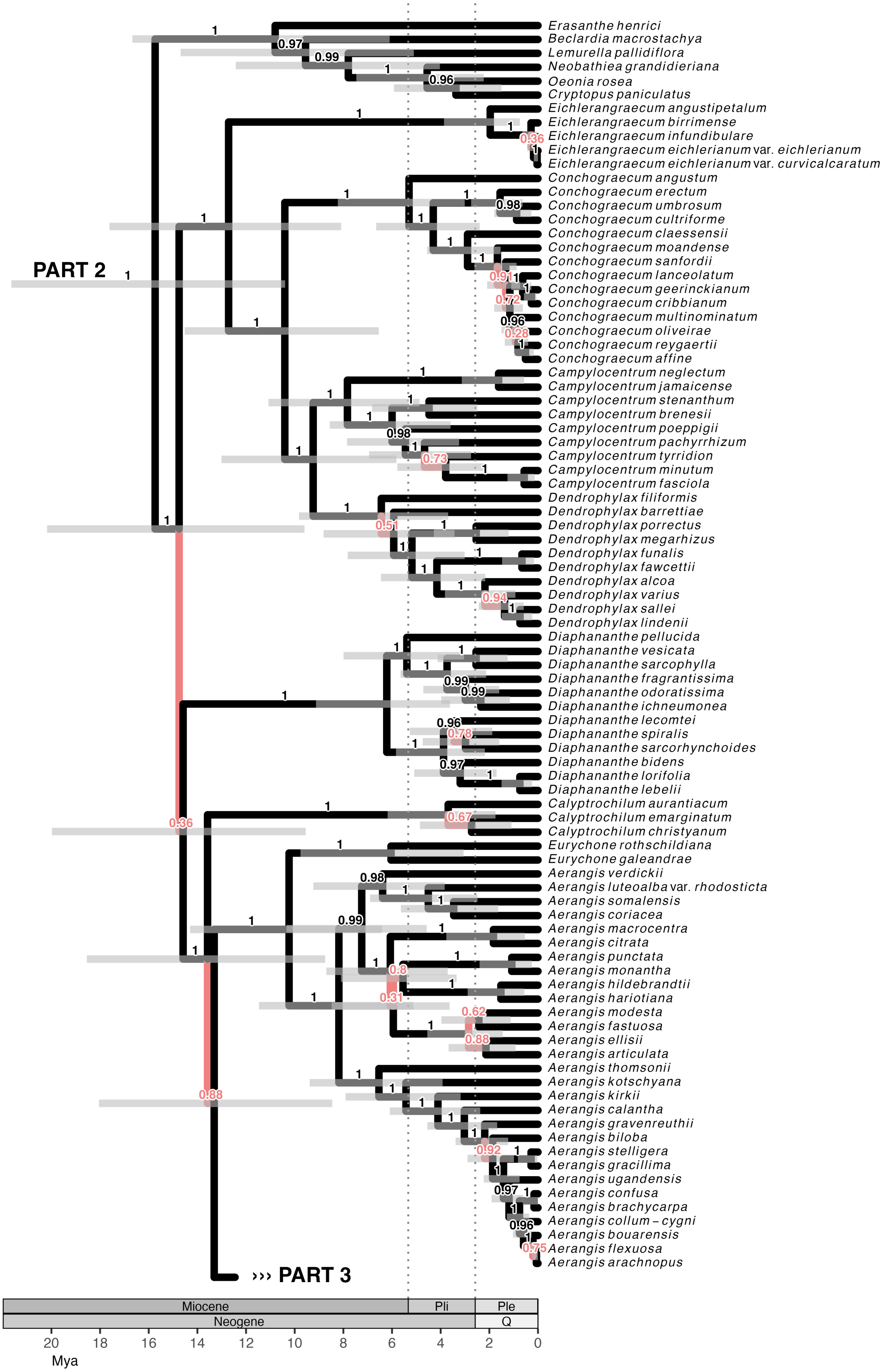
PART 2
››› PART 3
Erasanthe henrici
Beclardia macrostachya
Lemurella pallidiflora
Neobathiea grandidieriana
Oeonia rosea
Cryptopus paniculatus
Eichlerangraecum angustipetalum
Eichlerangraecum birrimense
Eichlerangraecum infundibulare
Eichlerangraecum eichlerianum var. eichlerianum
Eichlerangraecum eichlerianum var. curvicalcaratum
Conchograecum angustum
Conchograecum erectum
Conchograecum umbrosum
Conchograecum cultriforme
Conchograecum claessensii
Conchograecum moandense
Conchograecum sanfordii
Conchograecum lanceolatum
Conchograecum geerinckianum
Conchograecum cribbianum
Conchograecum multinominatum
Conchograecum oliveirae
Conchograecum reygaertii
Conchograecum affine
Campylocentrum neglectum
Campylocentrum jamaicense
Campylocentrum stenanthum
Campylocentrum brenesii
Campylocentrum poeppigii
Campylocentrum pachyrrhizum
Campylocentrum tyrridion
Campylocentrum minutum
Campylocentrum fasciola
Dendrophylax filiformis
Dendrophylax barrettiae
Dendrophylax porrectus
Dendrophylax megarhizus
Dendrophylax funalis
Dendrophylax fawcettii
Dendrophylax alcoa
Dendrophylax varius
Dendrophylax sallei
Dendrophylax lindenii
Diaphananthe pellucida
Diaphananthe vesicata
Diaphananthe sarcophylla
Diaphananthe fragrantissima
Diaphananthe odoratissima
Diaphananthe ichneumonea
Diaphananthe lecomtei
Diaphananthe spiralis
Diaphananthe sarcorhynchoides
Diaphananthe bidens
Diaphananthe lorifolia
Diaphananthe lebelii
Calyptrochilum aurantiacum
Calyptrochilum emarginatum
Calyptrochilum christyanum
Eurychone rothschildiana
Eurychone galeandrae
Aerangis verdickii
Aerangis luteoalba var. rhodosticta
Aerangis somalensis
Aerangis coriacea
Aerangis macrocentra
Aerangis citrata
Aerangis punctata
Aerangis monantha
Aerangis hildebrandtii
Aerangis hariotiana
Aerangis modesta
Aerangis fastuosa
Aerangis ellisii
Aerangis articulata
Aerangis thomsonii
Aerangis kotschyana
Aerangis kirkii
Aerangis calantha
Aerangis gravenreuthii
Aerangis biloba
Aerangis stelligera
Aerangis gracillima
Aerangis ugandensis
Aerangis confusa
Aerangis brachycarpa
Aerangis collum – cygni
Aerangis bouarensis
Aerangis flexuosa
Aerangis arachnopus
Miocene
Pli
Ple
Neogene
Q
20 18 16 14 12 10 8 6 4 2 0
Mya

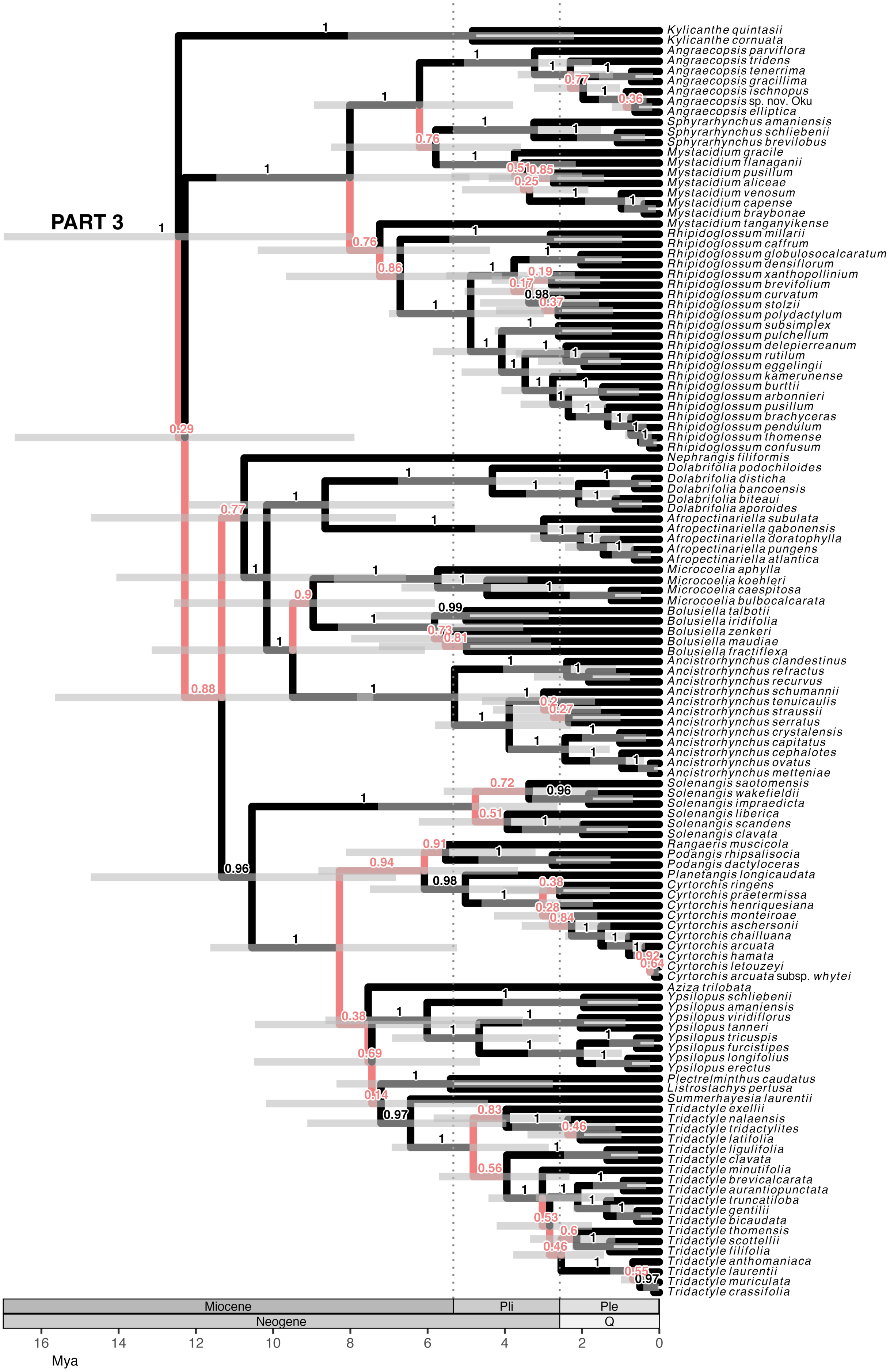
PART 3
Kylicanthe quintasii
Kylicanthe cornuata
Angraecopsis parviflora
Angraecopsis tridens
Angraecopsis tenerrima
Angraecopsis gracillima
Angraecopsis ischnopus
Angraecopsis sp. nov. Oku
Angraecopsis elliptica
Sphyrarhynchus amaniensis
Sphyrarhynchus schliebenii
Sphyrarhynchus brevilobus
Mystacidium gracile
Mystacidium flanaganii
Mystacidium pusillum
Mystacidium aliceae
Mystacidium venosum
Mystacidium capense
Mystacidium braybonae
Mystacidium tanganyikense
Rhipidoglossum millarii
Rhipidoglossum caffrum
Rhipidoglossum globulosocalcaratum
Rhipidoglossum densiflorum
Rhipidoglossum xanthopollinium
Rhipidoglossum brevifolium
Rhipidoglossum curvatum
Rhipidoglossum stolzii
Rhipidoglossum polydactylum
Rhipidoglossum subsimplex
Rhipidoglossum pulchellum
Rhipidoglossum delepierreanum
Rhipidoglossum rutilum
Rhipidoglossum eggelingii
Rhipidoglossum kamerunense
Rhipidoglossum burttii
Rhipidoglossum arbonnieri
Rhipidoglossum pusillum
Rhipidoglossum brachyceras
Rhipidoglossum pendulum
Rhipidoglossum thomense
Rhipidoglossum confusum
Nephrangis filiformis
Dolabrifolia podochiloides
Dolabrifolia disticha
Dolabrifolia bancoensis
Dolabrifolia biteaui
Dolabrifolia aporoides
Afropectinariella subulata
Afropectinariella gabonensis
Afropectinariella doratophylla
Afropectinariella pungens
Afropectinariella atlantica
Microcoelia aphylla
Microcoelia koehleri
Microcoelia caespitosa
Microcoelia bulbocalcarata
Bolusiella talbotii
Bolusiella iridifolia
Bolusiella zenkeri
Bolusiella maudiae
Bolusiella fractiflexa
Ancistrorhynchus clandestinus
Ancistrorhynchus refractus
Ancistrorhynchus recurvus
Ancistrorhynchus schumannii
Ancistrorhynchus tenuicaulis
Ancistrorhynchus straussii
Ancistrorhynchus serratus
Ancistrorhynchus crystalensis
Ancistrorhynchus capitatus
Ancistrorhynchus cephalotes
Ancistrorhynchus ovatus
Ancistrorhynchus metteniae
Solenangis saotomensis
Solenangis wakefieldii
Solenangis impraedicta
Solenangis liberica
Solenangis scandens
Solenangis clavata
Rangaeris muscicola
Podangis rhipsalisocia
Podangis dactyloceras
Planetangis longicaudata
Cyrtorchis ringens
Cyrtorchis praetermissa
Cyrtorchis henriquesiana
Cyrtorchis monteiroae
Cyrtorchis aschersonii
Cyrtorchis chailluana
Cyrtorchis arcuata
Cyrtorchis hamata
Cyrtorchis letouzeyi
Cyrtorchis arcuata subsp. whytei
Aziza trilobata
Ypsilopus schliebenii
Ypsilopus amaniensis
Ypsilopus viridiflorus
Ypsilopus tanneri
Ypsilopus tricuspis
Ypsilopus furcistipes
Ypsilopus longifolius
Ypsilopus erectus
Plectrelminthus caudatus
Listrostachys pertusa
Summerhayesia laurentii
Tridactyle exellii
Tridactyle nalaensis
Tridactyle tridactylites
Tridactyle latifolia
Tridactyle ligulifolia
Tridactyle clavata
Tridactyle minutifolia
Tridactyle brevicalcarata
Tridactyle aurantiopunctata
Tridactyle truncatiloba
Tridactyle gentilii
Tridactyle bicaudata
Tridactyle thomensis
Tridactyle scottellii
Tridactyle filifolia
Tridactyle anthomaniaca
Tridactyle laurentii
Tridactyle muriculata
Tridactyle crassifolia
Miocene
Pli
Ple
Neogene
Q
16
14
12
10
8
6
4
2
0
Mya

Figure S2. Dated maximum-credibility clade phylogenetic tree of 327 Angraecinae species

Posterior probabilities are indicated above branches. Branches with posterior probability <0.95 are coloured coral-red. Pli: Pliocene; Ple: Pleistocene: Q: Quaternary.

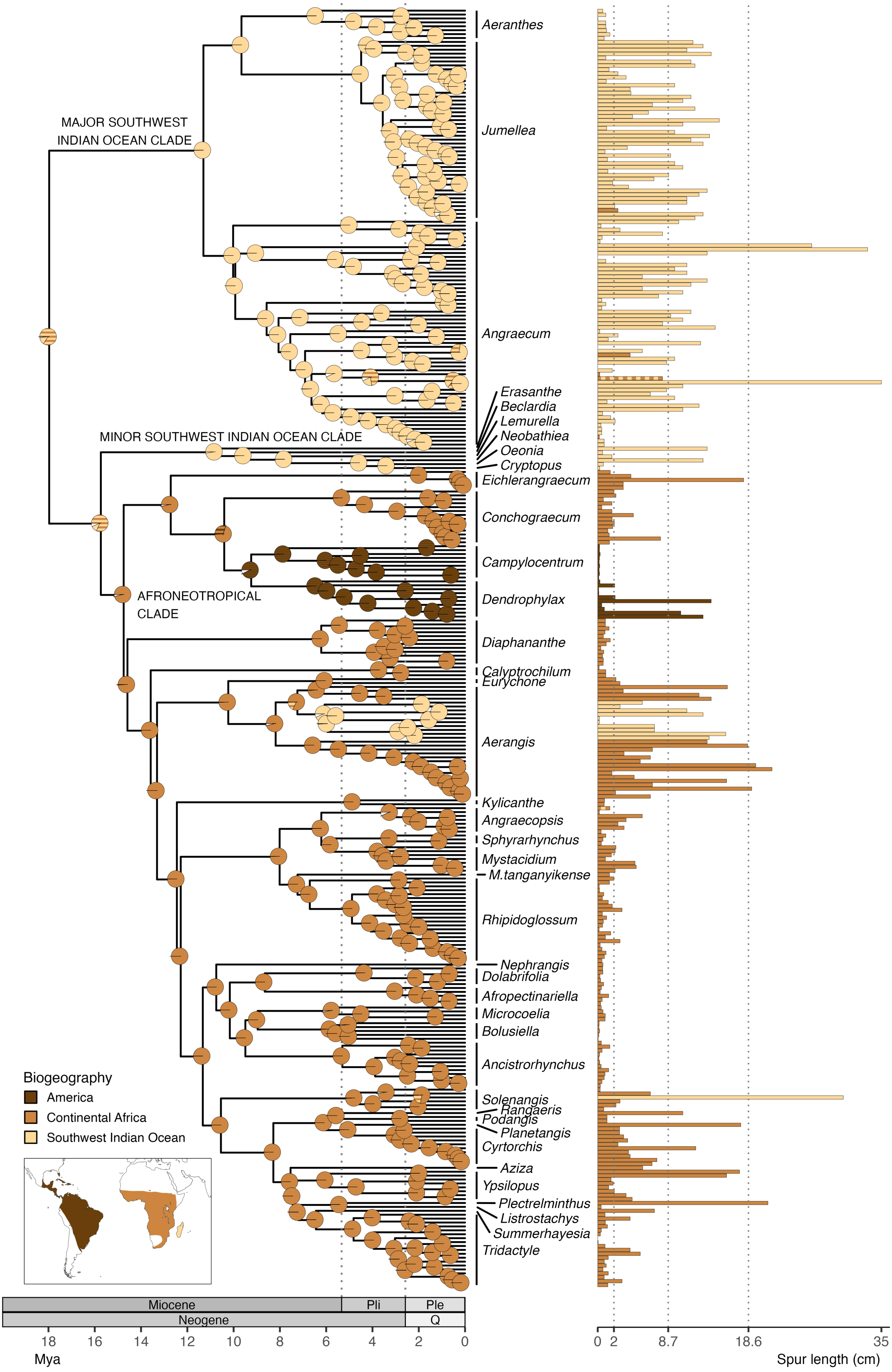
MAJOR SOUTHWEST INDIAN OCEAN CLADE
MINOR SOUTHWEST INDIAN OCEAN CLADE
AFRONEOTROPICAL CLADE
Aeranthes
Jumellea
Angraecum
Erasanthe
Beclardia
Lemurella
Neobathiea
Oeonia
Cryptopus
Eichlerangraecum
Conchograecum
Campylocentrum
Dendrophylax
Diaphananthe
Calyptrochilum
Eurychone
Aerangis
Kylicanthe
Angraecopsis
Sphyrarhynchus
Mystacidium
M.tanganyikense
Rhipidoglossum
Nephrangis
Dolabrifolia
Afropectinariella
Microcoelia
Bolusiella
Ancistrorhynchus
Solenangis
Rangaeris
Podangis
Planetangis
Cyrtorchis
Aziza
Ypsilopus
Plectrelminthus
Listrostachys
Summerhayesia
Tridactyle
Biogeography
America
Continental Africa
Southwest Indian Ocean
Miocene
Pli
Ple
Neogene
Q
18
16
14
12
10
8
6
4
2
0
Mya
0
2
8.7
18.6
35
Spur length (cm)

Figure S3. Estimation of ancestral geographic ranges in the dated phylogenetic tree of 327 angraecoid orchids and mean spur lengths at tips

(a) DEC*+J estimation of ancestral ranges using BioGeoBEARS; the pie charts at nodes represent the ancestral geographic ranges in proportion to their likelihood. These are solid-coloured when the range comprises a single area, or striped when the range comprises several geographical areas. Ranges with a probability <0.1 are merged in white parts of the pies. Map modified from the distribution of Vandeae in Pridgeon AM, Cribb PJ, Chase MW, Rasmussen FN, Pridgeon AM, Cribb PJ, Chase MW, Rasmussen FN, editors. 2014 *Genera Orchidacearum Volume 6: Epidendroideae (Part III)*. Oxford, New York: Oxford University Press. Pli: Pliocene; Ple: Pleistocene; Q: Quaternary. (b) Bar chart showing the mean spur length of species. Bars are filled following actual geographical ranges of species as in (a); vertical dashed lines denote spur lengths thresholds for flower syndromes (see table 1).

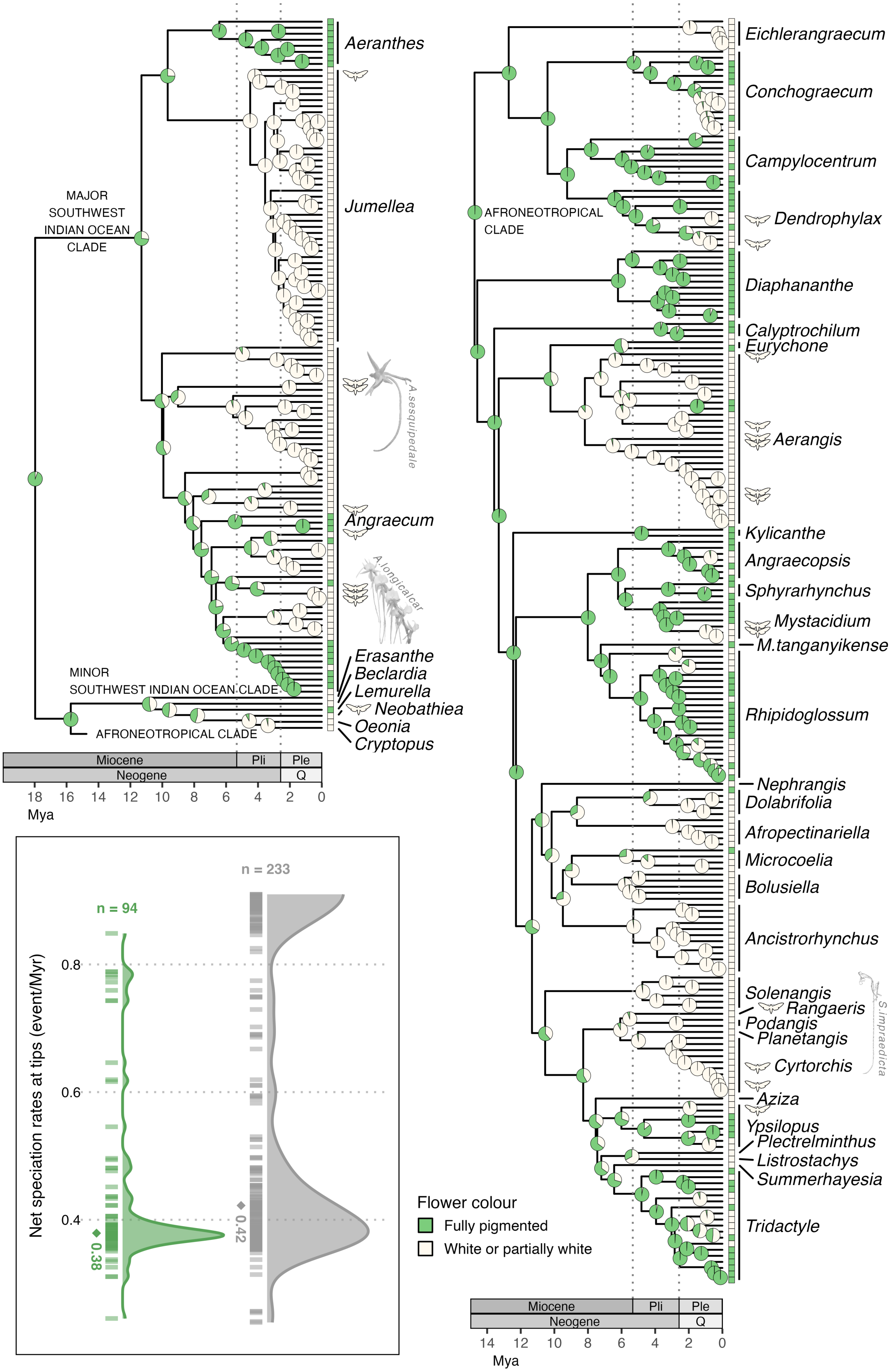
Aeranthes
Jumellea
MAJOR SOUTHWEST INDIAN OCEAN CLADE
A.sesquipedale
Angraecum
A.longicalcar
Erasanthe
Beclardia
Lemurella
Neobathiea
Oeonia
Cryptopus
MINOR SOUTHWEST INDIAN OCEAN CLADE
AFRONEOTROPICAL CLADE
Miocene
Pli
Ple
Neogene
Q
18 16 14 12 10 8 6 4 2 0
Mya
n = 233
n = 94
Net speciation rates at tips (event/Myr)
0.8
0.6
0.4
0.38
0.42
Eichlerangraecum
Conchograecum
Campylocentrum
Dendrophylax
Diaphananthe
Calyptrochilum
Eurychone
Aerangis
Kylicanthe
Angraecopsis
Sphyrarhynchus
Mystacidium
M.tanganyikense
Rhipidoglossum
Nephrangis
Dolabrifolia
Afropectinariella
Microcoelia
Bolusiella
Ancistrorhynchus
Solenangis
Rangaeris
Podangis
Planetangis
S.impraedicta
Cyrtorchis
Aziza
Ypsilopus
Plectrelminthus
Listrostachys
Summerhayesia
Tridactyle
Flower colour
Fully pigmented
White or partially white
14 12 10 8 6 4 2 0

Figure S4. Ancestral state estimation and state-dependent speciation rates of flower colour using hidden-state SSE models on the dated phylogeny of 327 species of angraecoid orchids

Pie charts at nodes represent ancestral states and their probabilities under a CID4 model. The character states of extant species are represented on the right side of the tree by coloured boxes. In addition, species for which a sphinx has been observed visiting or carrying pollinaria are indicated by a sphinx silhouette. Pli: Pliocene; Ple: Pleistocene; Q: Quaternary. The inset shows the net speciation rates at tips associated with each state with density plots; ‘n’ indicates the number of species per state; the median is indicated by a diamond with value below

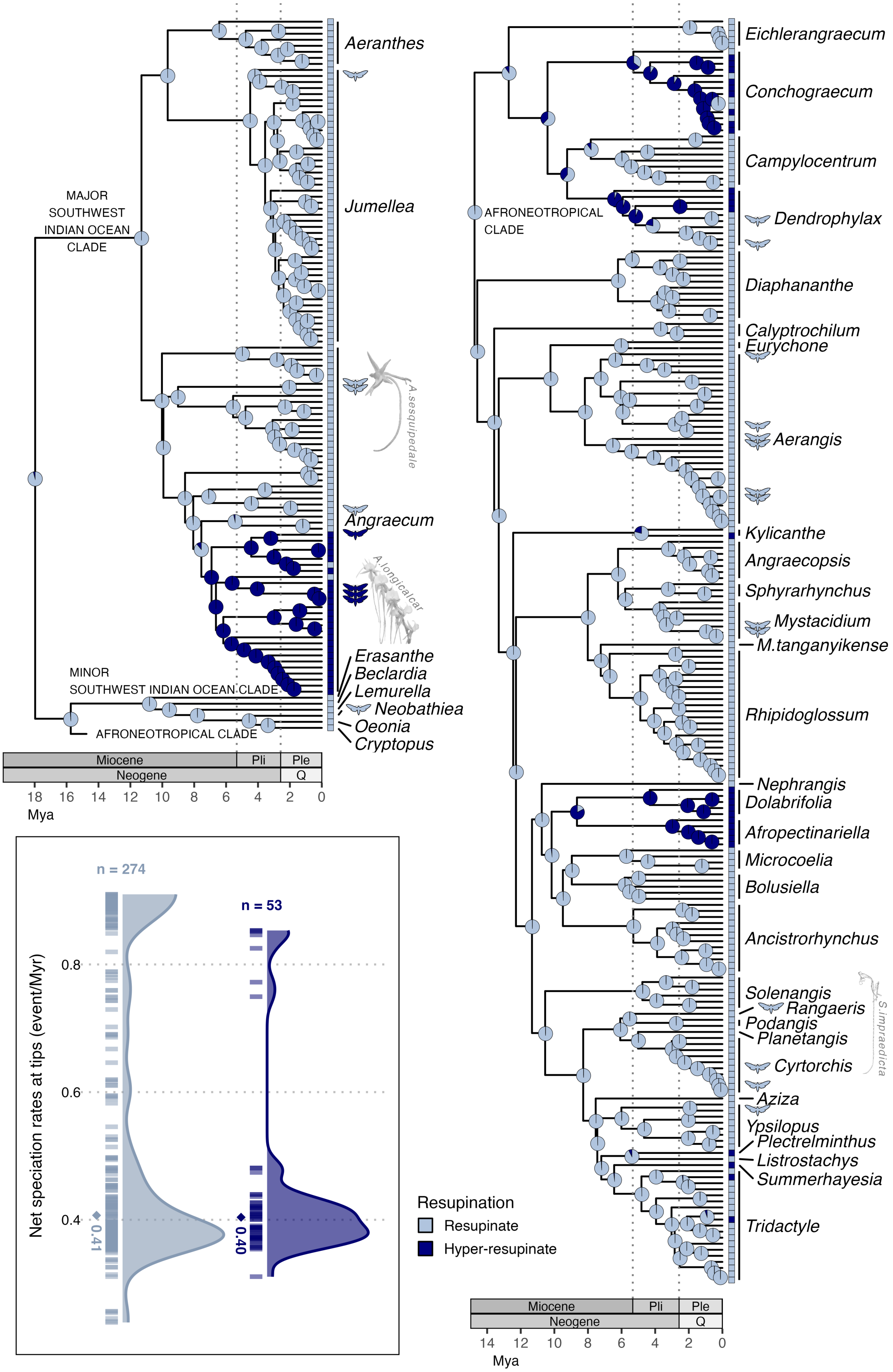
Aeranthes
Jumellea
MAJOR
SOUTHWEST
INDIAN OCEAN
CLADE
A.sesquipedale
Angraecum
A.longicalcar
Erasanthe
Beclardia
Lemurella
Neobathiea
Oeonia
Cryptopus
MINOR
SOUTHWEST INDIAN OCEAN CLADE
AFRONEOTROPICAL CLADE
Miocene
Pli
Ple
Neogene
Q
18 16 14 12 10 8 6 4 2 0
Mya
n = 274
n = 53
Net speciation rates at tips (event/Myr)
0.8
0.6
0.4
0.41
0.40
Eichlerangraecum
Conchograecum
Campylocentrum
AFRONEOTROPICAL
CLADE
Dendrophylax
Diaphananthe
Calyptrochilum
Eurychone
Aerangis
Kylicanthe
Angraecopsis
Sphyrarhynchus
Mystacidium
M.tanganyikense
Rhipidoglossum
Nephrangis
Dolabrifolia
Afropectinariella
Microcoelia
Bolusiella
Ancistrorhynchus
Solenangis
Rangaeris
Podangis
Planetangis
S.impraedicta
Cyrtorchis
Aziza
Ypsilopus
Plectrelminthus
Listrostachys
Summerhayesia
Tridactyle
Resupination
Resupinate
Hyper-resupinate
Miocene
Pli
Ple
Neogene
Q
14 12 10 8 6 4 2 0
Mya

Figure S5. Ancestral state estimation and state-dependent speciation rates of flower resupination using hidden-state SSE models on the dated phylogeny of 327 species of angraecoid orchids

Pie charts at nodes represent ancestral states and their probabilities under a CID4 model. The character states of extant species are represented on the right side of the tree by coloured boxes. In addition, species for which a sphinx has been observed visiting or carrying pollinaria are indicated by a sphinx silhouette. Pli: Pliocene; Ple: Pleistocene; Q: Quaternary. The inset shows the net speciation rates at tips associated with each state with density plots; ‘n’ indicates the number of species per state; the median is indicated by a diamond with value below.

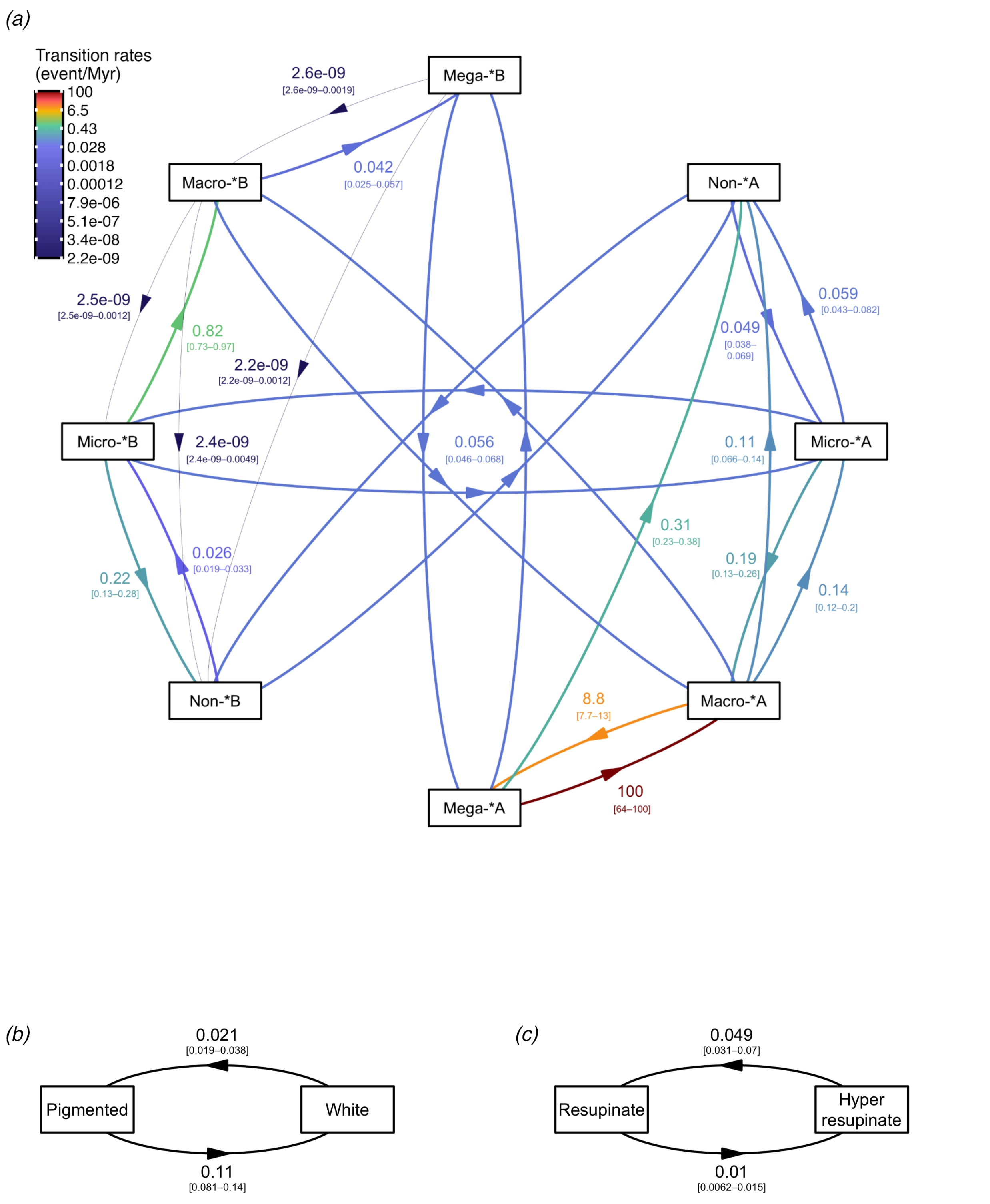

Figure S6. Transition rates of best-fitting SSE models for (a) floral syndrome, (b) flower colour and (c) resupination in angraecoid orchids

*: sphingophilous; confidence intervals are indicated in brackets; rate categories are indicated by capital letters A and B. Note that one rate hit the upper bound, set to 100.

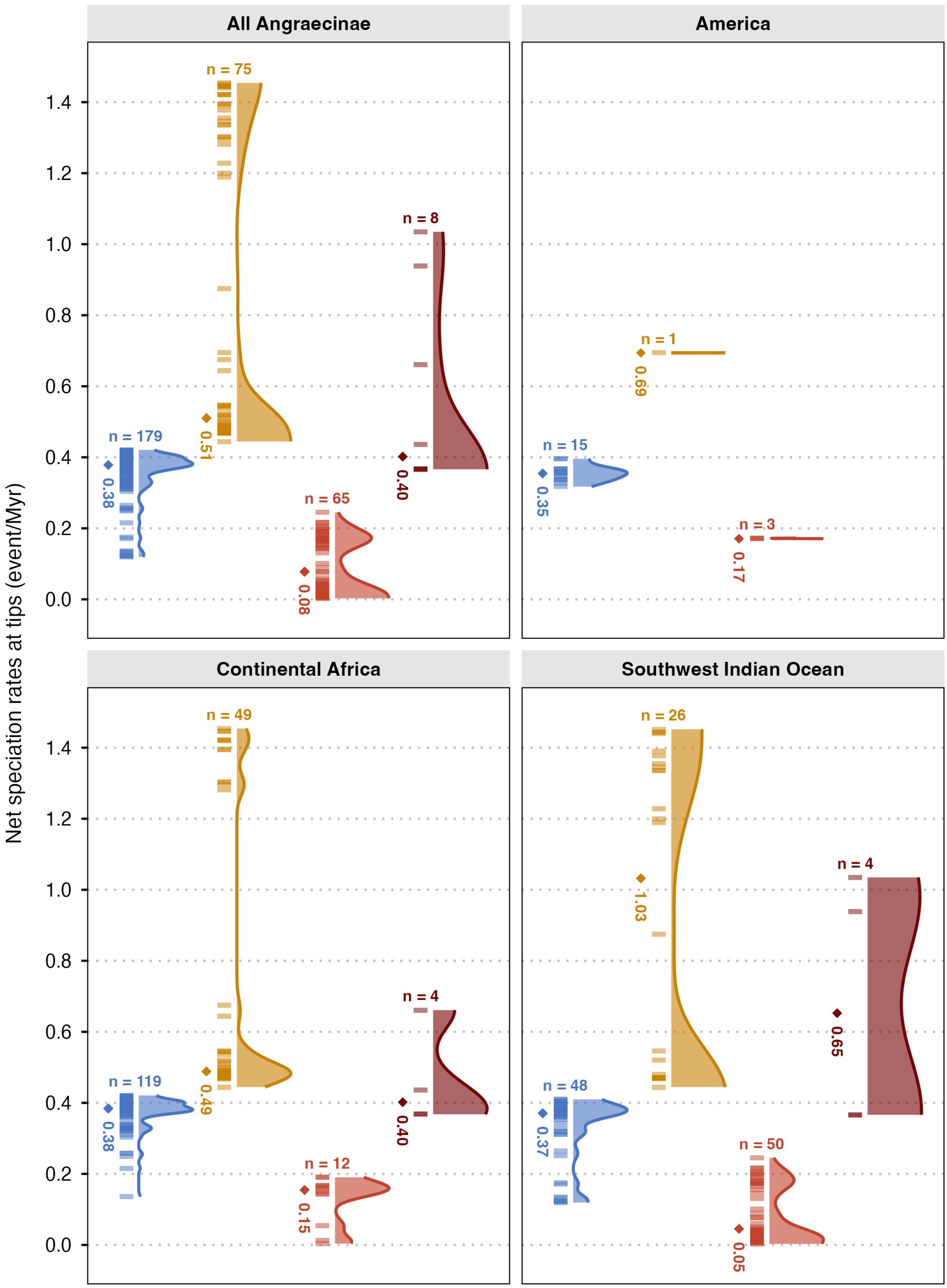
Net speciation rates at tips (event/Myr)
All Angraecinae
America
Continental Africa
Southwest Indian Ocean
0.0
0.2
0.4
0.6
0.8
1.0
1.2
1.4
n = 179
n = 75
n = 65
n = 8
0.38
0.51
0.08
0.40
n = 15
n = 1
n = 3
0.35
0.69
0.17
n = 119
n = 49
n = 12
n = 4
0.38
0.49
0.15
0.40
n = 48
n = 26
n = 50
n = 4
0.37
1.03
0.05
0.65

Figure S7. Net state-dependent speciation rates of floral syndromes in 327 angraecoid orchids and in three geographic areas

‘n’ indicates the number of species per state; the median is indicated by a diamond with value below. Please refer to figure 2 for signification of colours.

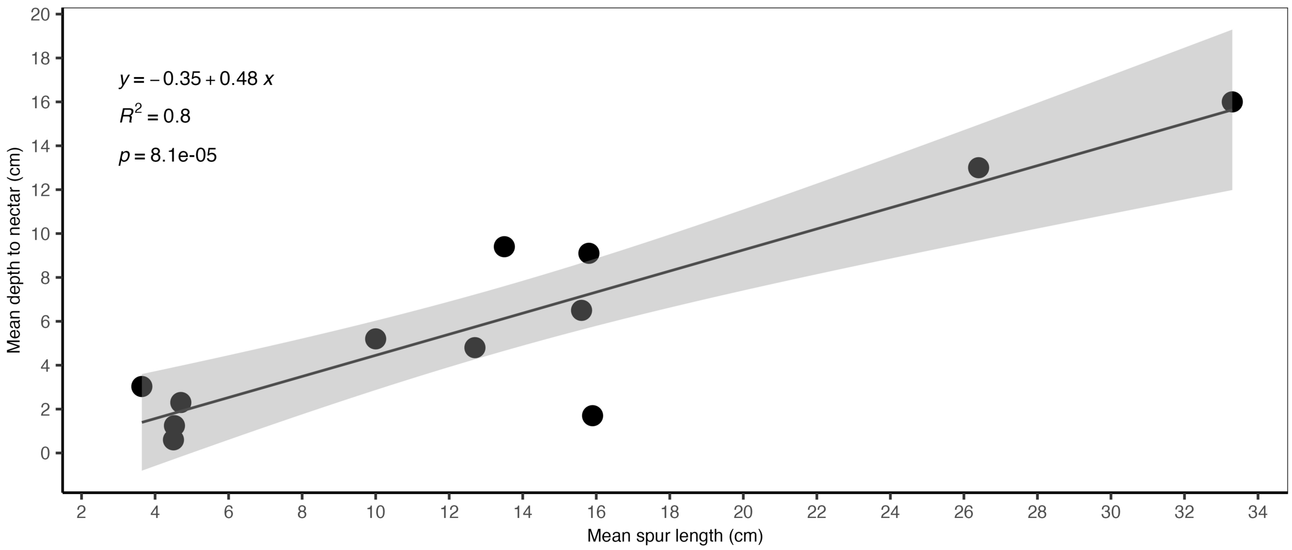

Figure S8. Linear regression (black line) with confidence interval (in grey) between mean hawkmoth proboscis and mean depth to nectar in 12 pollination case studies
